# Tonic dopamine and biases in value learning linked through a biologically inspired reinforcement learning model

**DOI:** 10.1101/2023.11.10.566580

**Authors:** Sandra Romero Pinto, Naoshige Uchida

**Author notes:** **Corresponding author** Correspondence to Naoshige Uchida and Sandra Romero Pinto.

## Abstract

A hallmark of various psychiatric disorders is biased future predictions. Here we examined the mechanisms for biased value learning using reinforcement learning models incorporating recent findings on synaptic plasticity and opponent circuit mechanisms in the basal ganglia. We show that variations in tonic dopamine can alter the balance between learning from positive and negative reward prediction errors, leading to biased value predictions. This bias arises from the sigmoidal shapes of the dose-occupancy curves and distinct affinities of D1- and D2-type dopamine receptors: changes in tonic dopamine differentially alters the slope of the dose-occupancy curves of these receptors, thus sensitivities, at baseline dopamine concentrations. We show that this mechanism can explain biased value learning in both mice and humans and may also contribute to symptoms observed in psychiatric disorders. Our model provides a foundation for understanding the basal ganglia circuit and underscores the significance of tonic dopamine in modulating learning processes.

## Introduction

Our ability to predict the outcomes of our actions is crucial in selecting and motivating appropriate actions. Systematic biases in future predictions or expectations, however, can lead to maladaptive behaviors, such as those observed in patients with various psychiatric disorders^1–4^. For example, overly negative or pessimistic predictions can contribute to major depression^1,5^, whereas excessively positive or optimistic predictions may be associated with pathological gambling, addiction, and mania^3,4,6–8^. Despite the importance of understanding the causes of biased future predictions, the biological mechanisms underlying them remain poorly understood.

Our future expectations and decisions are shaped by experiences of positive and negative events. The process of learning from outcomes has been modeled using reinforcement learning (RL) models^9–12^, where value predictions are updated based on reward prediction errors (RPEs), that is the discrepancy between received and expected outcomes. In addition to its role in learning, recent studies have indicated the importance of RPEs in mood; these studies have suggested that mood depends not on the absolute goodness of outcomes, but rather on the recent history of RPEs^13,14^.

In the brain, dopamine is thought to be a key regulator in this process of learning from positive and negative outcomes. The dynamics of dopamine are often categorized into two modes: tonic and phasic. Tonic dopamine refers to “baseline” dopamine that operates on a long timescale such as tens of seconds or minutes, while phasic activity refers to transient changes that occur at a much shorter, sub-second timescale, often triggered by external stimuli^15–18^. A significant body of evidence has shown that phasic responses of dopamine neurons convey RPEs and drive learning of values and actions^17–20^. On the other hand, changes in tonic dopamine might also modulate value learning, yet whether and how the level of tonic dopamine modulates learning remain poorly understood.

Previous studies have reported that patients with psychiatric disorders exhibit biased learning from positive versus negative outcomes. For one, some studies have shown that patients with major depression have a reduced sensitivity in learning from rewarding events, while their ability to learn from negative events remains relatively intact^1,5,21^. Similarly, patients with Parkinson’s disease are better at learning from negative than positive outcomes^22,23^. Analysis of these patients using RL models has suggested that biases in learning can be explained by alterations in specific parameters in RL models, such as the learning rate parameters or the sensitivity to positive and negative outcomes. For example, some studies have suggested that anhedonia in major depressive disorder may correspond to a reduced learning rate from positive compared to negative outcomes^1^.

Mechanistically, some of these changes in RL parameters can be linked to altered functions of dopamine. First, it has been shown that dopamine synthesis capacity, an approximate indicator of baseline dopamine levels, in the striatum, as measured using positron emission tomography (PET), correlates with learning rate parameters^24^. Second, dopamine medications can change the balance between learning from positive and negative outcomes^22,24,25^. Third, responses to positive outcomes in the nucleus accumbens (NAc), as measured based on blood oxygenation-dependent (BOLD) signals, are reduced in patients with psychiatric disorders such as depression^26–29^. These observations point to important roles of reinforcement learning processes and dopamine in regulating value learning. However, the parameters in RL models remain an abstract entity, and biological processes underlying changes in these parameters are still largely unknown.

One limitation in most RL models used in previous studies is that they do not reflect key neural circuit architectures in the brain (but see ^30–32^) nor recent findings on intracellular signaling and plasticity rules that can constrain how dopamine functions in biological circuits^33–35^. Incorporating these key biological factors may lead to better understanding of how changes in RL parameters may arise in psychiatric disorders. Furthermore, recent studies have found that the activity of dopamine neurons is consistent with a novel RL algorithm called distributional RL^36–38^. Distributional RL takes into account the diversity in dopamine signals, and a population of dopamine neurons together encodes the entire distribution of rewards, not just the average. Although distributional RL has shown to be efficient in solving various RL problems in artificial intelligence^37,39^, how distributional RL can be implemented in biological neural circuits and how distributional RL relates to biased value learning remain to be examined.

In this study, we sought to identify potential mechanisms that cause biased value predictions using biologically inspired RL models. To this goal, we first construct an RL model that incorporates recent biological findings, such as intracellular signaling and synaptic plasticity rules as well as the basic circuit architecture in the brain^32^. Based on this model, we propose two potential biological mechanisms that can cause optimistic or pessimistic biases in value predictions. We will then show that some existing data can be explained by one of these models. Finally, we will show how our model can provide an account of how biases in value predictions arise in psychiatric disorders.

## Results

### Basic reinforcement learning algorithms

Here we first formulate basic RL algorithms that will become the basis of our later models. In RL, an agent learns to predict the expectation of future rewards associated with a given state, a quantity termed as *value*^11^. For simplicity, we will drop the dependency on time here, but note that the basic results hold even if time is considered (Methods 1.1). Learning of value is driven by RPEs (*δ*), the discrepancy between the actual and expected reward (*r* and *V*, respectively) (Eq1). To improve the accuracy of the value prediction, RPEs are utilized to update the estimate of *V*. This is done iteratively by adding a fraction (α) of *δ* (Eq2) where *α* defines the learning rate.

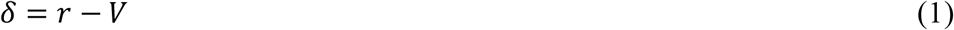

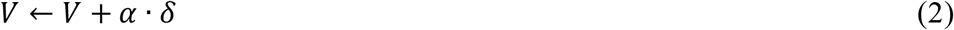

When the magnitude of reward *r* is fixed (i.e., deterministic environment), the value *V* learned through this algorithm (Eq 1 and 2) converges on *r* and the RPE converges on zero. When the magnitude of reward *r* varies stochastically trial-to-trial, the value at convergence fluctuates around the expected value of the reward distribution (see Methods 1) (Fig. 1a) and the RPE around zero.

**Figure 1.**
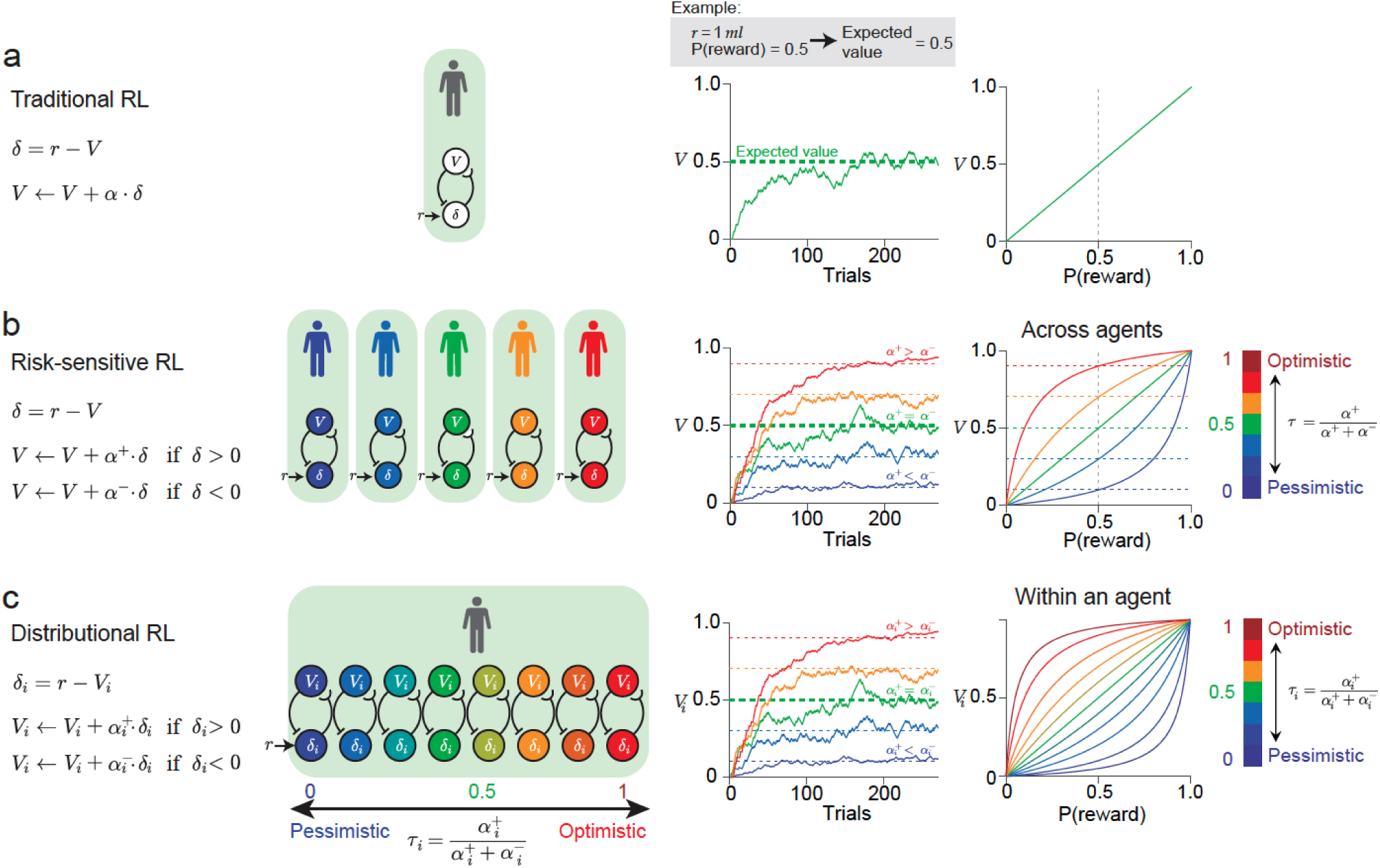
Reinforcement learning models. **a**. Traditional reinforcement learning with a single learning rate (*α*) for both positive and negative RPEs (*δ*) for the value updates (left). This update rule makes value estimate (*V*) converge on the expected value of the reward distribution (middle). When the reward probability is varied (i.e., for Bernoulli distributions), the *V* at convergence scales linearly with the reward probability (right). **b**. Risk-sensitive reinforcement learning with different learning rates (*α*^+^, *α*^−^) for positive and negative RPEs, respectively (left). This update rule makes value estimate (*V*) converge on the quantities that are higher or lower than the expected value of the reward distribution (middle). As the reward probabilities are varied, the convexity of the convergent value *V* changes depending on the asymmetry between *α*^+^ and *α*^−^ (Methods 1.3.3). The level of the bias is determined by the asymmetric learning rate parameter *τ* (right). **c**. Distributional reinforcement learning contains a set of value predictors (*V*_*i*_) each with a given learning rate for positive and negative RPEs (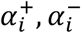, respectively) (left). This makes each value predictor converge on the quantity equal to the *τ*_*i*_-th expectile of the reward distribution. Thus, each value *V*_*i*_ represents an expectile, and together the set of *V*_*i*_ represents the entire distribution (Methods 1.2) (right).

#### Risk-sensitive RL

In the framework called risk-sensitive RL^40^, learning rates are defined separately for positive and negative RPEs (denoted by *α*^+^, *α*^−^).

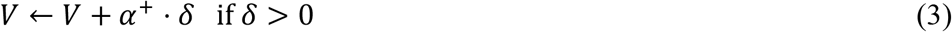

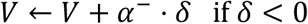

In the presence of stochastic rewards, when the learning rates between positive and negative RPEs are different, the value learned through this algorithm (Eq 1 and (3) does not converge on the expected value, but instead on a value higher or lower than the expected value depending on the relative amplitude of the learning rates *α*^+^, *α*^−^. This algorithm, therefore, develops optimistic or pessimistic value expectations, respectively. This learning algorithm is called “risk-sensitive” because values of probabilistic (risky) rewards are biased compared to deterministic (certain) rewards, and, therefore, the agent develops a preference between risky and certain rewards even when the expected values are the same (Fig. 1b).

#### Distributional RL

The concept of asymmetric updates has been utilized in a novel RL framework called distributional RL^36,37,41^. This algorithm allows an agent to learn the entire probability distribution of rewards, instead of the expected value which is typically the learning target in traditional RL algorithms (Fig. 1c). In distributional RL, an agent is equipped with a set of value predictors (*V*_*i*_), where *i* corresponds to the index of the value predictor (or “value neuron”). The value of the *i*-th neuron (*V*_*i*_) is updated based on the learning rates 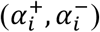 and the RPE (*δ*_*i*_) for that neuron *i*:

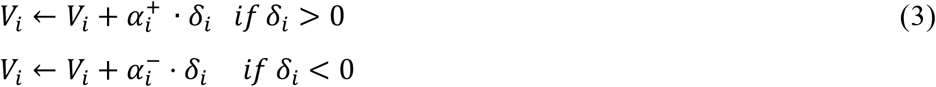

Similar to risk-sensitive RL, the learned value of each value predictor converges on estimates larger or lower than the expected value, determined by the ratio between 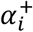 and 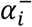.

Mathematically, each *V*_*i*_ converges on the *τ*_*i*_-th expectile of the distribution (Fig. 1c) where *τ*_*i*_ (asymmetric scaling factor) is defined by:

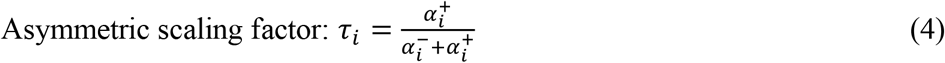

Expectiles are the solutions to asymmetric least squares minimization and generalize the mean of a distribution (with the mean being the 0.5^th^ expectile) as quantiles generalize the median (with the median being the 0.5^th^ quantile)^42^. Since a set of expectiles can define a distribution, the diversity of *τ*_*i*_ across the population enables learning of the entire probability distribution.

#### Problem

In both risk-sensitive RL and distributional RL, unbalance in learning rate parameters for positive and negative RPEs gives rise to optimistic and pessimistic biases in learned values. Importantly, however, the underlying biological mechanism regulating learning rate parameters (*α*^+^, *α*^−^) and asymmetry thereof (*τ*) remains unclear.

In the following sections, we will discuss potential biological mechanisms that regulate asymmetric learning rates (*α*^+^, *α*^−^). We will first modify the above RL algorithms to incorporate important neural circuit architectures in the brain. We will then propose two key biological mechanisms that can give rise to asymmetric learning rates (called Model 1 and 2). We will then show that our model can explain previous experimental data and psychiatric conditions.

### Incorporating biological features into RL models

The above RL models provide algorithmic-level formulations, yet they do not recapitulate fundamental characteristics of the neural circuits thought to perform RL in the brain^43–46^. We next incorporate some of the important circuit and synaptic properties into the model.

In the brain, it is thought that dopamine neurons in the ventral tegmental area (VTA) broadcast RPEs^17^ and modulate synaptic plasticity in dopamine-recipient areas^33,47^. The striatum is the major target of dopaminergic projections. It has been thought that spiny projection neurons (SPNs) in the striatum represent values, and dopamine modulates plasticity of synapses on SPNs^33,34,47,48^ (Fig. 2a). Under this framework, the value representations in SPNs are updated by dopaminergic RPEs. In most RL models, each value predictor is typically updated by both positive and negative RPEs. If the value is computed based on a weighted sum of some inputs (i.e., using linear function approximation^11^), the update rules described above (Eq 3 and 4) are equivalent to performing a semi-gradient descent that minimizes RPEs^11^ (see Methods).

**Figure 2.**
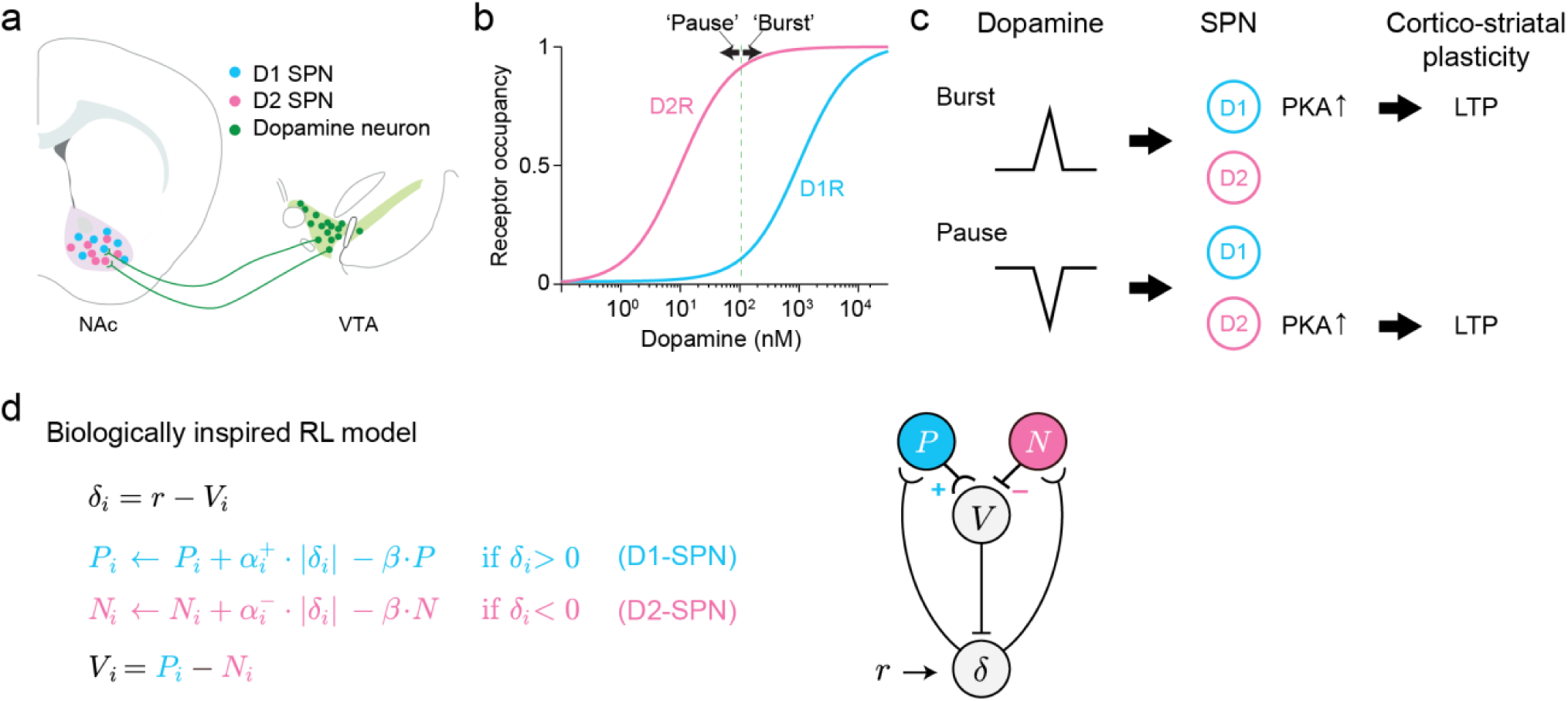
Biologically inspired reinforcement learning model. **a**. Schematic of the basal ganglia circuitry. Dopaminergic neurons in the VTA modulate plasticity at the level of the cortico-striatal synapses on SPNs in the NAc. The SPNs are subdivided depending on the dopamine receptor type they express (D1R or D2R). **b**. Dose-occupancy curves for the D1R and D2R describing receptor occupancies as a function of dopamine concentrations. The curves are shifted between each other due to the different affinities of the receptors. The arrows represent 3-fold increase (“burst”) and decrease (“pause)” in dopamine concentrations, which causes left-ward or right-ward shifts of the same magnitudes in the log-scale. **c**. Schematic of the plasticity rules of VTA-NAc circuitry^33–35^. Transient increases in dopamine, caused by bursts in firing rate of dopamine neurons, generates increases in PKA activity in D1R-expressing SPNs, leading to LTP in the cortico-striatal synapses. Transient decreases in dopamine, caused by pauses in firing rate of dopamine neurons, generates increases in PKA activity in D2R-expressing SPNs, leading to LTP in the cortico-striatal synapses. **d**. Schematic and equations of biologically inspired reinforcement learning model^32^ VTA, ventral tegmental area; NAc, nucleus accumbens; SPN, spiny projection neurons; D1R, D1-type dopamine receptor; D2R, D2-type dopamine receptor; PKA, protein kinase A; LTP, long-term potentiation.

The basic architectural assumptions of these RL models are, however, at odds with the RL circuitry in the brain. Importantly, in the striatum, there are two major classes of dopamine-recipient SPNs characterized based on the type of dopamine receptor that they express: D1- or D2-type dopamine receptors (D1R and D2R)^48^. SPNs expressing D1R and D2R constitute the so-called direct and indirect pathways and exert opposing effects on downstream “output” neurons, with each pathway promoting or opposing a certain output (e.g., movement).

In addition to the presence of direct and indirect pathways, there are two additional properties in these opposing populations that need to be considered^32^. First, D1R and D2R have different affinities to dopamine: high in D2R and low in D1R (EC_50_ affinity constant is 1 µM for D1R and 10 nM for D2R)^49,50^. The dose-occupancy relationship of D1R and D2R are sigmoidal but they are shifted with one another with respect to dopamine concentration (Fig. 2b). Importantly, at normal dopamine levels (approx. 50-100nM)^51,52^, D2Rs are mostly occupied while D1Rs are mostly unoccupied (Fig. 2b). Although whether the affinities of D1R and D2R differ at the molecular level has been questioned^53^, a recent study showed that intracellular signaling through protein kinase A (PKA) in D1- and D2-SPNs is triggered by a phasic increase and a decrease in dopamine, respectively, in behaving animals^35^. These results are consistent with (apparent) difference in affinities of D1R and D2R assumed in previous studies^49^, although the exact reason for the difference remains to be clarified^53^.

The second important property pertains to different learning rules in D1- and D2-SPNs which are predicted from different affinities of the receptors. Consistent with the observed PKA signals in these cells, recent studies have shown that glutamatergic inputs on D1-SPNs are potentiated by a transient *increase* in dopamine, whereas those on D2-SPNs are potentiated by a transient *decrease* in dopamine^34,35^ (Fig. 2c), supporting opposing plasticity rules between D1- and D2-SPNs.

There have been previous efforts to incorporate in RL models the direct and indirect pathways (also called “Go” and “NoGo” pathways, respectively) such as Opponent Actor Learning (OpAL^30^, OpAL*^54^) and Actor learning Uncertainty (AU)^32^ models. These previous models were developed as *Actor-Critic models*^11^. Here, we will build on the AU model to focus on the problem of value learning and extend it to support risk-sensitive RL and distributional RL. Our model has two separate populations of value predictors corresponding to D1R- and D2R-SPNs, that store the quantities P*i* and *N*_*i*_ respectively (Eq 6, Fig. 2d). Mimicking dopamine’s effect on potentiation, *P*_*i*_ or *N*_*i*_ will increase their estimates if an RPE is positive or negative, respectively, with the learning rates defined by 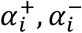(Eq. 6). Importantly, the value *V*_*i*_ can be obtained simply by taking the difference between *P*_*i*_ and *N*_*i*_. (Eq. 7).

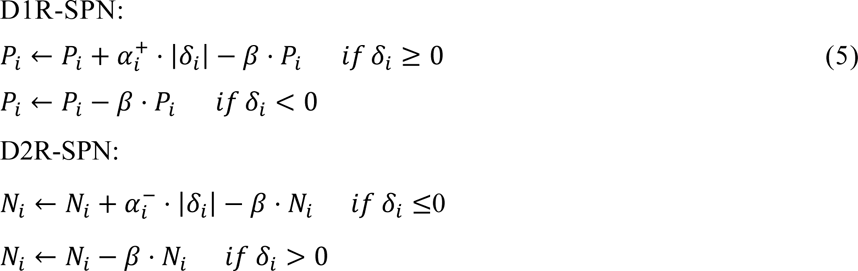

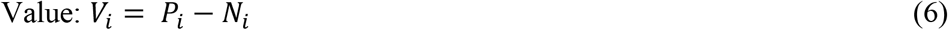

where *β* is a decay parameter which represents synaptic decay in the absence of RPEs. This model (Eq 6 and 7) preserves various essential properties of the previous RL models: (1) learning in *P* and *N* can be combined to provide a simple update rule for value *V*, and (2) this update rule approximates the gradient descent that minimizes RPEs (when *β* = 0, the update rule is equivalent to the gradient descent). Importantly, with *β* > 0, we can show that these simple learning rules guarantee convergence of value, without the need for additional mechanisms to modulate the learning rates over iterations (Methods 1.3).

For instance, in a stochastic environment where there is a probability *p* of receiving a reward of a fixed magnitude *r* = 1, the stochastic fixed point of the learned value *V*_*i*_ (i.e., convergence point) will be defined by Eq 7.

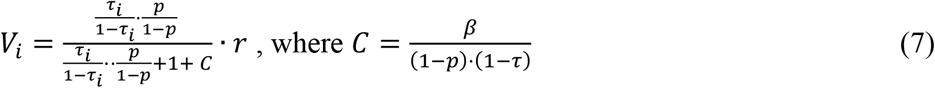

Note that Eq.7 contains an additional term *C* which depends on *β* and this decay factor *β* is important to stabilize the *P*_*i*_ and *N*_*i*_ estimates (avoid infinite increases) (Methods, 1.3.1, Extended Data Fig. 1).

This formulation now provides a mechanistic model suitable for risk-sensitive RL (when there is one value predictor) as well as distributional RL (when there are multiple value predictors), which incorporate the neural circuit architecture and plasticity rules of D1R- and D2R-SNPs found in the brain.

With this model at hand, we now discuss potential mechanisms that produce an asymmetry in learning rates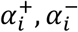, which, in turn, causes biases in value predictions. In principle, learning rate parameters can be a function of (1) the scaling of RPEs, i.e., the slope of dopamine responses as a function of RPE (δ), and (2) the scaling of value updates, i.e., the efficacy of dopamine-dependent synaptic plasticity at the level of SPNs. In the following, we discuss each scenario, emphasizing the role of either tonic or phasic dopamine activity in each of these mechanisms (Model 1 and 2, respectively). For simplicity, we will start with a model in which *α*^+^, *α*^−^ are equal for all neurons within both *P* and *N* populations, equivalent to risk-sensitive RL. We will then relax this assumption and introduce heterogeneity by allowing 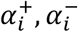 to vary across neurons, implementing a form of distributional RL.

#### Model 1: The role of baseline dopamine in asymmetric learning

As discussed above, D1R and D2R have different affinities to dopamine which leads to different levels of receptors’ occupancy at a given baseline dopamine level (Fig. 2b). Crucially, due to the sigmoidal shape of the dose-occupancy curves, the slope of the curve changes with baseline dopamine level, which means that a given dopamine transient leads to a different change in receptor occupancy depending on the baseline dopamine level (Fig. 3a,b). That is, the receptors’ *sensitivity* changes with baseline dopamine (Fig. 3c). In addition, a key consequence of the distinct receptors’ affinities is that an increase and decrease in baseline dopamine will cause opposite changes in the sensitivity of D1R and D2R. Specifically, an increase in dopamine will decrease D1R sensitivity relative to D2R, whereas a decrease in dopamine will increase D2R sensitivity relative to D1R (Fig. 3c,d).

**Figure 3.**
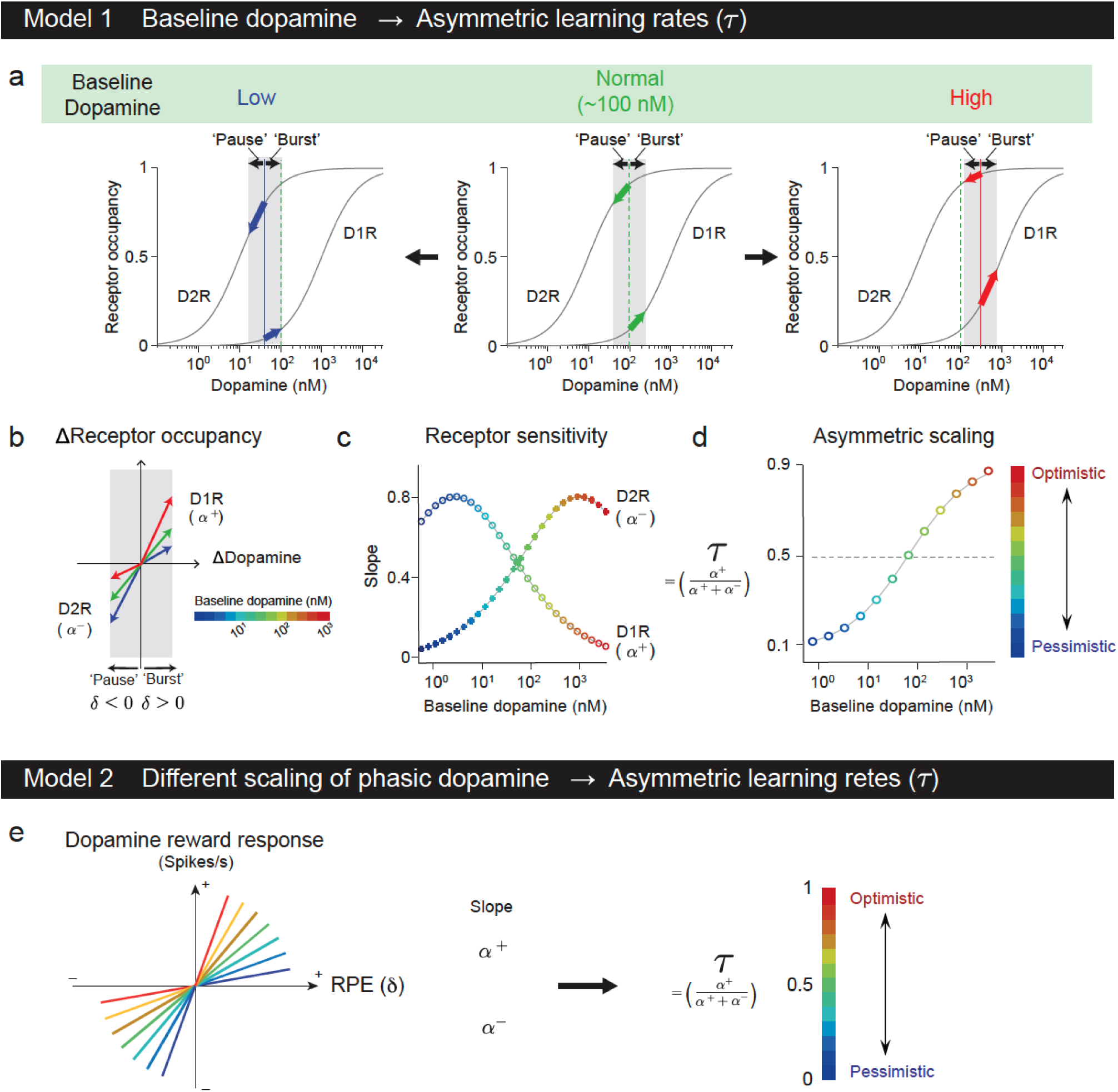
Potential mechanisms for asymmetric learning. **a**. Schematic of the mechanism by which increases or decreases in baseline dopamine modulates the degree to which bursts and pauses in dopamine causes changes in D1R and D2R occupancy. Increases in baseline dopamine makes dopamine pauses to cause greater decreases in D2R occupancy than the increases in D1R occupancy caused by dopamine bursts. Conversely, decreases in dopamine, makes dopamine bursts to cause smaller increases in D1R occupancy than the decreases in D2R occupancy caused by dopamine pauses. **b**. Schematic of the change in receptor occupancies in D1R and D2R, for a given transient increase (‘burst’) or decrease (‘pause’) in dopamine, receptively. A pause and a burst in dopamine correspond to *δ* < 0 and *δ* > 0 in the model. The slope is modulated by the baseline dopamine (colormap) and corresponds to the receptor’s sensitivity to dopamine transients. **c**. Receptor sensitivity for D1R and D2R as a function of baseline dopamine. In Model 1, we assume that the receptor sensitivity acts as a scaling factor on the PKA activity induced by burst and pauses. That is, PKA_D1_ ∝ *α*^+^ · *δ* · **1**_*δ*>0_ and PKA_D2_ ∝ *α*^−^ · *δ* · **1**_*δ*<0_. **d**. Asymmetric scaling factor (*τ*) as a function of baseline dopamine. Colors depict how ‘optimistic’ or ‘pessimistic’ the convergent value estimate will be when learning with a given *τ*. **e**.Model 2. Left, the relationship between dopamine reward responses (spikes/s) and RPEs. The slopes of these response functions correspond to the asymmetric learning rates (*α*^+^, *α*^−^) for positive and negative RPEs, respectively. Colors depict how ‘optimistic’ or ‘pessimistic’ the convergent value estimate will be when learning with a given asymmetric scaling factor.

Building on this insight in Model 1, we postulate that the learning rates for positive and negative RPEs are a function of the D1R and D2R sensitivity, respectively. This is supported by previous studies that have reported that the effect of dopamine transients of a given magnitude in SPNs’ plasticity can be modulated by the level of dopamine baseline^34^. In addition, it has been reported that the level of potentiation in SPNs^33,34^ or plasticity, which are related to intracellular signals^35^, scale with the magnitude of dopamine transients, keeping all else fixed. These observations can be summarized with the following rule:

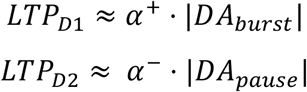

Where *α*^+^, *α*^−^ correspond to the receptors’ sensitivities and depend on the dopamine baseline level. This rule can be directly related to the update equations for the *P* and *N* populations in our model:

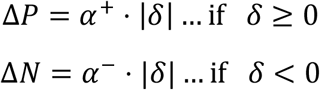

It can been shown that this learning rule is in agreement with the normative solution for the problem of value learning^11^ (Methods 1.6).

In short, in Model 1, a shift in the baseline dopamine level causes asymmetries in scaling of the value updates for positive versus negatives RPEs via the modulation of receptors’ sensitivities, which leads to value learning biases. This is a direct consequence of the dose occupancy relationships of D1R and D2R (Fig. 3b-d).

#### Model 2: Asymmetric scaling of phasic dopamine responses, inspired by distributional RL

In Model 2, we postulate that the learning rates *α*^+^ and *α*^−^ are a function of the scaling (i.e., ‘slope’) of dopamine responses evoked by positive and negative RPEs, respectively:

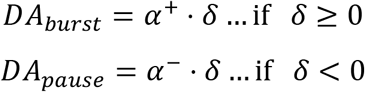

This is supported by a previous study on distributional RL that demonstrated that individual dopamine neurons vary in terms of how the magnitude of reward responses is scaled as a function of positive and negative RPEs (Fig. 3e)^36^.

In the distributional RL framework, individual dopamine neurons vary in terms of their asymmetric scaling factor *τ*_*i*_ and each of the multiple value predictors (*V*_*i*_) converges on the *τ*_*i*_-th expectile of the reward distribution (Eq. 4). However, in most applications of distributional RL, action selection is still based on the expected value of the reward distribution. Thus, the quantity relevant to action selection can be described using the population level average *τ*_*population*_, and biased value learning at the behavioral level could arise if *τ*_*population*_ is higher or lower than 0.5. This can occur from a differential loss of optimistic or pessimistic dopamine neurons. Another possibility is an overall upward or downward shift in the distribution of *τ*_*i*_ across the population due to, for example, intrinsic factors modulating the gain of dopamine phasic responses.

Risk-sensitive RL can be thought of as a special case of distributional RL which has only one value predictor. Here, the slope of the average dopamine evoked transient to positive and negative RPEs, may correspond to the population level learning rates for positive and negative RPEs (*α*^+^, *α*^−^), respectively. If the asymmetric scaling factor *τ* is higher or lower than 0.5, value learning will be biased (Fig. 3e).

### Testing for evidence of either model in experimental data

#### Tian and Uchida (2015)

We next examined whether Model 1 or 2 can explain empirical data obtained in experimental animals or humans. We first examined the data obtained in mice in our previous study^55^. In this study, the authors tested the effect of lesioning the habenula, a brain structure implicated in depression^56–58^, on the activity of dopamine neurons and on reward-seeking behavior. Head-fixed mice were trained in a Pavlovian conditioning task in which odor cues predicted reward with different probabilities (10%, 50%, 90%). After performing habenula (n=5) or sham (n=7) lesions (Fig. 4a), the spiking activity of VTA dopamine neurons was recorded while mice performed the task.

**Figure 4.**
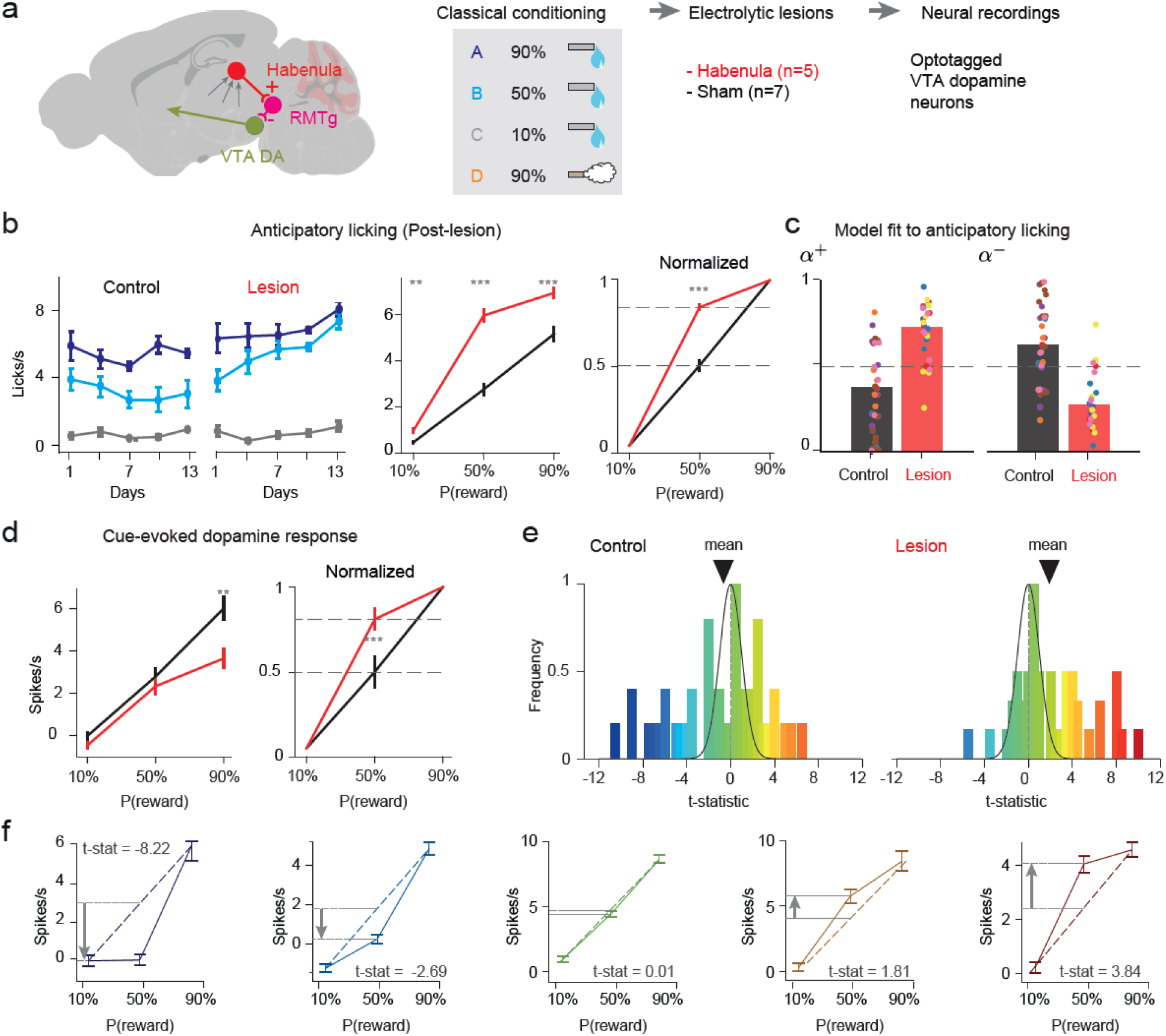
Habenula lesions leads to optimistic reward-seeking behavior and cue-evoked responses in dopamine neurons. **a**. Schematic of the experiment performed by Tian and Uchida (2015)^55^. Animals were trained in a classical conditioning task in which 3 odor cues predicted rewards of different probabilities (10%, 50%, 90%) and one odor cue predicted 80% probability of an air puff. Animals then underwent habenula (*n* = 5) or sham (*n* = 7) lesions and trained on the task again. The neural recordings were performed from optotagged VTA dopamine neurons once behavior stabilized. **b**. Anticipatory licking across sessions after lesions (left,). There was a significant increase in anticipatory licking to the 10% (U-statistic = –2.895, *P* = 0.003792, two-sided Mann-Whitney U-test), 50% (U-statistic = –5.579, *P* < 1 x 10^−9^, two-sided Mann-Whitney U-test) and 90% (U-statistic = –3.487, P =0.00048, two-sided Mann-Whitney U-test) cues (*n* = 31 for control *n* = 30 for lesion) that results from progressive changes across sessions. The anticipatory licking curves show a linear scaling with reward probability in the control group, and a convex curve for the lesion group (mean ± s.e.m across animals, U-statistic = –6.444, *P*< 1 x 10^−^, two-sided Mann-Whitney U-test for the 50% cue normalized response). These curves are predicted by RL agents with symmetric and asymmetric (*α*^+^ > *α*^−^) learning rates for the control and lesion groups, respectively, assuming a linear mapping between anticipatory licking and value prediction. **c**. RL model fits to the anticipatory licking on a trial-by-trial basis using a risk-sensitive RL models that allows for separate learning rates of positive and negative RPEs. Each dot represents a session (*n* = 35 control, *n* = 30 lesion) and each color a mouse (*n* = 7 control, *n* = 5 lesion). The fits show a significant difference in the learning rates between control and lesion groups (U-statistic = –4.679, *P* < 1.0 x 10^−5^, pooling sessions across mice in each group). **d**. Cue-evoked dopamine responses from opto-tagged VTA dopamine neurons (mean ± s.e.m across neurons, *n* = 45 control group, *n* = 44 lesion group). There was a decrease in the absolute magnitude of responses to the 90% cue (U-statistic = 3.249, *P* = 0.0011, two-sided Mann-Whitney U-test) after habenula lesions (left). The normalized cue-evoked responses show the similar pattern as the normalized anticipatory-licking with a linear and convex function for the control and lesion groups, respectively, with a significant increase in normalized response to the 50% cue after lesions (U-statistic = –3.824, *P* = 0.000131, two-sided Mann-Whitney U-test) These curves are predicted by agents with symmetric and asymmetric learning rates for control and lesion groups, respectively. **e**. Distribution of t-statistics comparing the cue-evoked response to the linear interpolation point between the 90% and 10% cue-evoked responses for each dopamine neuron. The distribution of t-statistics for the control and lesion cases was wider than what is expected from random noise (Monte Carlo test for standard deviation different from zero: *P* = 0.0222 control, *P* = 0.0217 lesion, 1000 batches). The distribution was shifted to values larger than 0 in the lesion case (Monte Carlo test for mean larger than zero: *P* = 1 control, *P* = 0.022 lesion, 1000 batches) indicative of an optimistic bias in the distribution. The lesion group distribution was also significantly shifted to higher values with respect to the control group distribution (U-statistic = –2.815, *P* = 0.0024, single-sided Mann-Whitney U-test). Arrow heads: the mean of the *t*-statistics. **f**. Example of *t*-statistics calculations for dopamine neurons taken from the control group (mean ± s.e.m across trials). A *t*-statistic value close to 0 indicates linear scaling of cue-evoked responses with reward probability; a *t*-statistics value lower or greater than 0 indicates a concave or convex function of cue-evoked responses against reward probability, indicative of a pessimistic or an optimistic bias, respectively.

After lesions, mice exhibited an elevated reward-seeking behavior (anticipatory licking) in response to cues predictive of probabilistic rewards, consistent with an optimistic bias in reward expectation (Fig. 4b, right). Importantly, anticipatory licking gradually increased over several sessions after lesions, suggesting that the optimistic bias developed through learning (Fig. 4b, left). To bring insight into the underlying cause of these biases, we fit two different RL models to the anticipatory lick responses on a trial-by-trial basis (Extended Data Fig. 2), assuming a linear relationship between value predictions and anticipatory licking. These models considered either a change in the sensitivity to rewards (Extended Data Fig. 2b) or asymmetric learning rates (Extended Data Fig. 2c). This analysis showed that the biases observed in the behavior could be explained by asymmetric learning rates, but not by reward sensitivity because the reward sensitivity was unchanged in the lesion group with respect to the control group (Extended Data Fig. 2c).

Dopamine neurons’ responses to reward-predictive cues reflect the increases in value expectation predicted by the cue with respect to baseline. The overall magnitudes of cue-evoked responses were not elevated in lesioned animals compared to control animals (Fig. 4d). However, the shape of the response curve indicated an ‘optimistic’ bias: although in control animals, cue responses scaled linearly with the expected value (i.e., reward probability), the response function of the lesioned animals was convex. In other words, in control animals the response to the 50%-reward cue was not significantly different from the quantity that results from the linear interpolation between the responses to 10%- and 90%-reward cues. In lesioned animals, however, the response to the 50%-reward cue was significantly greater than this quantity, which is indicative of an optimistic bias in value predictions (Fig. 4d, see Methods 1.3.3 for analysis of value predictions curve convexity). Such a change was observed at the level of the population average. Further analysis using individual neurons showed that when calculating a single-cell level metric that compares the 50%-reward cue to the same linear interpolation point, there was a broad distribution in this metric below and above the interpolated point both in the control and lesion groups (Fig. 4e-f). The distribution was, however, shifted in its mean in the lesion group (Fig. 4e). These analyses indicated that both anticipatory licking and dopamine cue responses have an optimistic bias as characterized by an overvaluation of probabilistic rewards, without still pointing to the underlying mechanism.

#### Model 2 cannot explain the optimistic biases in behavior and cue-evoked dopamine responses after Hb lesions

In Model 2, an optimistic bias in reward expectation can arise if the average of the asymmetric scaling factor at the population level (*τ*_*population*_) becomes greater than 0.5 (Fig. 5a,b).

**Figure 5.**
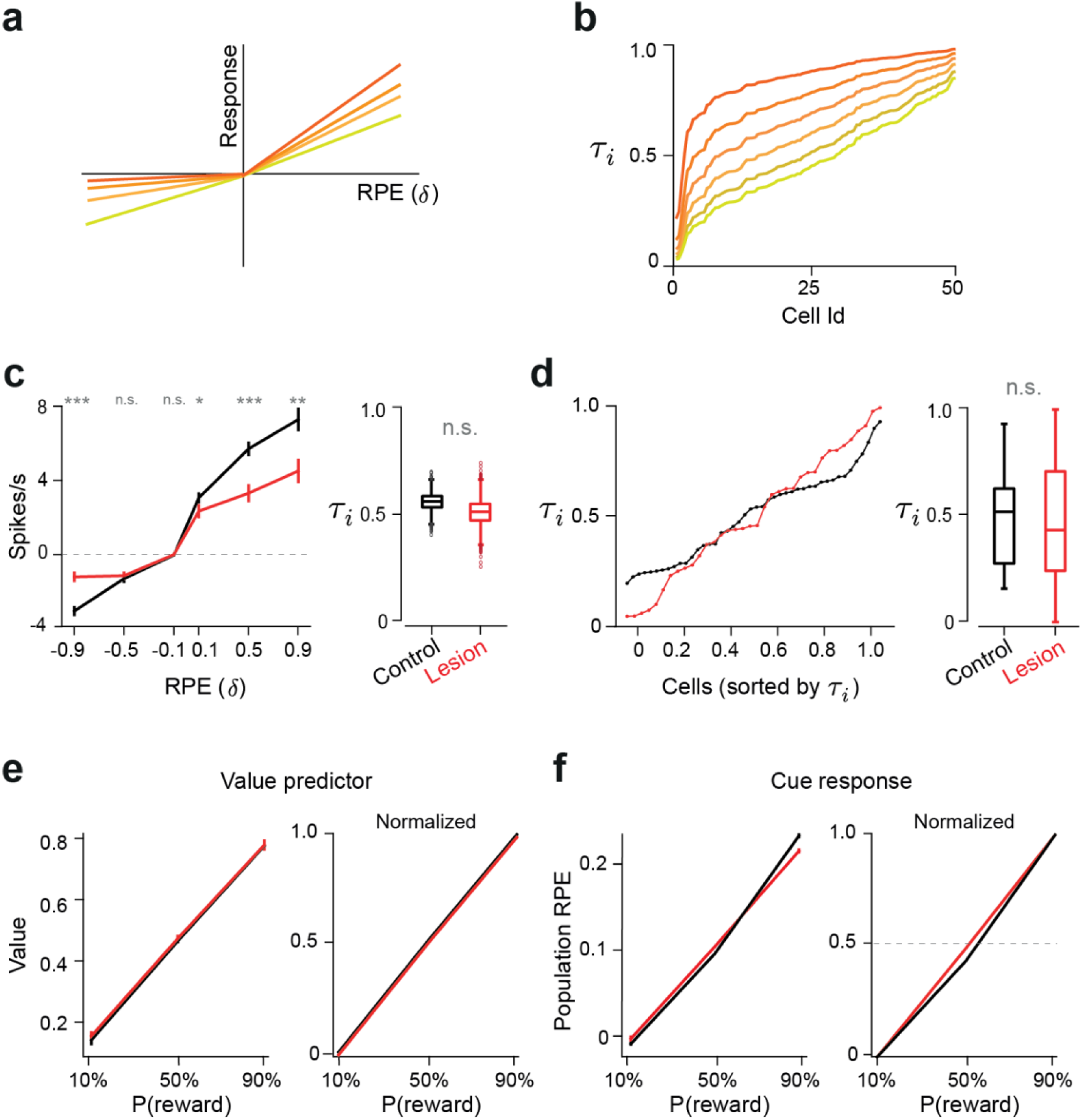
Model 2 cannot explain optimistic biases in behavior and cue-evoked dopamine responses of habenula lesioned animals. **a**. Possible changes in habenula lesion mice that could explain optimistic biases based on Model 2. At the level of the population dopamine responses, an optimistic bias can be caused by an increase in the slope of the average reward responses to positive RPEs and/or a decrease in the slope of the average reward responses to negative RPEs. **b**. At the level of the distribution of individual dopamine neuron responses, an optimistic bias can be caused by an overall increase in the mean of the distribution of asymmetric scaling factors (*τ*_*i*_), computed from each individual neuron response function. **c**. Observed reward responses as a function of RPEs, averaged across the population of dopamine neurons for the control and lesion groups (left, mean ± s.e.m across neurons, *n* = 45 control group, *n* = 44 lesion group). There was a significant decrease in the reward responses for the 50% cue (U-statistic = 3.726, *P* = 0.000195, two-sided Mann-Whitney U-test) and 90% cue (U-statistic = 2.987, *P* = 0.00281, two-sided Mann-Whitney U-test), and for the omission responses for the 90% cue (U-statistic = -4.940, *P* <10^−4^, two-sided Mann-Whitney U-test). Distribution of asymmetric scaling factors (*τ*), computed from the average response function over the recorded neurons for the control and lesion groups (right). The distributions are the result of bootstrapping by randomly sampling neurons in 5,000 iterations. The distribution of differences between the obtained asymmetric scaling factors (*τ*_*lesion*_ − *τ*_*control*_) was not significantly larger than zero (5^th^ percentile = –0.1605). **d**. Distribution of asymmetric scaling factors (*τ*_*i*_), computed from each individual neuron response function for the control and lesion groups. Each dot represents a single neuron (n = 45 control group, *n* = 44 lesion group), and the neurons were sorted by asymmetric scaling factors (*τ*_*i*_). The means were not significantly different (right) (*t*-statistic = 0.3277, *P* = 0.627, *t*-test). **e**. Value predictions based on a TD learning model trained using the assumptions of Model 2 and the asymmetric scaling factors derived from the data. The model did not show any optimistic bias in the value predictors of the model trained with the lesion-derived asymmetric scaling factors. **f**. TD errors at cue show no signs of an optimistic bias in the model trained with the lesion-derived asymmetric scaling factors. Centre of box plot shows the median; edges are 25th and 75th percentiles; and whiskers are the most extreme data points not considered as outliers.

To test this idea, we obtained the asymmetric scaling factors (*τ*_*i*_) from dopamine neurons based on their outcome responses: for each neuron, we constructed outcome response functions against the magnitude of RPEs (Fig. 5c, Extended Data Fig. 3a,b). The response functions were obtained based on (1) whether reward was delivered (positive RPEs) or not (negative RPEs), and on (2) the magnitude of the reward expectation given by the reward probabilities predicted by each cue (0.1, 0.5, 0.9) (Extended Data Fig. 3a,b). We then obtained the point at which the responses are more likely to be below or above baseline (i.e., ‘zero-crossing points’)^36^ (Extended Data Fig. 3c), and computed 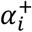 and 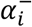 as the slopes of the responses in the positive and negative domains with respect to this zero-crossing point (Extended Data Fig. 3d), respectively. In both control and lesioned animals, asymmetric scaling factors tiled a wide range between 0 and 1 and presented other signatures consistent with distributional RL^36^ (Extended Data Fig.4). Nonetheless, although the variance of the distribution of asymmetric scaling factors was greater in lesioned animals, the mean did not change, indicating a lack of bias between 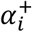and 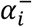at the population level (Fig. 5d). This was also the case when the asymmetric scaling factor was derived directly from the population average response (Fig. 5c). Thus, contrary to the conclusion in our previous study^15^, these analyses indicated that changes in reward responses (and the resulting scaling factor *τ*) do not explain the optimistic biases in behavior nor cue responses in lesioned animals (Fig. 5e,f).

#### Model 1 can explain the optimistic biases in behavior and cue-evoked dopamine responses by Hb lesion

In addition to changes in the magnitude of dopamine RPEs, we observed that the baseline firing rates of dopamine neurons were elevated in lesioned animals (Fig. 6a). According to Model 1, if these changes are followed by an increase in the baseline dopamine levels in the striatum, this should give rise to biased value learning (*α*^+^ > *α*^−^) and an optimistic bias in value expectation. In this way, this change in baseline firing can explain optimistic biases observed in lesioned animals. However, it remains unclear whether the observed change in baseline firing can result in functionally relevant levels of changes in the receptor occupancies discussed above.

**Figure 6.**
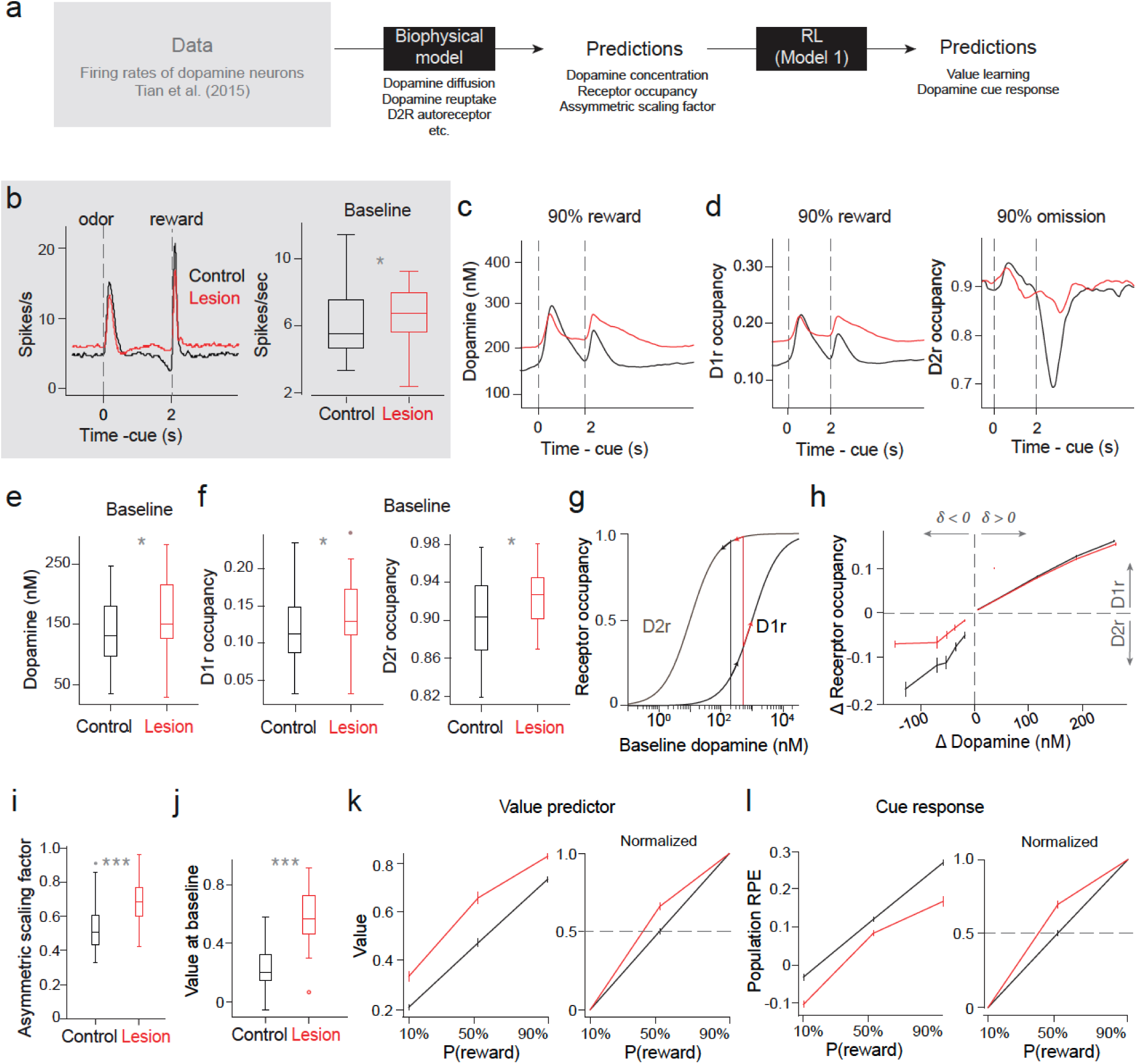
Model 1 can account for optimistic biases in reward-seeking behavior and cue-evoked dopamine responses. **a**. Schematic of the analysis. A biophysical model was used to predict dopamine concentrations, receptor occupancies, and value learning based on firing rates of dopamine neurons recorded in Tian et al. (2015). **b**. Average firing rates of dopamine neurons across the population for the control and lesion groups (left, n= 45 control group, n= 44 lesion group). Baseline firing rates were significantly greater in the lesion compared to the control group (right) (U-statistic = –2.429, *P* = 0.0151, single-sided Mann-Whitney U-test). **c**. Dopamine concentrations predicted from the firing rates of dopamine neurons based on the biophysical model of dopamine. Predictions for 90% reward trials are shown. **d**. Receptor occupancies predicted by the same biophysical model. Predictions for rewarded (left) and reward omission (right) trials in 90% reward trials are shown separately for D1R (left) and D2R (right), respectively (*n* = 45 control group, *n* = 44 lesion group). **e**. Mean dopamine concentrations at baseline predicted by the model (U-statistic = –2.109, *P* = 0.0175, single-sided Mann-Whitney U-test). **f**. Mean receptor occupancies at baseline predicted by the model (*n* = 45 control group, *n* = 44 lesion group). There is a significant increase in occupancies for both the D1R and D2R in the lesion compared to the control group (U-statistic = –2.1664, *P* = 0.0151, U-statistic = –2.1328, *P* = 0.0165 for D1R and D2R respectively, single-sided Mann-Whitney U-test). **g**. Schematic showing the model predicted changes in dopamine concentrations and receptor occupancies for the control (black) and lesion (red) groups. The arrows depict the increase or decrease in occupancy for a positive or negative dopamine transient of a fixed magnitude. **h**. Changes in receptor occupancy as a function of dopamine transients predicted by the model. The slope for the positive and negative domains correspond to the receptor sensitivities of D1R and D2R (*α*^+^, *α*^−^), respectively. **i**. Asymmetric scaling factors derived from the receptors’ sensitivities for the control and lesion groups (i.e., *τ* in model 1, *n* = 45 control group, *n* = 44 lesion group). There was a significant increase in the lesion group with respect to controls (U-statistic = –12.205, *P* < 1.0 x 10^−6^, single-sided Mann-Whitney U-test). Note that the increase in the asymmetry was driven mainly due to decreases in D2R sensitivity (panel h) **j**. Value predictions at baseline in a TD learning model trained with the receptor sensitivities derived from the biophysical. There was a significant increase in the value predictors at baseline in the model using the lesion group’s derived parameters with respect to control. controls (*t*-statistic = –6.417, P < 1.0 x 10^−6^, *t*-test). **k**.Value predictions at convergence of a TD learning model trained using the assumptions of Model 1 and the asymmetric scaling factors derived from receptors’ sensitivities predicted by the biophysical model. The model led to a significant increase in the value predictions for all cues (U-statistic = –4.690, *P* < 1.0 x 10^−4^, U-statistic = –4.734, *P* < 1.0 x 10^−4^, U-statistic = –4.602, *P* < 1.0 x 10^−4^, single-sided Mann-Whitney U-test, for the 10%, 50% and 90% reward probability cues) and an optimistic bias in the normalized value prediction to the 50% reward probability cue (*t*-statistic = –5.576, *P* < 1.0 x 10^−4^, *t*-test) in accordance with the anticipatory licking observed in the data. **l**. Predicted cue responses. There is an overall decrease in RPEs in lesioned animals (left) due to an increase in the baseline (pre-cue) value prediction (U-statistic =4.932, *P* < 1.0 x 10^−5^, U-statistic = –3.658, *P* = 0.00025, U-statistic = 4.734, *P* < 1.0 x 10^−4^, single-sided Mann-Whitney U-test for the 10%, 50% and 90% reward probability cues), which is consistent with the decreases in the absolute magnitudes of dopamine cue-evoked responses in the lesion group (Fig. 4c). The normalized TD errors at for the 50% reward probability cue show signs of an optimistic bias (U-statistic = –4.624, P < 1.0 x 10^−4^, single-sided Mann–Whitney U-test). Centre of box plot shows the median; edges are 25th and 75th percentiles; and whiskers are the most extreme data points not considered as outliers.

To quantitatively predict dopamine concentrations in the striatum and resulting receptor occupancies of D1R and D2R, we used a biophysical model commonly used in the field^59^ (Fig.6a). This model has the firing rate of dopamine neurons as its input, and considers diffusion of dopamine, dopamine reuptake, and D2-autorreceptor-mediated inhibition of dopamine release to predict the dopamine concentration in the striatum (Fig. 6b,e). In addition, it considers the affinities of D1R and D2R to estimate their occupancy levels (Fig. 6c,f). After estimating these two variables (dopamine concentration and receptor occupancy), we derived the receptor sensitivities (Fig. 6g-h). The receptor sensitivities were quantified as the slope of the resultant changes in receptor occupancy given the observed baseline and phasic responses of dopamine neurons. We then trained Model 1 using the receptor sensitivities as learning rates (*α*^+^ and *α*^−^) for both control and lesioned animals.

The biophysical model indeed supported that the observed change in dopamine neuron firing can cause a significant increase in dopamine concentration (Fig. 6e) and in D1 and D2 receptor occupancies at baseline (Fig. 6g). These changes are expected to cause a significant asymmetry in receptor sensitivities favoring D1 receptors over D2 receptors (Fig. 6h-i).

These receptor sensitivities were directly used as the asymmetric learning rates in a temporal-difference (TD) learning version of Model 1 (see Methods 1.3, 3.3). After training, the model incorporating the predicted asymmetries in learning rates (*α*^+^, *α*^−^) produced optimistic biases in value predictions and in normalized cue responses, similar to those observed in lesioned animals (Fig. 6k-l). The model simulating control animals developed no significant biases.

Additionally, the overall decrease in the magnitude of cue responses, observed in lesioned animals, was reproduced in Model 1 using TD learning (Fig. 6k). This occurs because TD learning calculates RPEs based on the change in values between before and after cue presentation, and the “baseline” (pre-cue) reward expectation was also increased by optimistic value learning (Fig. 6k). These results, together, indicate that Model 1 provides a parsimonious account of the data: a change in baseline firing of dopamine neurons, rather than changes in phasic responses, is the likely mechanism that led to optimistic biases in reward-seeking behavior as well as cue-evoked dopamine responses in habenula lesioned animals.

#### Model 1 and model 2 play complementary roles in the encoding of asymmetric learning rates

Although Model 2 did not explain the optimistic biases in the data in habenula-lesioned mice, the distributional RL version of Model 2 explained other features of the data (Extended Data Fig. 3-4). As mentioned, in both control and lesioned animals, asymmetric scaling factors tiled a wide range between 0 and 1^36^ (Extended Data Fig.4). Furthermore, cue-evoked responses of individual neurons showed a wider distribution than what is expected by noise (Figure 4d). Finally, the core prediction of distributional RL – a positive correlation between the asymmetric scaling factors of the RPE responses of individual dopamine neurons and their zero-crossing points^36^ – was also present in controls and after Hb lesions. Together these results support that the basic features of distributional RL are present in a way consistent with Model 2.

To complement this analysis, we tested whether Model 2 could have explained the signatures of the data if asymmetric scaling factors (τ) derived from dopamine responses were indeed overall biased (Extended Data Fig. 5). As expected from the model’s fixed-point analysis (Methods 1.3), if we imposed a shift in the mean of the distribution of asymmetric scaling factors (i.e., *τ*_*population*_ > 0.5), the value predictors indeed exhibited optimistic biases (Extended Data Fig. 5e,f). However, the model did not reproduce the optimistic bias in cue-induced TD errors observed in the data (Extended Data Fig.5g,h). This is due to an interaction of the biases in prediction at “baseline” (pre-cue) and the cue, together with the optimistic asymmetry in the scaling of the TD errors at cue themselves. Importantly, this was found in both versions of Model 2, distributional and risk-sensitive RL (Extended Data Fig. 5a-d and e-h). The difficulty of explaining biased dopaminergic cue responses further makes the Model 2 an unlikely mechanism to explain the optimistic biases in the data.

Altogether the data supports a model in which the mechanisms of Model 1 and 2 play complementary roles in the encoding of asymmetric learning rates. The mechanism of Model 2 explains the variability in single neuron responses, consistent with the expectile code in distributional RL. On the other hand, the mechanism of Model 1 at the population level, generating asymmetries in learning rates and biases in value expectations, which might require context-dependent regulation^60^.

Taken together, the above results suggest that Model 1 and 2 coexist in the brain. This can be formalized as follows:

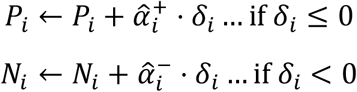

where:

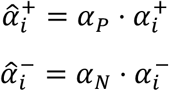

where 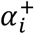 and 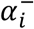 correspond to the asymmetric scaling of dopamine RPEs at the single-cell level (Model 2) and *α*_*P*_ and *α*_*N*_ correspond to the asymmetric scaling of synaptic plasticity in D1R and D2R at the population level (Model 1).

### Linking asymmetric learning and baseline dopamine levels in healthy subjects

#### Cools et al., (2009)^24^

There have been very few studies that examined the relationship between baseline dopamine levels and asymmetry in learning from positive and negative outcomes. As a rare case for such examinations, Cools et al.^24^ provided intriguing data in humans. They compared the performance in reversal learning and the quantity called ‘dopamine synthesis capacity’. Dopamine synthesis capacity is estimated by injecting the positron emission tomography (PET) tracer [^18^F]fluorometatyrosine (FMT) and is thought to be correlated with baseline dopamine levels^61,62^. This study found that higher dopamine synthesis capacity was correlated with better learning from gains but not with learning from losses (Fig. 7b). As a result, in reversal learning, subjects with higher dopamine synthesis capacity learned from gains than losses, reported as the ‘relative reversal learning (RRL)’ index in their study (Fig. 7b). This result, thus, provides direct evidence supporting our Model 1.

**Figure 7.**
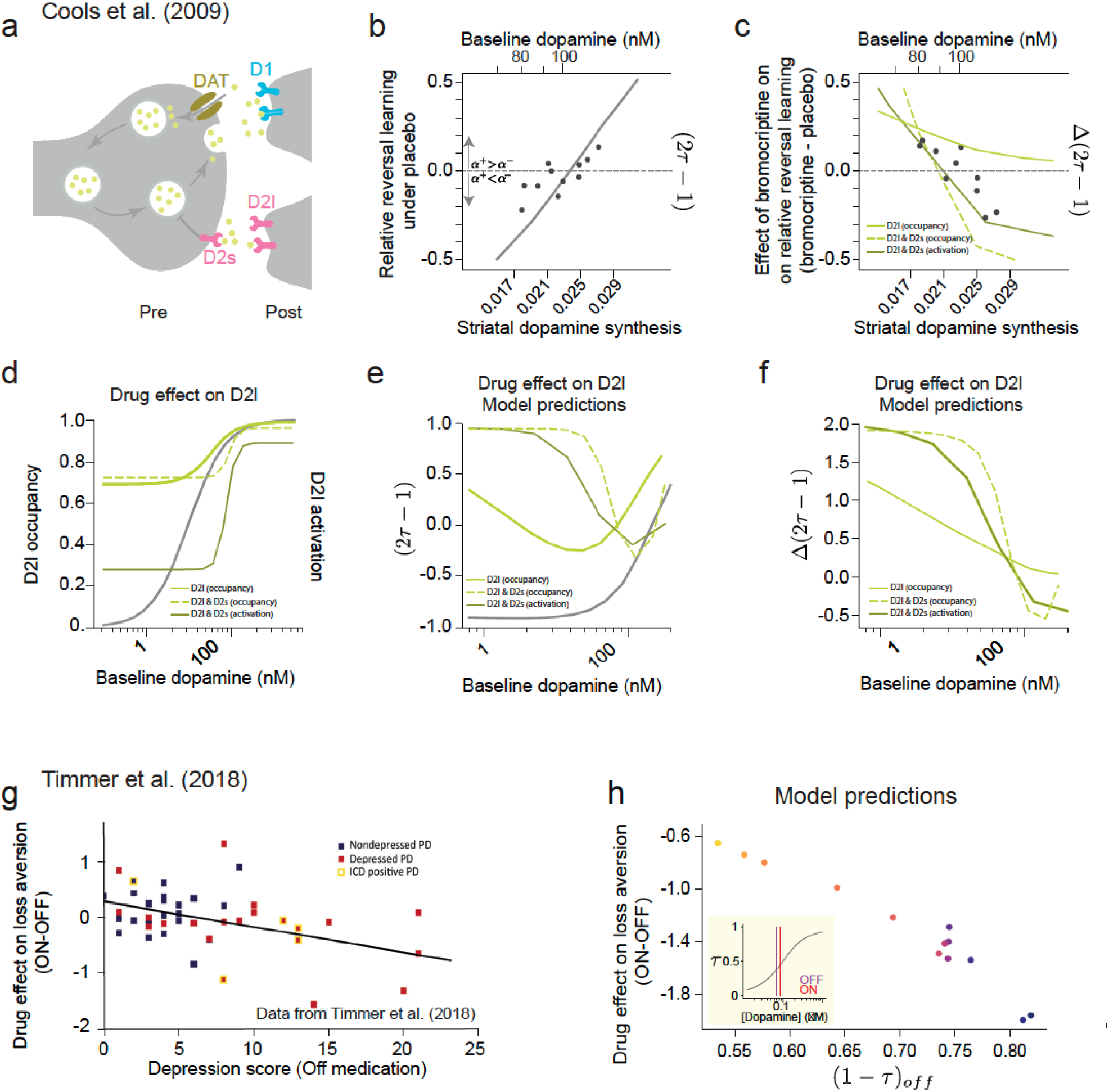
Model 1 predicts asymmetric learning rates in healthy humans given inter-individual differences in baseline dopamine, and in Parkinson’s disease patients given inter-individual differences in depressive-like symptoms. **a**. Schematic of the events occurring at dopaminergic axon terminal. Pre- and post-synaptic sites predominantly express D2s (short) and D2l (long) subtypes, respectively. **b**. ”Relative reversal learning (RRL)” under placebo conditions as a function of dopamine striatal synthesis capacity measured with PET radio imaging (black dots, left y-axis, bottom x-axis). Figure taken from Cools et al. (2009)^24^. Positive values of RRL indicate a bias favoring learning from gains relative to losses, and vice versa for negative values of RRL. There was a positive relationship between RRL and dopamine synthesis capacity. Model 1 predictions of RRL (2*τ* − 1 in Model 1) as a function of baseline dopamine using the receptors occupancy curve, recapitulate the positive relationship shown in the results from Cools et al. (2009)^24^ (gray line, right y-axis, top x-axis). **c**.The change in RRL induced by bromocriptine was negatively correlated with striatal dopamine synthesis capacity. Figure from Cools et al. (2009)^24^ (black dots, left y-axis, bottom x-axis). Model 1 recapitulates qualitatively the effect of bromocriptine in RRL, equivalent to Δ(2*τ* − 1). The solid light green line represents the Δ(2*τ* − 1) when considering bromocriptine’s effect on *D2l occupancy only*; the dashed line represents the Δ(2*τ* − 1) when both *D2l and D2s occupancy* was considered; and the dark green line represents the Δ(2*τ* − 1) when both *D2l and D2s activation* was considered (this includes the fact that bromocriptine is a partial agonist for the D2l and D2s receptors). The curves were obtained by imposing a concentration of 10^0.8^ nM of bromocriptine in the biophysical model. **d**. Receptor occupancy curves for the D2l receptors at baseline (grey line) and in the presence of 10^0.8^ nM of bromocriptine: Solid light green line corresponds to considering bromocriptine’s effects on D2l receptors occupancy alone; dashed line, corresponds to considering bromocriptine’s effects on both D2l and D2s receptors; solid dark green line corresponds to considering bromocriptine’s effect on the activation curves of both D2l and D2s receptors. The binding of the drug to the D2l receptors alone causes an increase in occupancy. This happens to a larger extent when starting from a low dopamine level at baseline than in high dopamine levels. The binding of the drug to D2s receptors in addition to D2l receptors causes a rightward shift in the curves. The activation levels are lower than 1 even at the drug levels where occupancy is close to 1, due to the lower efficiency of bromocriptine in receptor activation (Methos 4.1). See Extended Figure 6 and 7 for the effect of changing bromocriptine’s concentration and efficiency of activation. **e**. Same as in panel **d**. but now reporting (2*τ* − 1) calculated from the D2l receptor’s occupancy and activation curves. An increase in 2*τ* − 1 happens to a larger extent when starting from a low dopamine level than from high dopamine level. **f**. Same as in panel **d**. but now reporting Δ(2*τ* − 1) calculated from the D2l receptor’s occupancy and activation curves. Model 1 recapitulates qualitatively the effect of bromocriptine on RRL. **g**. The effect of PD medication (L-DOPA) on loss aversion is negatively correlated with their off-medication depression score. Figure from Timmer et al. (2018)^71^. **h**. Model 1 recapitulates qualitatively the effect of PD medication in loss aversion. We assumed that the asymmetry in favor of learning from losses relative to gains (1 − *τ*)_*off*_ scales with the baseline dopamine levels. Given this, we derived a distribution of off-medication baseline dopamine levels centered around the mean (1 − *τ*)_*off*_ derived from the data of Timmer et al. (2018)^71^ (see methods 0). We then imposed a fixed increase in baseline dopamine to simulate L-DOPA effects. We derived the new loss-aversion parameter (1 − *τ*)_*on*_ at the shifted baseline dopamine levels. The y-axis shows the change in loss aversion for each sample of the distribution of baseline dopamine levels. If the off-medication depression score is correlated with (1 − *τ*)_*off*_ then model would predict the result in Timmer et al. (2018)^71^. PD: Parkinson’s disease.

In addition, they found that dopamine synthesis capacity predicts the effectiveness of bromocriptine (D2 partial agonist) in altering learning rate asymmetry: bromocriptine’s ability to bias learning from gains over losses (i.e., positive change in RRL) was negatively correlated with dopamine synthesis capacity (Fig. 7c). We found that this result can also be explained by Model 1. For this, we simulated the effects of bromocriptine with the biophysical model used above, and derived the asymmetric learning rates from the slopes of the D2R occupancy (Fig. 7d, Extended Data Fig. 6a,b) or activation curves (Fig. 7d, Extended Data Fig. 6c,d). The RRL parameter reported by Cools et al. corresponds to the asymmetric scaling factor *τ*, and is equivalent to (2*τ* − 1) (as described in the Methods 4.1). We then computed what would be the change in this parameter Δ(2*τ* − 1) induced by bromocriptine (Fig. 7e-f, Extended Data Fig. 6e-l).

This analysis revealed that by considering the asymmetries in learning rates induced by changes in the baseline occupancy of the receptors, our model can capture their results in a qualitative manner. Intuitively, the less dopamine there is at baseline, the lower the occupancy of D2R at placebo conditions. This leads to a larger increase in D2R occupancy induced by D2 agonist in low dopamine baseline conditions (Fig. 7d, Extended Data Fig. 6a) and, thus, a larger increase in asymmetry in learning form gains over losses, if D1R occupancy is kept fixed These effects still hold even if we consider, in addition to bromocriptine’s effects in postsynaptic receptors (D2 long or D2l), its effect on inhibition of dopamine release via presynaptic (D2 short or D2s) autoreceptors^63,64^ (Fig. 7d, Extended Data Fig. 6b). This can be simulated as a decrease in dopamine level, which leads to a shift in the occupancy curves to the right. Finally, we can consider effect of the *partial* agonism of the drug, that leads to a lower activation level of receptors even if the occupancy is maximal (Fig. 7d, Extended Data Fig. 6c-d). Even after considering this last factor, the results remain qualitatively the same as those found in the original study. These results were robust to a relatively wide range of values in the simulation’s parameters (Extended Data Fig. 7, 8).

### Linking psychiatric conditions to baseline dopamine levels

#### Timmer et al., 2018

Various psychiatric disorders are characterized by abnormal future predictions or mood. Our Model 1 raise the possibility that an overall decrease in baseline dopamine level in the striatum would enhance learning from negative outcomes over learning from positive outcomes leading to persistent pessimistic future value expectations, a hallmark of depressive-like symptoms (Fig. 3a,b). A piece of evidence supporting this in the human literature is the greater learning rates for losses over gains in patients with Parkinson’s disease (PD)^22,25^, its comorbidity with depression^25,65^ that can precede the PD diagnosis^65–67^, and the reports of decreased dopamine transporter binding in the ventral striatum in depressed PD patients compared to non-depressed PD patients^68,69^.

In addition, the progression of dopaminergic axonal loss in PD is topographically unbalanced: the axonal loss is more prominent in the dorsal striatal regions^70^ than in the ventral ones. This leads to uneven dopamine baseline levels across the striatum that would interact with the global increases in dopamine induced by dopaminergic medications in PD patients. We hypothesize that a behavioral readout of the degree of this unevenness might be the presence or absence of depression as a comorbidity: *patients with depression might have lower dopamine levels in the ventral striatum*. Thus, if indeed baseline dopamine levels are correlated with depression, this comorbidity could be predictive of the effects of PD medication.

We examined a previous study that provided evidence for this hypothesis ^71^. Here, PD patients with and without depression history were tested in a gambling task, under presence or absence of medication (‘ON’ and ‘OFF’ medication states). The authors fitted a ‘loss aversion’ parameter to the behavioral performance, which is equivalent to 1 − *τ* in our model, under some assumptions (Methods). Their results were consistent with our model predictions. In the OFF-medication state, there was a (near-significant) main effect of depression group (with or without depression) on the learning rate asymmetry: patients with a depression history tended to be more loss averse than nondepressed patients (*P* = 0.052). This is consistent with a decrease of dopamine levels in the ventral striatum and thus a regime of *α*^+^ < *α*^−^ in value learning. Importantly, in the ON-medication state, the medication effects on the asymmetry in learning rates were predicted by the degree of severity of depression: patients with larger depression scores exhibited greater drug-induced decreases in loss aversion (Fig. 7g), which would correspond to an increase in 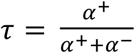 in our model This is consistent with our Model 1: higher degrees of depression might be correlated with lower levels of baseline dopamine, making the D1R sensitivity more susceptible to an artificial increase in baseline dopamine with L-DOPA medication (Fig. 7h; further details discussed in Methods).

## Discussion

A hallmark of various psychiatric disorders is overly optimistic or pessimistic predictions about the future. Using RL models, we sought to identify potential biological mechanisms that give rise to biased value predictions, with a particular focus on the roles of phasic versus tonic dopamine. Our results demonstrate that variations in tonic dopamine levels can modulate the efficacy of synaptic plasticity induced by positive versus negative RPEs, thereby resulting in biased value learning (Model 1). This effect arises due to sigmoidal shapes of the dose-occupancy curves and different affinities of dopamine receptors (D1R and D2R); alterations in the tonic dopamine level result in changes in the slope of the dose-occupancy curve (and thus, sensitivity) of dopamine receptors at the baseline dopamine concentration. We show that this mechanism offers a simple explanation for how changes in tonic dopamine levels can result in biased value learning in a few examples of value learning in mice and humans. Additionally, we show that this mechanism may underlie symptoms of various psychiatric and neurological disorders. Although altered phasic dopamine responses could have been a natural suspect as a candidate mechanism for biased value learning^37,38^, our study provides a novel mechanism; the interaction between tonic and phasic dopamine can give rise to biased value learning, even when phasic dopamine responses remain relatively unchanged.

### The impact of properties of dopamine receptors on reinforcement learning (RL)

Our results highlight the importance of considering properties of dopamine receptors and neural circuit architecture (i.e., direct and indirect pathways) in RL models. Based on different affinities of dopamine D1 and D2 receptors, it has been proposed that D1- and D2-SPNs play predominant roles in learning from positive and negative dopamine responses^32,72–75^. In support of this idea, recent experiments have demonstrated that PKA signaling in D1- and D2-SPNs is primarily driven by a phasic increase and decrease of dopamine, respectively^35^. Furthermore, LTP-like changes in D1- and D2-SPNs are triggered by a phasic increase and decrease of dopamine, respectively^33,34^. These recent pieces of evidence suggest that these plasticity rules are a basic principle of the RL circuitry in the brain. Here we explored the properties of this RL model and found the impact of the shape (slope) of receptor occupancy curves and showed that the tonic dopamine levels can modulate the relative efficacy of learning from positive versus negative RPEs.

One assumption in our model is that after a change in the tonic dopamine level, intracellular signaling reaches a steady inactive state, and it is the *change* in receptor occupancy that matters for inducing synaptic plasticity, rather than the *absolute* level of receptor occupancy reached during phasic dopamine responses. We note that absolute level might also contribute, yet it is expected that an increase or decrease in absolute occupancy levels will cause effects in the same direction as the effects of relative change that we explored in this study.

Additionally, our model, which incorporates the new plasticity rules, the opponent circuit architecture and properties of D1/D2 dopamine receptors, provides insights into the basic design principle of the brain’s RL circuit. It should be noted that the dose occupancy curves were plotted as a function of the logarithm of dopamine concentration, which makes the occupancy curves into sigmoidal shapes (Fig. 3, Extended Data Fig. 9). This logarithmic scaling is important in two ways. First, considering two sigmoidal curves for D1R and D2R together, the curves are approximately *symmetric* around the normal baseline dopamine level (Fig. 3a, Normal). Second, logarithmic scaling means that a fold-change in dopamine concentration will lead to the same leftward or rightward shift in these plots. It has long been argued that signaling of RPEs by dopamine neurons is curtailed by the fact that dopamine neurons have relatively low firing rates (2-8 spikes per second), and inhibitory responses of dopamine neurons tend to be smaller than excitatory responses^76,77^. Importantly, if we consider logarithmic scaling of dopamine concentration, the problem of this asymmetry is substantially mitigated (Extended Data Fig. 10). For example, with the baseline firing of 6 spikes per second, a phasic increase to 18 spikes per second and a phasic decrease to 2 spikes per second will cause the identical *fold*-changes in spiking (i.e., 3-fold changes in both directions), which would lead to a similar *fold*-changes in dopamine levels (Extended Data Fig. 11) and similar *percent* increase and decrease in receptor occupancy in D1R and D2R, respectively (Fig. 3a). Consequently, the system achieves symmetry in its response to positive and negative dopamine responses of observed magnitudes.

This may help understand *why* the basal ganglia circuit employs the opponent circuit architecture in the first place. In the model used in the present study, the value is encoded as the difference between the activity of D1- and D2-SPNs (*V* = *P* − *N*)^32^. We propose that this opponent circuit architecture, together with the logarithmic scaling of dopamine concentration, allows the system to effectively learn and encode both positive and negative values, which are contributed by the increase of firing in D1- and D2-SPNs, respectively. This would allow to expand the dynamic range of value coding, without requiring high baseline firing rates. Thus, at the normal dopamine baseline, learning from positive and negative dopamine responses is well balanced. When the tonic dopamine level deviates from the normal level, however, then the symmetry is broken and value learning becomes biased, as explored in the present study.

### The role of tonic dopamine levels in psychiatric disorders

As mentioned above, our modeling results provide an account for biased value predictions observed in various psychiatric and neurological conditions. For one, our model provides a link between findings in depressive-like states in animal models and the value learning biases exhibited by humans.

In a rodent model of depression, it has been reported that spontaneous activity of dopamine neurons is decreased^78^ (but see^79,80^). In addition, decreased spontaneous firing of dopamine neurons has been observed as a result of chronic pain-induced adaptations that correlate with anhedonia-like behavior^81^. Furthermore, maternal deprivation, which increases susceptibility to anhedonia, led to an upregulation of D2R expression in the VTA^82^, which is expected to decrease the excitability of dopamine neurons via its autoreceptor function. Finally, chronic administration of corticosteroids, a method to mimic anxiety and anhedonia-like states, results in an increase in somatodendritic dopamine concentration which then decreases dopamine excitability via D2R hyper-activation^83^. These results of decreased dopamine excitability correlated with anhedonia-like states are consistent with findings of increased burst firing of lateral habenula (LHb) neurons^56^ and potentiation of glutamatergic inputs onto the habenula^57^ in depression models. This is further supported by reports that depressive-like behavioral phenotypes can be ameliorated by optogenetic activation of dopamine neurons^84^ and the anti-depressant effects of ketamine might be mediated by the inhibition of bursting in the LHb^58^

The mechanism by which a broad change in dopamine excitability could lead to depressive-like states remains to be revealed. Just by assuming that a decrease in spontaneous firing leads to a decrease in baseline dopamine level in the striatum, our model readily predicts that learning from negative outcomes will be emphasized over learning from positive outcomes (Fig. 3a,b), as has been reported in some studies of patients with major depressive disorder (MDD)^1^. In addition, RL agents learning in these conditions exhibit enhanced risk-aversive behavior, pessimistic outcome expectations, and increased sensitivity to losses compared to gains, all of which are signatures of depressive-like conditions^1,5,21,85,86^. This contrasts with findings of increased dopamine synthesis capacity in pathological gambling patients^87^, who show the opposite behavioral signatures^3^.

An additional line of research relevant to our proposal is PD patients and pathological gambling as a comorbidity. Previous work has emphasized the interaction between the degree of dopaminergic loss and the effects of PD medications^88–90^, which can sometimes result in the development of addictive disorders such as pathological gambling. As mentioned, the loss of dopaminergic axons in PD patients has been reported to happen predominantly in the dorsal regions of the striatum^70^. Thus, at the onset of the motor impairment symptoms, which is when L-DOPA medication tends to be prescribed, dopamine level is expected to be low in the dorsal striatum while it might be relatively intact in the ventral striatum. This can lead to ‘overdose’ of dopamine by medication: while L-DOPA might take dopamine levels in the dorsal striatum back to its original set-point, it might cause an ‘overdose’ in the ventral striatum^89,91^. Our model predicts that this overdose would lead to decreases in D2R sensitivity relative to D1R. Assuming that the ventral striatal regions have a dominant role in value learning, this would result in excessive optimistic expectations and risk seeking, two key behavioral features of pathological gambling and addictive disorders. We provided indirect evidence for this hypothesis; future work should directly test these predictions.

It should be noted that we did not consider changes in dopamine receptors density, which have also been related to value learning biases^92^ and psychiatric conditions^93^. Future studies should explore the influence of this additional factor in the encoding of asymmetric learning rates (i.e.,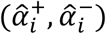.

### Tonic dopamine as a modulator of ‘mood’

Mood refers to a person’s emotional state as it relates to their overall sense of well-being. Although the exact neural substrate of mood remains unknown, recent studies have indicated that mood reflects not the absolute goodness of outcomes but rather on the discrepancy between actual and expected outcomes in recent history^13,14^. That is, mood depends on the cumulative sum of RPEs that occurred recently^13^. It has also been proposed that mood, in turn, affects the way we perceive and learn from positive and negative outcomes (RPEs)^13^.

Our model provides a unified mechanism for these two aspects of mood; both subjective feeling of mood and biased learning from positive versus negative outcomes can arise from changes in baseline dopamine levels which can be modulated by recent history of phasic dopamine responses. It was proposed that this history dependent modulation of learning is an adaptive mechanism that allows organisms to adapt quickly to slow changes in environments based on the “momentum” of whether the situation is changing in a better or worse direction on a slow timescale (e.g. seasonal change)^13,14^. The models presented in the present study may provide mechanistic insights into such mood-dependent modulation of learning and perception.

### Neural circuits for distributional reinforcement learning (RL)

We examined the possibility that optimistic biases in reward seeking behavior and dopamine cue responses observed in habenula-lesioned mice can be explained by Model 2, either based on risk-sensitive RL (the average response) or distributional RL (responses of a diverse set of individual dopamine neurons). We did not find evidence supporting this possibility. However, the present study makes two important contributions with respect to distributional RL. First, we can show that our model, which incorporated direct and indirect pathway architecture, can support distributional RL (Extended Data Fig. 12, 13). It would be interesting to examine what additional features and functions could be gained by having this opponent architecture. Second, we largely replicated the previous results^36^ using an independent data set. That is, the signatures of distributional RL were present in this data set (Extended Data Fig. 3-4), and dopamine cue-evoked responses did show an optimistic bias. This provides further evidence for a distributional code in dopamine neurons, and shows that there is an overall elevated distributional representation in dopamine cue responses in habenula lesioned animals.

### Concluding remarks

Taken together, our biologically inspired RL model provides a foundation to link findings in the brain and formal models of RL. Our work highlights a causal impact of baseline dopamine on biasing future value predictions, which may underlie mood and some abnormalities observed in psychiatric patients and could be used to regulate risk sensitive behavior.

## Methods

### 1. Reinforcement learning model

Here we provide formal definitions and the framework of reinforcement learning used in this study. We have focused our model formulations to the problem of *prediction*, in which an agent learns to predict the value function^11^. The problem of *control* (the problem of how an agent selects and executes actions) is not considered. In RL, an agent’s objective is to maximize the total cumulative rewards. It does so by learning the value associated with each state in an environment. For now, we will develop the model dropping the dependency on time within each episode. Here, the target to learn is the value function as defined by

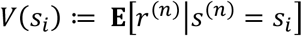

Where *r*^(*n*)^ is the reward experienced in the episode *n* (i.e., trial) of visiting state *s*_*i*_. Learning of *V*(*s*) is driven by reward prediction errors (RPEs, *δ*), the discrepancy between the actual and expected reward:

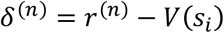

The value is updated for the experienced state according to:

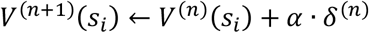

This is also known as the Rescorla-Wagner (RW) delta rule^94^. The reward in each trial is sampled from a reward distribution specific to a given state: *r*^(*n*)^∼*R*(*s*_*i*_). With the learning rule above, the value converges on the expected value of this reward distribution. This can be shown with a stochastic fixed-point approach; the convergence point is derived by obtaining the value of *V*(*s*_*i*_) at which the change in *V*(*s*_*i*_) from trial *n* to trial (*n* + 1) is expected to be zero (i.e., is zero on average):

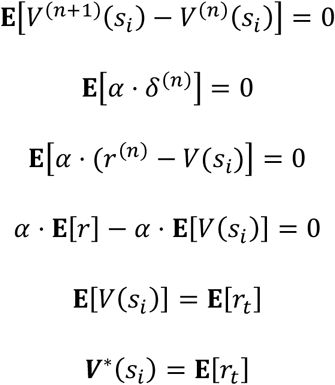

Where ***V***^*^(*s*_*i*_) is the stochastic fixed-point: the value around which *V*(*s*_*i*_) is expected to fluctuate after learning and corresponds to the learning target above.

#### 1.1. Temporal difference learning

Now we will consider time and extend the models to the temporal difference (TD) learning framework^11^. Dopamine responses have been shown to present key signatures of TD errors^95^. Therefore, TD learning models allow us to directly link the model variables to dopamine neural responses.

We can derive TD learning by defining a different environmental structure and learning objective. We start by considering arbitrary states (*s*_*t*_), which transition at each time step following a Markov process, and at each time step the agent samples a random reward from a probability distribution *r*_*t*_∼*R*(*s*_*t*_).

The learning objective is now the value of a given state *V*(*s*_*t*_) defined as the *expected cumulative sum of all future rewards* starting from state *s*. Rewards are discounted by a constant discounting factor (*γ*, with 0 ≤ *γ* ≤ 1) each time step. The expectation is taken over stochastic state transitions and sampled rewards:

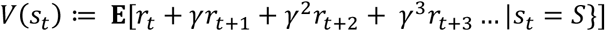

Where *s*_*t*_ is the state at time *t, r*_*t*_ is the reward sampled at time *t* and *V*(*s*_*t*_) is the value of the state *s*_*t*_. Since the environment and transitions are assumed to follow a Markov process, the equation above can be rewritten in a recursive manner. This is known as the Bellman equation^11^:

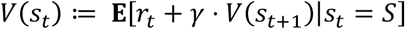

The agent approximates the true value *V*(*s*_*t*_) with a learned estimate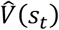. With this approximation, before learning converges, the estimates for the left- and right-hand sides are not equal. Thus, after sampling a reward *r*_*t*_∼*R*(*s*_*t*_) from the environment, the difference between the two terms in the Bellman equation represents the error in value prediction, called the temporal difference reward prediction error (TD RPE, *δ* below),

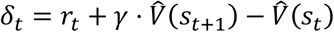

With *α* as the learning rate, the updates for the value estimates are:

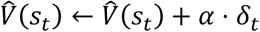

With this definition, the TD RPE contains the difference between the estimated value of states evaluated at consecutive time points. If we fix the discounting factor to be *γ* = 1, then 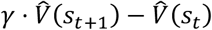 is the temporal derivative of the value function. As a result of this property, unexpected increases and decreases in value result in positive and negative transient changes in TD RPE, respectively^95^.

If dopamine responses encode TD RPEs, then cue-evoked responses can be formulated as:

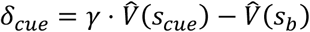

Where *δ*_*cue*_ is the TD RPE induced by the cue, 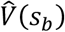 is the value prediction at baseline and 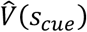 is the value prediction elicited by the cue (which reflects the expected value predicted by each trial type). As the 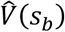 is the same across all trial types and represents the average value predictions across them, then *δ*_*cue*_ is dominated by the expected value of each trial type. This is a useful feature that we used in our simulations for the habenula lesion experiment.

#### 1.2. Distributional TD learning

In Results, we used a distributional TD learning model to test whether the subtle changes in the distribution of asymmetric scaling factors observed after lesions could lead to the observed changes in cue responses after learning.

In distributional TD learning, our learning objective is the entire distribution over cumulative discounted future rewards, instead of the value defined above^36,37,39^. We will call this the *return distribution, Z*(*s*_*t*_). We can thus write an analogue of the Bellman equation, the ‘distributional Bellman equation’:

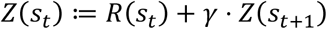

The target to learn in distributional TD is now *V*_*i*_(*s*_*t*_) that minimizes for the expectile regression loss:

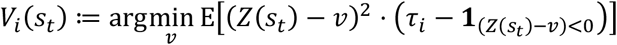

Where *Z*(*s*_*t*_) is a random variable, representing the return distribution, and 1_*f*_ is the indicator functions that is equal to 1 if the condition in the subscript {*f* ≔ (*Z*(*s*_*t*_) − *v*) < 0} is met, and 0 otherwise.

Minimizing the expectile regression loss makes *V*_*i*_(*s*_*t*_) to converge on the 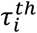 expectile of the return distribution^39^.

The target is learned by taking samples from the estimated return distribution^39^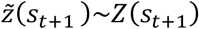and from the reward distribution *r*_*t*_∼*R*(*s*_*t*_), to compute the TD error:

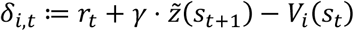

Note that 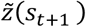 is random so the TD error is also random, and *δ*_*i,t*_ ≠ *r*_*t*_ + *γ* · *V*_*i*_(*s*_*t*+1_) − *V*_*i*_(*s*_*t*_). For more information regarding the sampling method employed in the simulations see Methods Section 3.3.

In addition, the updates are performed with different learning rates 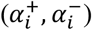 for positive and negative *δ*_*i*_. This asymmetry in the weighting of the errors used to update *V*_*i*_(*s*_*t*_) is essential to minimize the expectile regression loss.

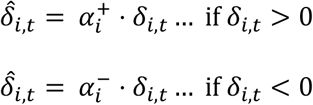

The reliance on a single sample for 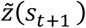 suffers from high variance. Therefore, for performing the updates we average across a set of *M* updates, each depending on a single sample *δ*_*i,t*_.

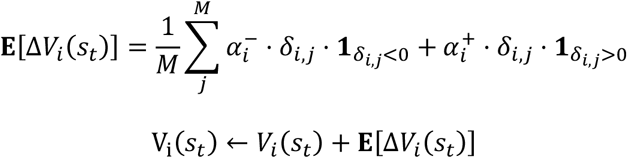

This learning rule will asymptotically converge to the *τ*_*i*_-th expectile of the return distribution^39^.

#### 1.3. TD learning with D1 and D2 populations

It is straightforward to extend the TD learning algorithm to have separate populations for D1 and D2 SPNs^32^. We employed this model to derive dopamine cue responses with Model 1 (Fig. 6i). In this model, the same computation of TD RPE of standard TD learning is still used. Yet, this model differs in the updates and computation of 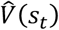.

As mentioned previously, the updates in the *P*_*i*_ and *N*_*i*_ populations happen exclusively with positive or negative TD RPEs, respectively:

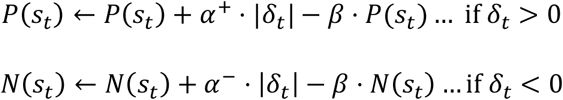

Where *α*^+^ and *α*^−^ are the learning rates for the *P* and *N* populations, that we postulate is modulated by baseline dopamine levels. The variable *β* ∈ (0,1) is the decay factor, which we keep constant throughout the simulations and serves to stabilize *P*(*s*_*t*_), *N*(*s*_*t*_).

The computation of value estimate 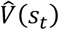 is given by:

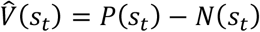

##### 1.3.1. Convergence of risk sensitive TD learning

We now discuss the convergence of the proposed TD learning algorithm with D1 and D2 populations. This analysis builds on the work in risk-sensitive reinforcement learning ^40^ and the already established results of convergence for stochastic iterative algorithms (e.g., TD learning) (Bertsekas & Tsitsiklis, 1996^96^, Proposition 4.4, p. 156).

**Theorem**: The results by Bertsekas & Tsitsiklis (1996)^96^ establish that, given a sequence *r*_*t*_ ∈ ℝ^*m*^ generated by the iterative algorithm:

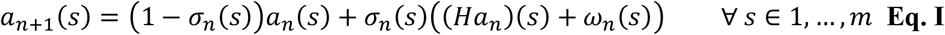

The variable *a*_*n*_ converges to the unique solution *a*^*^ of the equation: *Ha*^*^ = *a*^*^ with probability = 1, assuming the following conditions are fulfilled:

1. The step sizes σ_*i*_(*i*) are non-negative and satisfy:

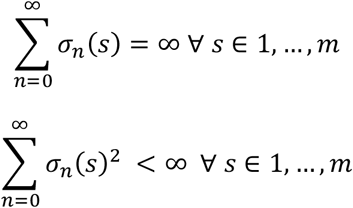
2. The noise term ω_*n*_(*s*) satisfies:
  - E[ω_*n*_(*s*)|ℱ_*n*_] = 0 ∀ *s, n*, where ℱ_*n*_ denotes the history of the process up to and including time step *n*
  - Given any norm ‖·‖ on ℝ^*m*^ there exist constants A and B such that: 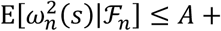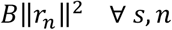
3. The mapping H is a *maximum norm contraction* (see below for definition)

To prove convergence, we will first discuss the case of risk-sensitive TD learning following ^40^ and then discuss TD learning with D1 and D2 populations.

We define the risk sensitive TD-learning rule as:

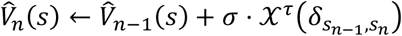

Where:

- 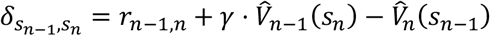
- The step index is *n* ∈ 0,…, ∞
- The step size σ is kept constant across iterations.
- For simplicity in calculations we follow ^40^ and make use of the operator *X*^*K*^ with *K* ∈ (−1,1)

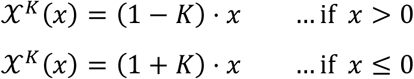

It is simple to show that the asymmetric scaling factor used in this paper is a scaled version of the operator. That is: *τ* = 0.5(1 − *K*) and 1 − *τ* = 0.5(1 + *K*).
- In addition, as in ^40^, given that the function *X*^*τ*^(*x*) is piece-wise differentiable we can apply the mean value theorem to show that for each pair of numbers (*a, b*) there exists a ℰ_*a,b,K*_ ∈ [1 − |*K*|, 1 + |*K*|], such that: 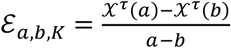. This relationship will become useful in the future.

We will re-format the update rule to better match the iterative algorithm above:

Adding and subtracting 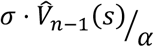

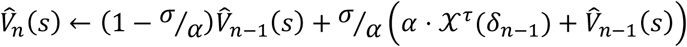

Defining an operator that will become useful:

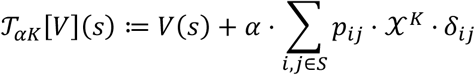

Defining the noise term as:

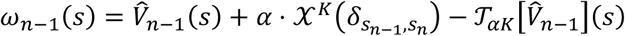

Then our update rule above becomes:

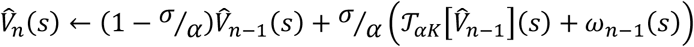

The formulation above can be directly compared to the one of stochastic iterative algorithm theorem (Eq.I), and now we can check whether the conditions for convergence are met.

1. The conditions for the learning rate, are a direct consequence of our choice of the parameter which is a constant in our model and 0 < *α* < 1.
2. It has been shown that showed that the conditions for the noise term ω_*n*−1_(*s*) as formulated above are satisfied^40^.
3. Finally, the operator *J*_*ατ*_[*V*](*s*) is a contraction mapping as also shown in Bersekas (1996)^40^. Therefore, the variable *V*_*n*_ converges to the unique solution *V*^*^ for which:

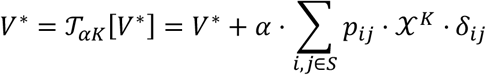

We elaborate now on the proof for the contraction mapping of the operator *J* _*ατ*_[*V*](*s*), as this will be useful for the proof of the D1 D2 TD learning model.

###### Definition of contraction mapping

Let (*X, d*) be a metric space (a set *X*, with a notion of distance, *d*, between points). A mapping *J*: *X* → *X* is a contraction mapping if there exists a constant *c*: 0 ≥ *c* > 1 such that for all *x* ∈ *X*:

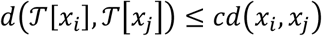

That is, a contraction mapping maps points closer together.

Elaborating now on the operator *J*_*ατ*_[*V*](*s*) and using |·| as our distance metric:

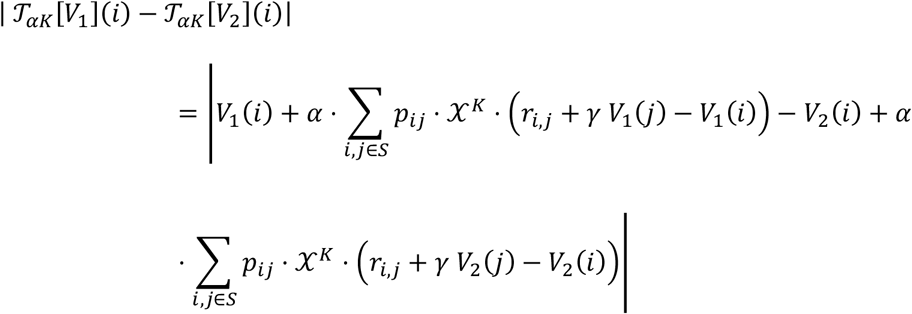

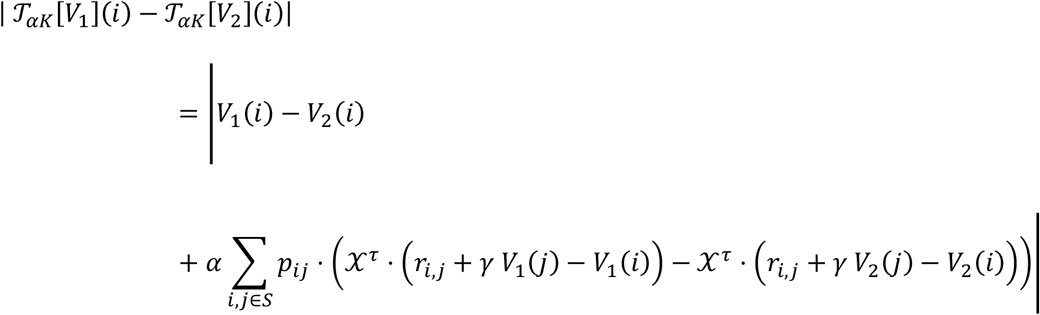

Using the relation defined above ℰ_*a,b,K*_ ·(*a* − *b*) = *X* ^*K*^(*a*) − *X*;^*K*^(*b*)

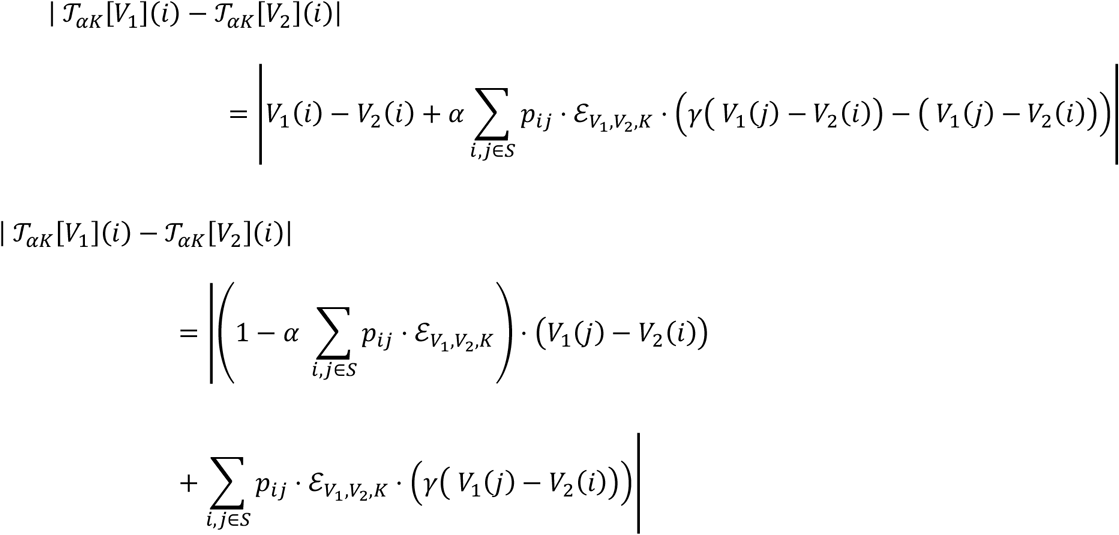

Given that ℰ_*a,b,K*_ ∈ [1 − |*K*|, 1 + |*K*|] and assuming *α* ∈ (0, (1 + |*K*|)^−1^):

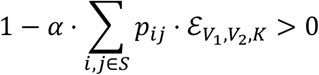

Taking this term outside the | · | and rearranging:

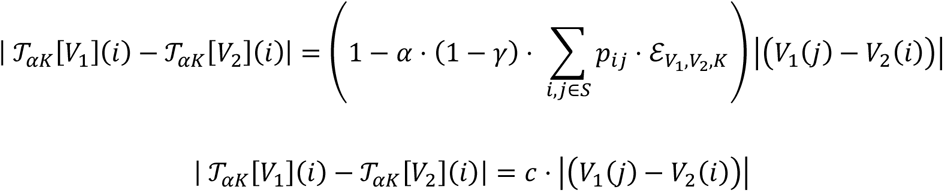

Where the term:

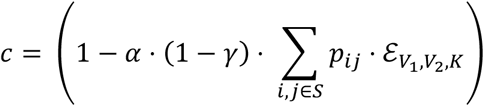

To get the upper boundary of *c* we use the minimum value for the sum, where 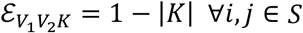. And use the assumption that *α* ∈ (0, (1 + |*K*|)^−1^):

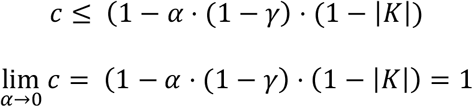

To get the lower boundary of *c* we use the maximum value for the sum, where 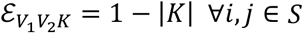. And use the assumption that *α* ∈ (0, (1 + |*K*|)^−1^):

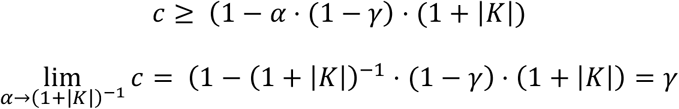

Therefore: *γ* < *c* < 1

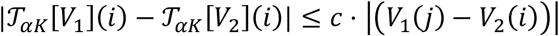

And the operator *J*_*ατ*_[*V*](*s*) is a contraction mapping, under the condition *α* ∈ (0, (1 + |*K*|)^−1^)

##### 1.3.2. Convergence of TD learning with D1 and D2 populations

We define the D1-D2 TD-learning rule as:

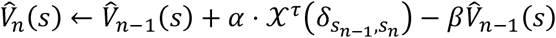

Note this update rule is analogous to the risk sensitive TD learning rule except for the last term that emerges from the decay factor in the *P, N* populations of our model.

Performing the same re-arrangement as above we reach:

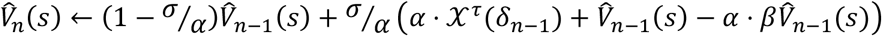

We define a new operator *J* ′_*αK*_ [*V*](*s*):

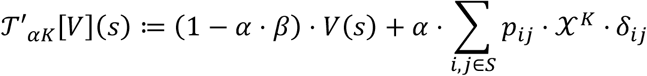

And the noise then is defined as:

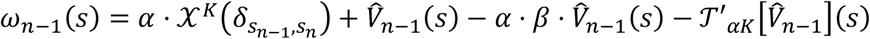

The update becomes:

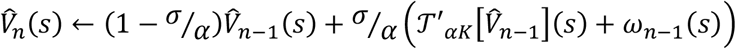

The noise term reduces to the same expression as the one of TD learning, and so it fullfils the requirements for the theorem of stochastic iterative algorithms. We will now test whether the operator *J* ′_*αK*_ [*V*](*s*) also represents a contraction map.

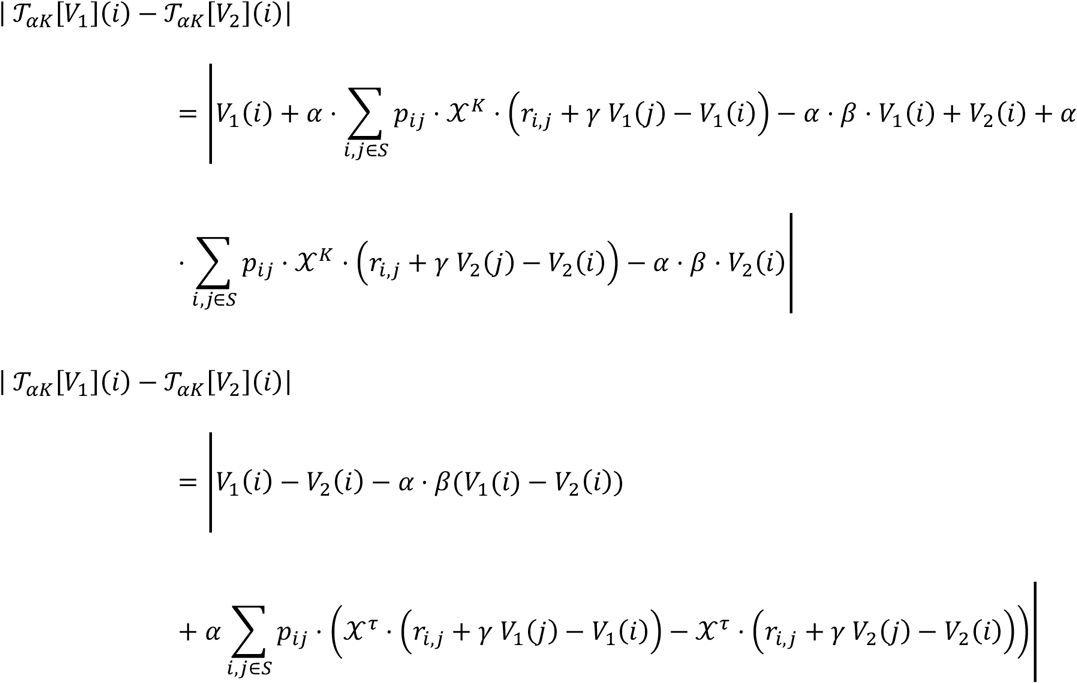

Using the relation defined above ℰ_*a,b,K*_ · (*a* − *b*) = *X* ^*K*^(*a*) − *X* ^*K*^(*b*)

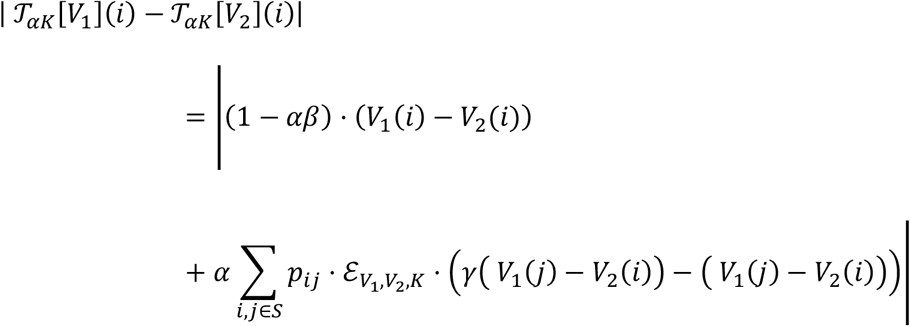

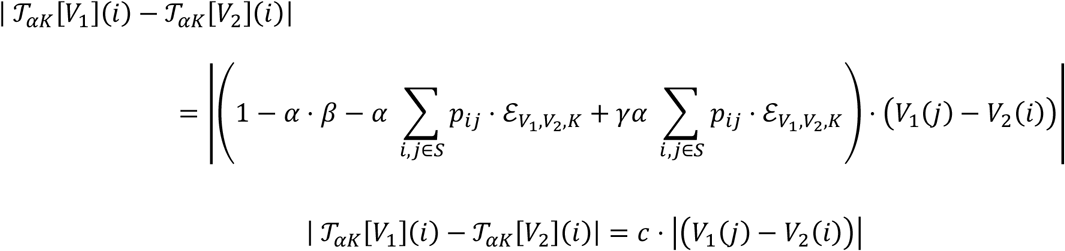

Where the term:

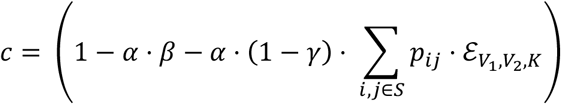

To get the upper boundary of *c* we use the minimum value for the sum, where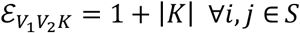, and use the assumption that *α* ∈ (0, (1 + |*K*|)^−1^):

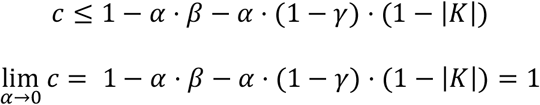

To get the lower boundary of *c* we use the maximum value for the sum, where 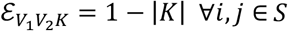, and use the assumption that *α* ∈ (0, (1 + |*K*|)^−1^):

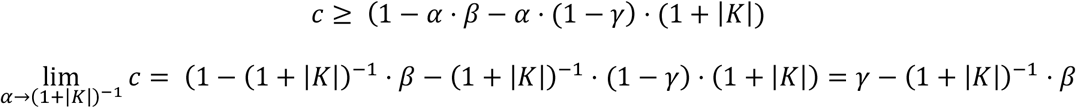

Given that we want *c* ≥ 0 we can find the parameter ranges to achieve this:

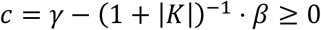

Given that: *K* ∈ (−1,1), we use the minimum value of |*K*| = 0 to find the limit of c:

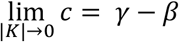

So the condition *γ* ≥ *β* needs to be present to keep: 0 ≤ *c* < 1.

Under these conditions, the operator *J* ′_*αK*_ [*V*](*s*) also represents a contraction map.

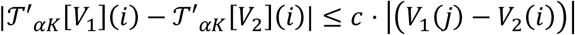

###### Stochastic fixed point for the value estimate

Having shown convergence of the algorithm, we will now derive the convergent points for our algorithm using stochastic fixed points. For clarity, we estimate the stochastic fixed point dropping the dependency on time.

If learning between *P* and *N* is symmetric *α*^+^ = *α*^−^ = *α*. We derive the convergent estimate of *V*(*s*_*t*_) with a fixed-point approach. First, we subtract the *P* and *N* update equations, to arrive to the update in the 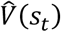 between the (*n*) and the (*n* + 1) update:

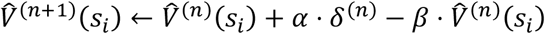

Where the superscripts indicate the iteration number. We can now derive the stochastic fixed-point for 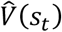:

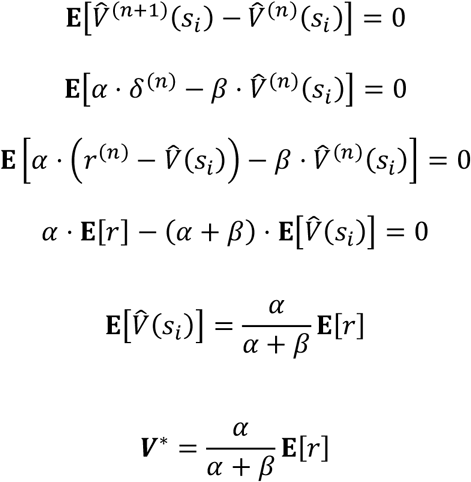

Where 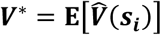 is the value around which 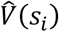 is expected to fluctuate after convergence.

Throughout this study, we have manipulated the learning rates between *P* and *N* to be asymmetric *α*^+^ ≠ *α*^−^ or, equivalently, *τ* ≠ 1 − *τ*. We can find the stochastic fixed point for this more general case:

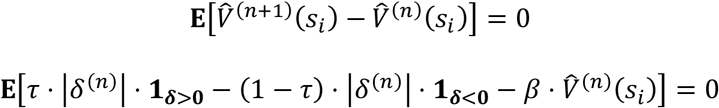

To take the expectation we use the definition: E[*X*] = ∑_*i*_ *p*(*x*_*i*_) · *x*_*i*_. For a Bernoulli distribution, *p*(*x*_*i*_) takes two values:

- *p*(*x*_*i*_) = *p* if reward is delivered and, thus *r* = 1, *δ*_*t*_ > 0,
- *p*(*x*_*i*_) = (1 − *p*) if reward is not delivered and, thus *r* = 0, *δ*_*t*_ < 0,

Therefore, we can resolve the expectation and expand the RPEs:

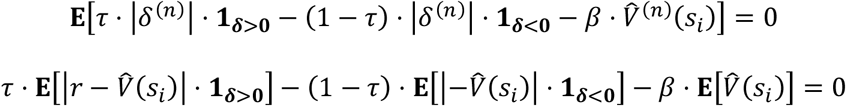

Taking the absolute values:

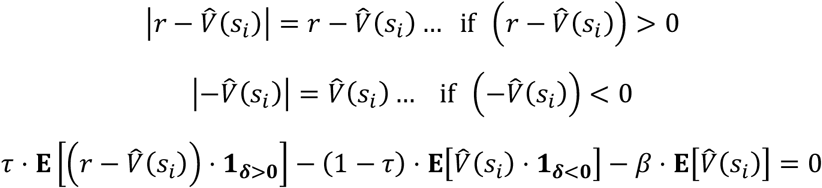

Replacing stochastic fixed point: 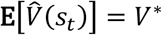 and taking the expectations:

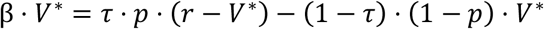

Rearranging and isolating *V*^*^, we obtain:

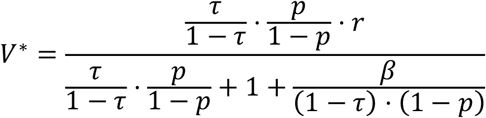

###### Stochastic fixed point for P and N populations

We have mentioned that the decay term (β) in the update equations serves to stabilize the estimates of the *P* and *N* populations (i.e., avoid infinite increases). We can observe the influence of *β* by computing the stochastic fixed points for these variables.

For the *P* population:

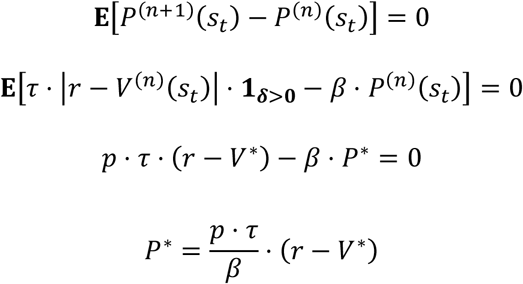

Similarly, for the *N* population:

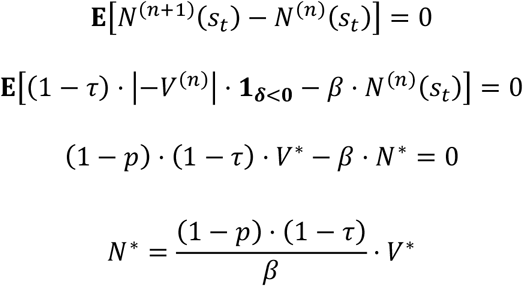

As it can be seen in the stochastic fixed points *P*^*^, *N*^*^, the term 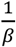 is a proportionality constant. Therefore:

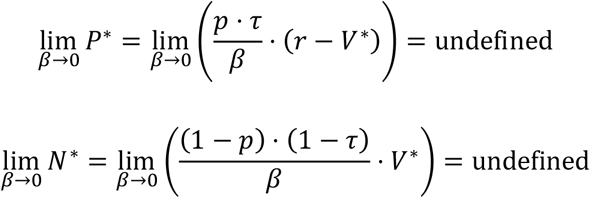

So, *β* ≠ 0 needs to be met for the stochastic fixed points *P*^*^, *N*^*^ to exist. In Extended Data Fig. 1 we show empirically that the convergence rate is slower as *β* gets closer to 0, but it is always achieved.

##### 1.3.3. Sensitivity of learned variables in D1-D2 model to parameters

The conditions for the D1-D2 model to reproduce the data from our habenula lesion experiment and some of the previous studies are that:

1. The bias in *V*^*^ induced by the asymmetric learning rates doesn’t change the monotonicity of the learned values as a function of the true expected value of the return distribution **E**[*R*(*s*)]. In other words, regardless of the level of ‘optimism’ or ‘pessimism’, *V*^*^ monotonically increases with **E**[*R*(*s*)].
2. Asymmetric learning rates change the concavity of *V*^*^ as a function of **E**[*R*(*s*)]: ‘Optimistic’ or ‘pessimistic’ value functions are concave or convex with respect to **E**[*R*(*s*)], respectively.

We will now analyze whether these conditions are met, considering the range of parameters of relevance: 0 < *τ* < 1, *r* ≠ 0 and 0 <*β* < 1

For the condition 1 to be met, the first derivative of *V*^*^ with respect to **E**[*R*(*s*)] should always be positive. In the case of Bernoulli return distributions, the derivative of *V*^*^ with respect to p(reward) is

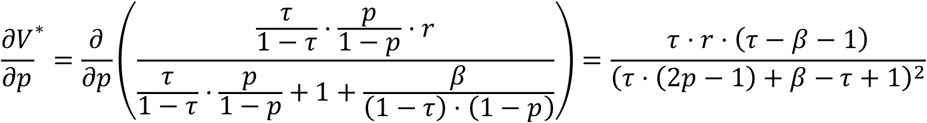

We can look at the fixed points of this expression, as they correspond to the value of *p* at which the derivative changes the sign. This expression has fixed points at: *τ* = 0, *r* = 0, and *β* = *τ* − 1. Given our parameters’ ranges: 0 < *τ* < 1, *r* ≠ 0 and 0 <β < 1, none of those fixed points are present within those ranges. In addition, it can be seen that this 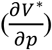 is positive for the parameter values within those ranges.

Therefore, knowing that the derivative won’t reach any fixed point, *V*^*^ is always a growing monotonic function with respect to *p*.

For the condition 2 to be met, we can analyze the second derivative of *V*^*^ with respect to **E**[*R*(*s*)] as it indicates the convexity of a function. The conditions to be mat are:

- *V*^*^ is convex if it is ‘pessimistic’: if 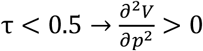
- *V*^*^ is concave if it is ‘optimistic: if 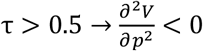

In the case of Bernoulli return distributions, we take the second derivative of *V*^*^ with respect to p(reward):

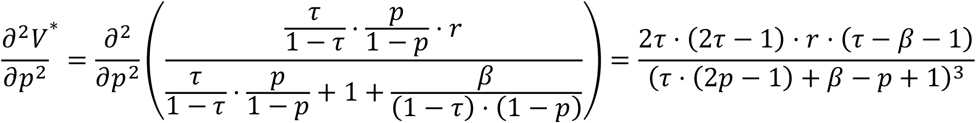

We can again look at the fixed points of this expression. These happen at: *τ* = 0, *r* = 0, and *β* = *τ* − 1 and *τ* = 0.5. Among them, the only fixed point within our parameters range is the latter. In addition, by replacing *τ* in the expression above, it is easily shown that it is positive if *τ* < 0.5 and negative if *τ* > 0.5. Thus, given that the ranges for the parameters are such that the second derivative won’t reach any other fixed point, condition 2 will always be met.

#### 1.4. Distributional TD learning with D1 and D2 populations

The signatures of distributional RL were preserved in dopamine neurons firing rates after habenula lesions (Extended Data Fig. 3-4). Therefore, we considered a third alternative to model 1 and 2, that assigns different functions to each of the mechanisms for asymmetric learning rates.

In this model (Extended Data Fig. 13) the single cell asymmetric scaling factors 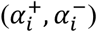 give rise to a distributional expectile code for value and are implemented at the level of the scaling of RPE-evoked responses of dopamine neurons:

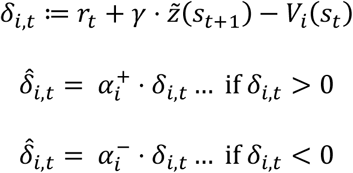

The modulation of receptor sensitivities, carried out downstream at the SPN level, gives rise to the global rescaling of the value updates (*η*^+^, *η*^−^) (Extended Data Fig. 13A):

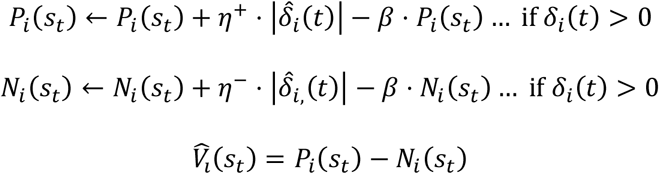

These set of update equations are equivalent to a modified version of the update equation of distributional RL:

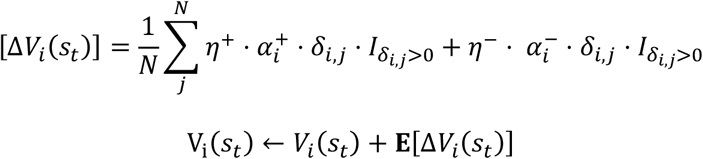

Thus, this model can give rise to biases in value learning (Extended Data Fig. 13), while keeping intact information about the value distribution. By employing the results from the biophysical model (Fig. 6), we found that this distributional TD model can parsimoniously explain all aspects of the data in the habenula lesion study (Extended Data Fig. 12B), including the features of a distributional code and the optimistic biases observed in behavior and dopamine cue-evoked responses (Extended Data Fig. 12).

#### 1.5. Dependency of model on assumption: Log vs. linear scaling of receptor occupancy curves

Through this work, we have used the dose-occupancy curves of D1 and D2 receptors to derive the receptor sensitivities that result in the asymmetric scaling factors in Model 1. It is important to note that the slopes of the receptor occupancy curve (= receptor sensitivity) were obtained from the receptor occupancy curves plotted as a function of log of dopamine concentrations.

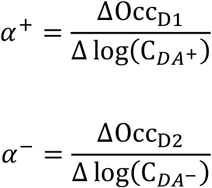

To show that this assumption is not essential, we now derive the receptors sensitivities assuming linear changes in dopamine levels due to RPE-evoked responses.

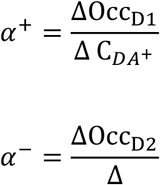

As shown in Extended Data Fig. 9, the choice of a linear versus log scale affects the absolute magnitude of the derived receptor sensitivities, but the normalized metric 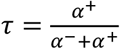 holds the same relationship to baseline dopamine levels with a small shift in the curve (Extended Data Fig.9, right panel). The normalized metric is the factor determining the update asymmetries and, thus, the stochastic fixed points at which the variables converge.

#### 1.6. Normative motivation for two-factor learning rule

We have used in the previous models a so-called *two factor learning rule*, where the value updates depend only on the presynaptic activity (i.e., state input) and TD RPEs. Here, we motivate this choice from a normative approach based on previous work^11^.

Consider a linear approximation for value, where the value function 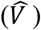 is the output of a single linear neuron. Here, 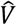 is a linear function of the input feature-vector representing the state **x**(*s*) = (*x*_1_(*s*),…, *x*_*n*_(*s*)), parametrized with a weight vector **w** = (***w***_1_,…, ***w***_*n*_).:

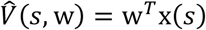

To put it into neural terms, we can think of *x*_*i*_(*s*) as the presynaptic activity onto the value neuron 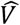, with a synaptic efficacy ***w***_*i*_.

As before, the agent computes the TD error based on this linear approximation for value and the sampled reward:

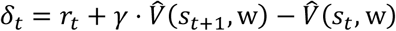

In the problem of *value prediction*, the agent aims to achieve the highest accuracy of prediction. One way to achieve this is to perform *stochastic gradient descent* (SGD) with respect to the parameters (w) of the value function to minimize the objective function such as the squared error 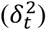. We can define this optimization problem as: 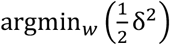 where we have deliberately chosen the constant 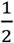 for clarity, but it doesn’t change the end results.

To perform SGD in this minimization problem, the parameters (w) should be updated in the opposite direction of the gradient of the loss with respect to the parameters (i.e., opposite to ∇_w_ (½ δ^2^)):

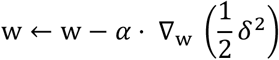

Where *α* is the learning rate. To compute the gradient, we use the chain rule:

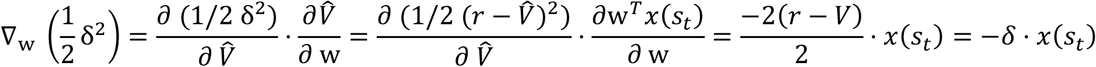

Therefore, the update for the parameters of the value function is:

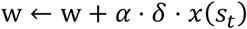

The term *δ* · x(*s*_*t*_) in the equation above is what we call a *two-factor learning rule*, dependent only on the presynaptic activity and not contingent on the post-synaptic activity.

The development of the TD learning model with D1 and D2 populations (section 1.3) has respected this learning rule, complying with what is required for SGD in the value prediction problem. Note that we have implicitly developed our models with a complete serial compound representation (CSC) of the states^11^, where x(*s*_*t*_) = 1 in a single element *x*_*i*_(*s*_*t*_)representing the current state and 0 otherwise. It can be shown that with this representation, the update equation above is equivalent to:

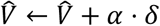

### 2. Computational model of dopamine release and receptor occupancy

To predict changes in dopamine concentrations and receptor occupancies (Fig. 6), we employed a biophysical model developed elsehwhere^59^. It presents two interacting dynamical systems. The first system models the change in receptor occupancies while the second the change in dopamine levels per unit time.

In the first system, the occupancy of receptors is modelled as a binding reaction between dopamine (*DA*) and D1 or D2 receptors (*R*), using the constants for forward and backward reactions (*k*_*on*_, *k*_*off*_).

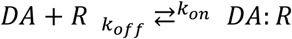

This formulation results in the following equation for the change in receptor occupancy *Occ*(*t*) per unit time:

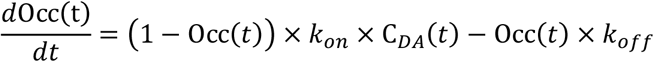

The values used for the association and dissociation constants for each receptor type (*k*_*on*_ and *k*_*off*_, respectively) are detailed in Table 1.

**Table 1.** Biophysical model parameters.

| Parameter | Abbreviation | Value |
| --- | --- | --- |
| DA association constant to D2 autorreceptors | $k_{on}^{D2term}$ | $0.3 M^{-1}s^{-1}$ |
| DA dissociation constant to D2 autorreceptors | $k_{off}^{D2term}$ | $0.003 s^{-1}$ |
| DA association constant to D1 receptors | $k_{on}^{D1}$ | $0.01 M^{-1}s^{-1}$ |
| DA dissociation constant to D1 receptors | $k_{off}^{D1}$ | $10 s^{-1}$ |
| DA association constant to D2 receptors | $k_{on}^{D2}$ | $0.2 M^{-1}s^{-1}$ |
| DA dissociation constant to D2 receptors | $k_{off}^{D2}$ | $2 s^{-1}$ |
| Release probability from terminals | $P_r$ | $1 a. u.$ |
| Release capacity from terminals | $\gamma_{pr_n}$ | $2 a. u.$ |
| D2 autorreceptor efficacy | $\alpha$ | $3 a. u.$ |
| DAT maximal uptake capacity | $V_{max}^{pr_n}$ | $1500 nMs^{-1}$ |
| Michaelis-Menten parameter DAT-mediated DA uptake | $K_m$ | $160 nM$ |
| Constant for dopamine removal not mediated by DAT's | $K_{nonDAT}$ | $0.04 nM$ |
| Bromocriptine association constant to D2 autorreceptors | $k_{on}^{Drug,D2s}$ | $0.02083 M^{-1}s^{-1}$ |
| Bromocriptine dissociation constant to D2 autorreceptors | $k_{off}^{Drug,D2s}$ | $0.1 s^{-1}$ |
| Bromocriptine association constant to D2 receptors | $k_{on}^{Drug,D2l}$ | $0.04 M^{-1}s^{-1}$ |
| Bromocriptine dissociation constant to D2 receptors | $k_{off}^{Drug,D2l}$ | $0.1 s^{-1}$ |

a.u. = arbitrary units
M = mols
s = seconds

In the second system, the change in dopamine concentration (*C*_*DA*_(*t*)) is a function of both dopamine release and uptake.

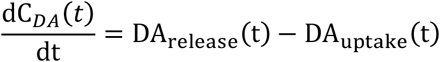

Dopamine release is a product of firing rate (*v*(*t*)) and release capacity (*γ*(*t*))

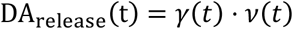

Where:

1. *v*(*t*) is the firing rate of dopamine neurons, provided by the neural data.
2. 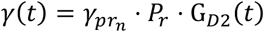 is defined as the increase in *C*_*DA*_(*t*) by a single synchronized action potential:
  a. *P*_*r*_ = 1 release probability in the absence of presynaptic D2-autorreceptors,
  b. 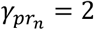 release capacity in the absence of presynaptic D2-autorreceptors. This value was set to be deliberately high and anticipates a ∼50% reduction by terminal feedback.
  c. G_*D*2_(*t*) is a multiplicative gain that represents the modulation of dopamine release by D2-autorreceptors. This is a decaying function of the occupancy of D2-autorreceptors 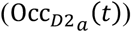 which is modelled by the same binding reaction explained above. The gain is parametrized by the autoreceptor efficacy, *α* = 3. The smaller the *α* the less the decay in release with receptor occupancy.

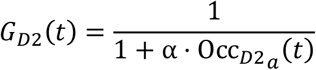

Dopamine uptake is a function of the uptake of dopamine by the dopamine transporter (DAT) and other non-DAT sources

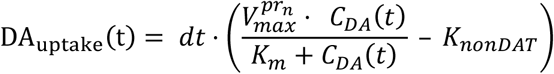

Where:

- 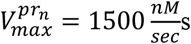 the maximal uptake capacity assuming approximately 100 terminals in the near surroundings.
- *K*_*m*_ = 160 *nM*, is the Michaelis-Menten parameter for uptake mediated by DAT
- *K*_*nonDAT*_ = 0.04 *nM* is a constant for the dopamine removal not mediated by DAT. For example, monoamine oxidase (MAO) and noepinephrine transporter (NET) mediated uptake.

The variables of the model reported in Fig. 6 correspond to: Occ_*D*1_(*t*), Occ_*D*2_(*t*), C_*DA*_(*t*). We used as input to the model the firing rates derived from the electrophysiological recording of optogenetically identified dopamine neurons conducted in Tian and Uchida (2015)^55^. This modeling, while considering major processes, does not take into account all of the complexity of the biological environment in the brain, yet we used this model to obtain an approximate estimate of the order of changes in dopamine concentrations and receptor occupancies.

## 3. Simulation details of habenula lesion data

### 3.1. Biophysical model simulations

We used the computational model described previously (methods section 2) ^59^ to calculate the extracellular dopamine levels and estimate the occupancy of postsynaptic receptors from the habenula lesion dataset. The model was driven by the average spike rate of dopamine neurons recorded from control or lesioned animals. For each recorded dopamine neuron, the simulations were carried on a trial by trial basis that consisted of a time window [-15, 20] sec with respect to cue onset. A relatively large window was used to allow for the relevant variables to stabilize in its baseline, as the simulations were initialized at zero.

For each trial, spikes were first binned with 10-ms windows and then smoothed by a Gaussian kernel (σ = 0.3 × (*ISI*_*mean*_)). All trials were then averaged across trials, to determine the mean single-cell response for dopamine release and D1 and D2 receptor activation. Final average dopamine concentrations and receptor occupancies were obtained from the average of all mean single-cell responses.

#### Computation of receptors sensitivities from the model results

We computed the receptor sensitivity from the occupancies Occ_*D*1_, Occ_*D*2_ and their theoretical dose-occupancy curves. Starting from the occupancy at baseline, we derived the change in occupancy as a function of the transients in dopamine concentration C_*DA*_ elicited by RPE-evoked dopamine responses, at the level of the population average.

The ratio between these quantities corresponds to the receptors’ sensitives. These are transferred as *α*^+^and *α*^−^ to our reinforcement learning model (model 1):

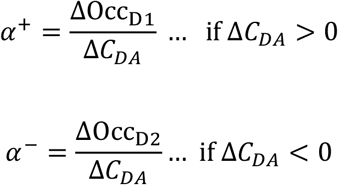

Where Δ*C*_*DA*_, Δ*Occ*_*D*1_, Δ*Occ*_*D*2_ are the changes computed with respect to baseline, as: 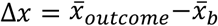, for each variable *x* = {*C*_*DA*_, Occ_*D*1_, Occ_*D*2_}. Where 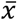 denotes the population average response for each group. The outcome responses were taken as the average from [0,1] sec after outcome onset, while the baseline was taken as the average from [-1, 0] sec with respect to cue onset.

### 3.2. Model 1 simulations

The simulations for Model 1 were carried out with a TD learning model with D1 and D2 populations (methods section 1.3). We ran the simulations using the resultant receptor sensitivities from the biophysical model as the population-level asymmetric learning rates in Model 1 (i.e., the learning rates *α*^+^, *α*^−^ for *P* and *N* updates). The simulations were run for 3,000 trials on the Pavlovian conditioning task used in the study^55^. We assumed a uniform distribution of trial types across the session. Each trial consisted of 4 states (baseline, cue, delay, reward), assuming Markovian dynamics between them. All variables were initialized at zero. The model had as hyper-parameters a discounting factor of *γ* = 0.99 and a decay term *β* = 0.002. We report in Fig. 4, Model 1 results assuming a uniform scaling of TD RPEs across the neuronal population. In Extended Data Fig. 12 we show that this model reproduces key signatures of the data irrespective of the choice of the decay factor β.

The results are not dependent on a uniform scaling of TD RPEs. Given that distributional RL signatures were preserved in the data even after habenula lesions, we also considered Model 1 under the distributional TD learning framework (Extended Data Fig. 13). For this, we used the distribution of single cell asymmetric scaling factors 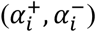 derived from the dopamine neurons firing rates. This model also reproduced key signatures of the data irrespective of the choice of the decay factor *β* (Extended Data Fig. 12).

### 3.3. Model 2 simulations

The simulations for Model 2 were carried out with a TD learning model. As with Model 1, simulations were run for 3,000 trials on the Pavlovian conditioning task^55^. We assumed a uniform distribution of trial types across the session. Each trial consisted of 4 states (baseline, cue, delay, reward), assuming Markovian dynamics between them. All variables were initialized at zero. The model had as parameters a discounting factor of *γ* = 0.99.

We used the distribution of single cell asymmetric scaling factors derived from the firing rates of dopamine neurons as 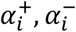. In section 1.2 we emphasized that in order to accurately compute the TD RPE in distributional TD, we require taking samples from the estimated return distribution 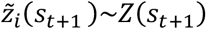. We did this by running an optimization process where we minimize for the expectile loss between the taken samples 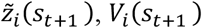 from the model, and *τ*_*i*_ as estimated from the data. The problem was defined as argmin 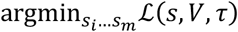 where:

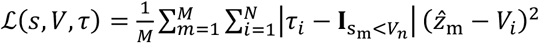, for *N* neurons and *M* samples

In the simulations, we took *M* samples where *M* equals the number of neurons (*N*) and performed an update taking the expectation across all samples as described in the methods section 1.3.

## 4. Simulations details for replications of previous experimental results

### 4.1. Cools et al. (2009)

We simulate the results from Cools et al. (2009) (Fig. 7, Extended Data Fig. 6-7) in which they tested the effects of bromocriptine in altering learning rate asymmetry^24^. In their study, they performed a reversal learning task and reported a parameter called ‘relative reversal learning (RRL)’, equivalent to the difference between the positive and negative learning rates in our model. We computed this as: *α*^+^*α* + +*α*— *α* − *α* + +*α*−= *τ* − 1 − *τ* = 2*τ* − 1, reported in Fig. 7 E,F, where the parameters *α*^+^, *α*^−^ were computed from the slopes of the D2l (postsynaptic D2 receptors) and D1 occupancy curves (2*τ* − 1)_*occ*_ or activation curves (2*τ* − 1)_*act*_ The change in relative reversal learning in Fig. 7 H-I was calculated as taking the difference between drug and the ‘control’ condition as:

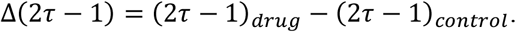

We simulated the effect of bromocriptine using the biophysical model for dopamine release and receptor occupancy (Section 2, Methods). We added an additional ligand for D2 receptors to the update equations for occupancy:

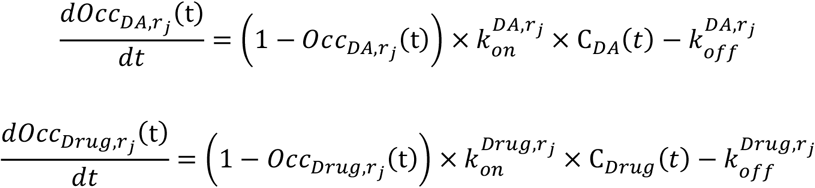

Where 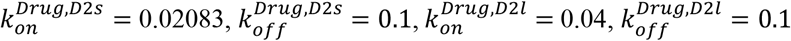 are reported in Table 1 ^97^.

To calculate the effects of efficiency of the drug, we calculated the activation of D2l and D2s receptors in the following way:

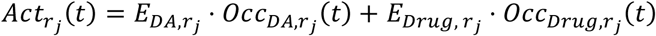

Where 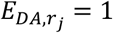 is the efficiency of dopamine on the receptors activation, and 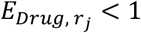 the efficiency of the drug, for *r*_*j*_: {*D*1, *D*2*s, D*2*l*}. The parameter for D1 receptors was kept at *E*_*Drug,D*1_ = 0 for all simulations.

To simulate the effects of D2s activation by the drug in D2l occupancy in Fig. 7b,e,h we report the effects of *E*_*Drug,D*2*s*_ = 0 (solid lines) and *E*_*Drug,D*2*s*_ = 0.6 (dashed lines). To simulate the effect of the drug in D2s and D2l activation in Fig. 7c,f,i we report the effects of *E*_*Drug,D*2*s*_ = 0.6, *E*_*Drug,D*2*l*_ = 0.6.

We show how the qualitative nature of the effects of the drug in relative reversal learning still hold regardless of whether the parameter *τ* is computed from the occupancy curves (Extended Data Fig. 7, Fig. 7n,e,h) or the activation curves (Extended Data Fig. 8, Fig. 7c,f,i). In addition, in Supplementary Figure

8-9 we show that the qualitative results still hold regardless of the choice of the efficiency parameters *E*_*Drug,D*2*s*_ and *E*_*Drug,D*2*l*_.

### 4.2. Timmer et al. (2018)

In this study^25^ they reported a ‘loss aversion’ parameter (λ in their results).

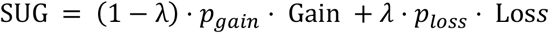

Where SUG is the ‘subjective utility’ for a given option, and *p*_*gain*_ = *p*_*loss*_.

In our formulation, we assume that the task in the study is performed under steady state conditions after having learned with a learning rate (*τ*). With this assumption, the SUG at task performance is equivalent to the convergent *V* estimate after learning. We will show that at these steady state conditions (1 − *τ*) is equivalent to (λ).

Starting with the solution for *V*:

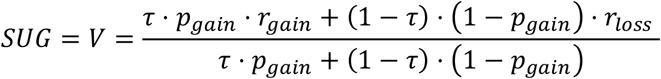

Replacing for *p*_*gain*_ = 0.5:

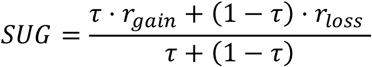

Given that: *τ* + (1 − *τ*) = 1

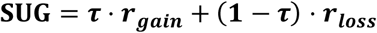

Therefore, our model, applied to their task, gives rise to the same SUG computation, with λ equivalent to (**1** − **τ**).

To generate Fig. 8F, we performed the following steps:

1. We first estimated the theoretical change in baseline DA elicited by the medication. For this, we computed the equivalent *τ* for the λ they report in the OFF and ON medication conditions (λ_*OFF*_ = 1.51, λ_*ON*_ = 1.19), using the relationship: (1 − *τ*) = λ. We then computed the baseline DA levels that would give rise to the *τ*_*ON*_ and *τ*_*OFF*_. With this, we computed the change in baseline DA (Δ*DA*) equivalent to the change Δ*τ* = *τ*_*ON*_ − *τ*_*OFF*_. This Δ*DA* is the theoretical change in baseline DA elicited by the medication (Fig. 8F).
2. To generate Fig. 8F, we sampled a set of λ from a Gaussian distribution centered at a mean of *μ*_λ_ = 1.51 and a standard deviation of 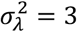, to emulate the distribution of λ_*OFF*_ they report in the OFF condition. We then computed the equivalent *τ* for that set of λ with the relationship above. We will call this the distribution of *τ*^′^_*OFF*_.
3. We used the derived *τ*^′^_*OFF*_ distribution to compute the equivalent dopamine levels. We imposed a change in baseline DA equal to the Δ*DA* computed in the first step and computed the new set of *τ* for that set of new baseline DA levels (*τ*^′^_*ON*_). The ‘drug effect in loss aversion’ reported in Fig. 8F is the *τ*^′^_*ON*_ − *τ*^′^_*OFF*_ for each sample.

## 5. Details on habenula lesion data

### 5.1. Animals, surgery and lesions

The rodent data we re-analyzed here were first reported in Tian and Uchida (2015)^55^. Below we provide a brief description of the methods. Further methodological details can be found in the original paper. !2 mice were used. Bilateral habenula lesions were performed in five animals. Seven animals were in the control group including two with sham-lesion operation, one with only small contra-lateral side lesion of the medial habenula, and four animals without operations in the habenula. During surgery, a head plate was implanted on the skull, and adeno-associated virus (AAV) that express channelrhodopsin-2 (ChR2) in a Cre-dependent manner was injected into the VTA (from bregma: 3.1 mm posterior, 0.7 mm lateral, 4–4.2 mm ventral). After recovery from surgery, mice were trained on the conditioning task, after which mice were randomly selected to be in lesion or sham-lesion group. Electrolytic lesions were made bilaterally using a stainless-steel electrode (15 kU, MicroProbes, MS301G) with a cathodal current of 150 mA. Each side of the brain was lesioned at two locations (from bregma: 1.6 mm/1.9 mm posterior, 1.15 mm lateral, 2.93 mm depth, with a 14 angle). For sham-lesion operations, no current was applied. In the same surgery, a microdrive containing electrodes and an optical fiber was implanted in the VTA (from bregma: 3.1 mm posterior, 0.7 mm lateral, 3.8–4.0 mm ventral)^98^.

### 5.2. Behavioral task

Twelve mice were trained on a probabilistic Pavlovian task. Each trial the animal experienced one of four odor cues for 1 s, followed by a 1-s pause, followed by a reward (3.75 μl water), an aversive air puff or nothing. Odor 1 to 3 signaled a 90%, 50% and 10% probability of reward, respectively. Odor 4 signaled a 90% probability of air puff. Odor identities were randomized across trials and included: isoamyl acetate, eugenol, 1-hexanol, p-cymene, ethyl butyrate, 1-butanol, and carvone (1/10 dilution in paraffin oil). Intertrial intervals were exponentially distributed. An infrared beam was positioned in front of the water delivery spout and each beam break was recorded as one lick event. We report the average lick rate over the interval 500–2,000 ms after cue onset.

### 5.3. Electrophysiology

Recordings were made using a custom-built microdrive equipped with 200-μm-fiber optic-coupled with eight tetrodes. DA neurons were identified optogenetically^98^. A stimulus-associated spike latency test (SALT) algorithm^99^ was used to determine whether light pulses significantly changed a neuron’s spike timing.

### 5.4. Neural data analysis

Data analyses were performed using MATLAB R2021b (Mathworks). To measure firing rates, peristimulus time histograms (PSTHs) were constructed using 1-ms bins. These histograms were then smoothed by convolving with the function 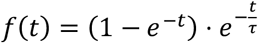. where *τ* was a time constant set to 20 ms as in ^18^. 44 dopamine neurons were recorded from lesioned animals (5 animals, 30 sessions), and 45 dopamine neurons were recorded from control animals (7 animals, 35 sessions). We pooled all the cells across animals in each group for analysis. Cue-evoked responses were defined as the average activity from 0 to 400 ms after cue onset. Outcome-evoked responses were defined as the average activity from 2000 to 2600 ms after cue onset.

The normalization of cue response shown in Fig. 4 was carried out following a previous work^36^ on a per-cell basis as: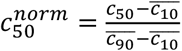, where 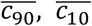 correspond to the mean across trials within a cell for the 90% and 10% probability cure responses. To derive the t-statistics in Fig. 4d, we performed a two-tailed t-test of the cell’s normalized responses to the 50% cue against the average midway point between responses to the 10% cue and responses to the 90% cue.

The derivation of asymmetric scaling factors from outcome responses (*τ*_*i*_), was carried out following ^36^, with some modifications to adapt it to the task. The procedure is illustrated in Extended Data Fig. 3.

- To compute the reversal points, outcome responses were first aligned to the RPE for each trial type, computed with the true expected value of each reward distribution. Assuming a fixed reward value of 1 (arbitrary units), the expected value for the 90%, 50%, 10% reward probability trials corresponded to 0.1, 0.5, 0.9, respectively. Given this, omission responses from the 90%, 50%, 10% reward probability trials correspond to RPEs of -0.9, -0.5 and -0.1. The rewarded responses from the 90%, 50%, 10% reward probability trials correspond to RPEs of 0.1, 0.5 and 0.9. The reward value is arbitrary and doesn’t have an effect in this computation as it only shifts the RPE axis by a fixed amount. The reversal point for each cell (*Z*_*i*_) was defined as the RPE that maximized the number of positive responses to RPEs greater than *Z*_*i*_ plus the number of negative responses to RPEs less than *Z*_*i*_. The distribution of reversal points is reported in Extended Data Fig. 4. To obtain statistics for reliability of the computed reversal points, we partitioned the data into random halves and estimated the reversal point for each cell separately in each half. We repeated this procedure 1000 times with different random partitions, and we report the distribution of Pearson’s correlation across these 1000 folds (Extended Data Fig. 4).
- After measuring reversal points, we fit linear functions separately to the positive and negative domains. Given that dopamine’s responses are non-linear in the reward space but present a putative utility function^100^, we approximated the underlying utility function from the dopamine responses to RPEs of varying magnitudes. We used these empirical utilities instead of raw RPEs for computing the slopes that correspond to 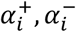. We then computed the asymmetric scaling factors as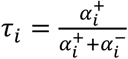. We performed the same cross-validation procedure used for the reversal points. The distribution of R value across the 1000 folds are reported in Extended Data Fig. 4.

A key prediction of distributional RL^36^ is the presence of a correlation (across cells) between reversal points *Z*_*i*_ and asymmetric scaling factors *τ*_*i*_. To elucidate whether signatures of distributional RL were still present after lesions, we followed the procedure given by Dabney et al. (2020)^36^ to compute this correlation. We first randomly split the data into two disjoint halves of trials. In one half, we first calculated reversal points 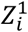 and used them to calculate 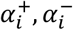. In the other half, we again calculated the reversal points 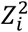. The correlation we report in Extended Data Fig. 4 is between 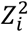 and 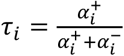.

### 5.5. Model fitting to the anticipatory licking responses

For each trial we computed the average lick rate over the interval 500–2,000 ms after cue onset. For each model, we fit the free parameters to the lick rates using maximum likelihood estimation. The optimization was performed using the SciPy optimization toolbox (Python) that minimized the difference between the predicted lick rates and the ground truth ones, with a uniform prior distribution over the parameters. The fits were done considering three RL models that had between 2 and 3 parameters. The models, parameters and bounds used for each of them are detailed in table 2.

**Table 2.** Reinforcement learning models fit to the behavioral data from Tian & Uchida.

| Model | Formulation | Parameters | Parameter bounds |
| --- | --- | --- | --- |
| TD learning | $\delta = r - V$ $V \leftarrow V + \alpha \cdot \delta$ $Licking = \beta \cdot V$ | $\alpha, \beta$ | $\alpha \in [.001, 1]$<br>$\beta \in [.1, 10]$ |
| TD learning with reward sensitivity | $\delta = \rho \cdot r - V$ $V \leftarrow V + \alpha \cdot \delta$ $Licking = \beta \cdot V$ | $\alpha, \rho, \beta$ | $\alpha \in [.001, 1]$<br>$\rho \in [.001, 10]$<br>$\beta \in [.1, 10]$ |
| Risk sensitive TD learning | $\delta = r - V$ $V \leftarrow V + \alpha^+ \cdot \delta \quad \text{if } \delta > 0$ $V \leftarrow V + \alpha^- \cdot \delta \quad \text{if } \delta < 0$ $Licking = \beta \cdot V$ | $\alpha^+, \alpha^-, \beta$ | $\alpha^+ \in [.001, 1]$<br>$\alpha^- \in [.001, 1]$<br>$\beta \in [.1, 10]$ |

## Data availability

The neural data and simulation results reported in this article have been shared in a public deposit source in: https://osf.io/cr5mv/?view_only=bd13a2d2de1947699b56ce70610b0e9b

## Code availability

The accession codes for the data as well as the code for analysis and simulations are available at: https://github.com/sandraromerop/D1D2_Dopamine

## Acknowledgements

We thank Ju Tian for providing the data used in this study; Jakob Dreyer for making the biophysical model available for open-source use; Roshan Cools, Peter Dayan and Sam Gershman for valuable feedback on the manuscript; Dan Polley, Mark Andermann and Elizabeth Phelps for mentorship and helpful discussions. We thank Bernardo Sabatini for discussions, and Isobel Green, Paul Masset, Mitsuko Watabe-Uchida and all the rest of the Uchida lab members for helpful comments at the different stages of this work. This work is supported by NIH BRAIN Initiative grants (R01NS116753, U19NS113201), the Simons Collaboration on Global Brain, and the Bipolar Disorder Seed Grant from the Harvard Brain Science Institute.

## Author information

### Contributions

S.R.P. and N.U. conceived the project. S.R.P performed the modeling work. S.R.P. wrote the first draft and S.R.P and N.U. edited the paper.

## Extended data Figures

**Extended Data Fig. 1.**
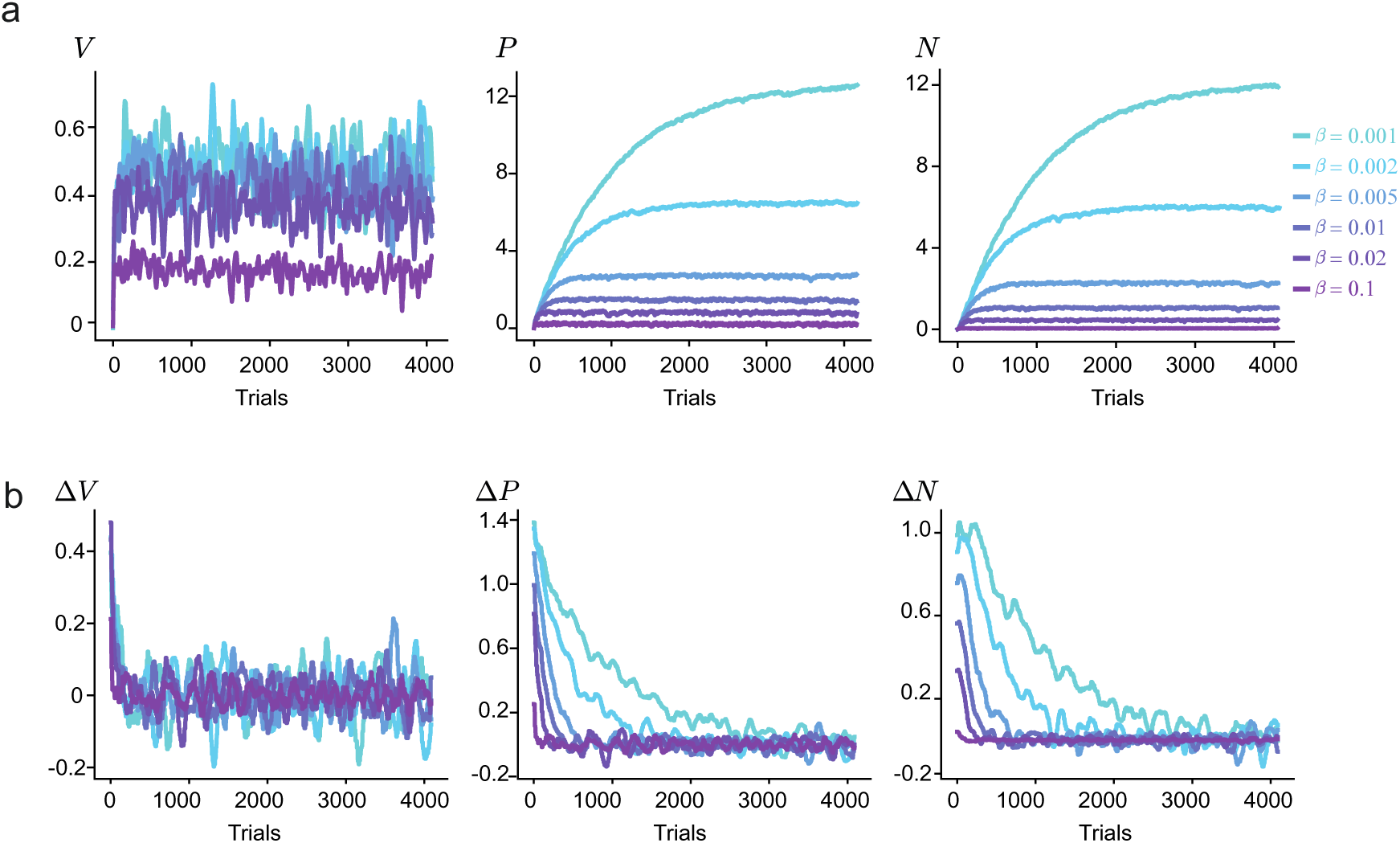
Variables of model 1 show convergence irrespective of the value of the decay factor.a. Value predictors *V* (left), *P* population (middle) and *N* population (right) across trials of training for an RL agent of model 1. Color of lines denotes the value of the decay factor (β) in the update rules for the *P* and *N* populations. Colormap is the same for all figures (left). All the model variables show convergence for every value of the decay factor β. **b**.Difference in the variables estimates between consecutive trials of training, for the value predictors *V* (left, ΔV), *P* population (middle, ΔP) and *N* population (right, ΔN). All the variables show convergence for every value of the decay factor *β* (shown as a ΔV, ΔP, ΔN equal to zero).

**Extended Data Fig. 2.**
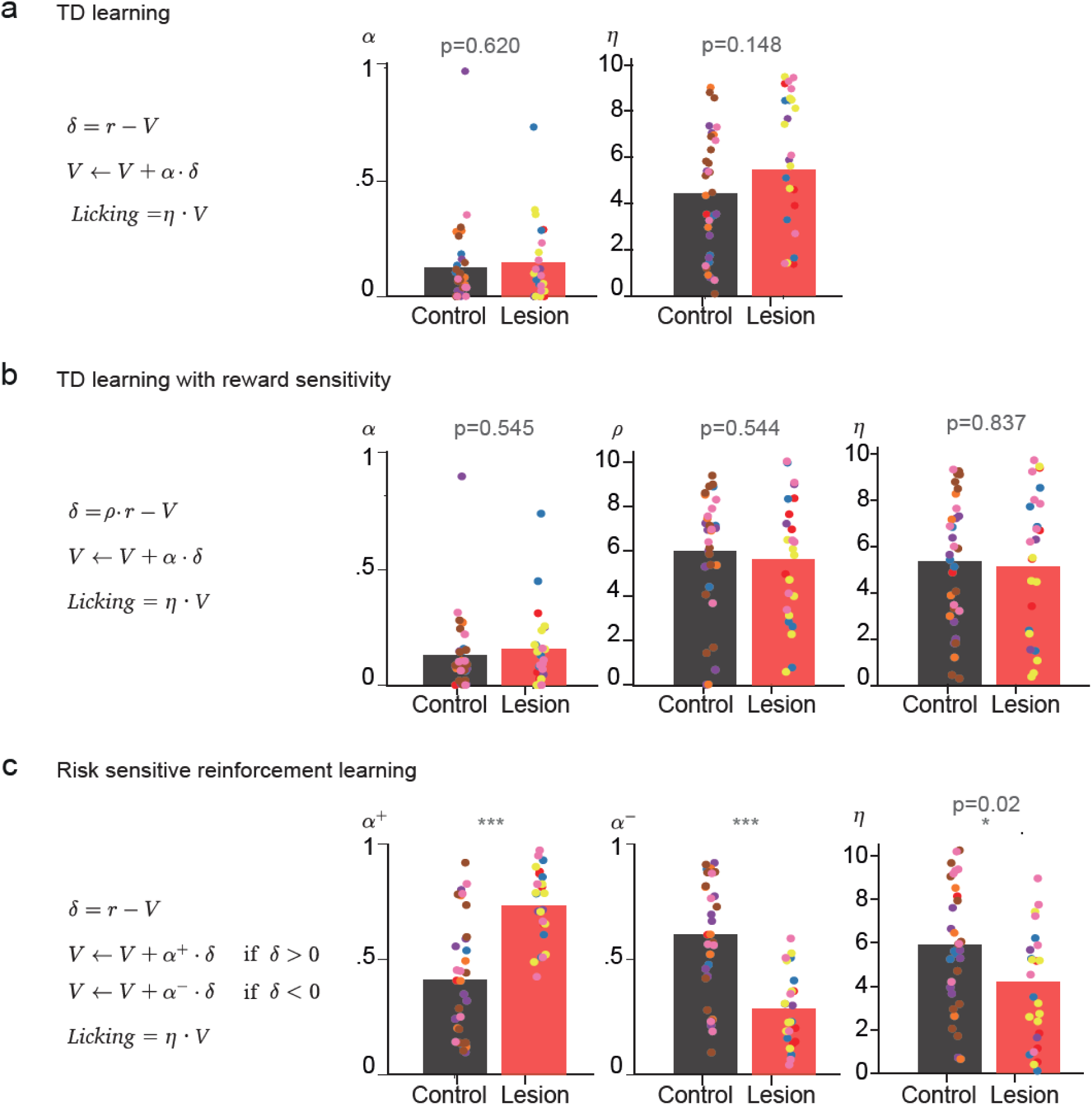
RL model fits to the trial-by-trial anticipatory licking responses. **a**. TD learning fits reveal no significant difference across groups in the learning rate (left, U-statistic = –4.954, P =0620, two-sided Mann-Whitney U-test) nor in the linear mapping between value predictions and anticipatory licking responses (U-statistic = –1.445, *P* = 0.148, two-sided Mann-Whitney U-test). **b**.Model fits of TD learning with reward sensitivity reveal no difference across groups in the learning rate (left, U-statistic = 0.206, *P* = 0.836, two-sided Mann-Whitney U-test) nor in the linear mapping between value predictions and anticipatory licking responses (middle, U-statistic = –0.7844, *P* = 0.4327, two-sided Mann-Whitney U-test), nor in the reward sensitivity (right) (U-statistic 0.545, *P* = 0.605, two-sided Mann-Whitney U-test). **c**.Model fits of TD learning with asymmetric learning rates for positive vs negative RPEs. This model reveals a significant difference across groups in the asymmetry between *α*^+^ and *α*^−^ (U-statistic = –4.678, *P* < 1.0 x 10^−5^, two-sided Mann-Whitney U-test) and a small but significant difference between the linear mapping between value predictions and anticipatory licking responses (right, U-statistic = 2.33, *P* = 0.02, two-sided Mann-Whitney U-test).

**Extended Data Fig. 3.**
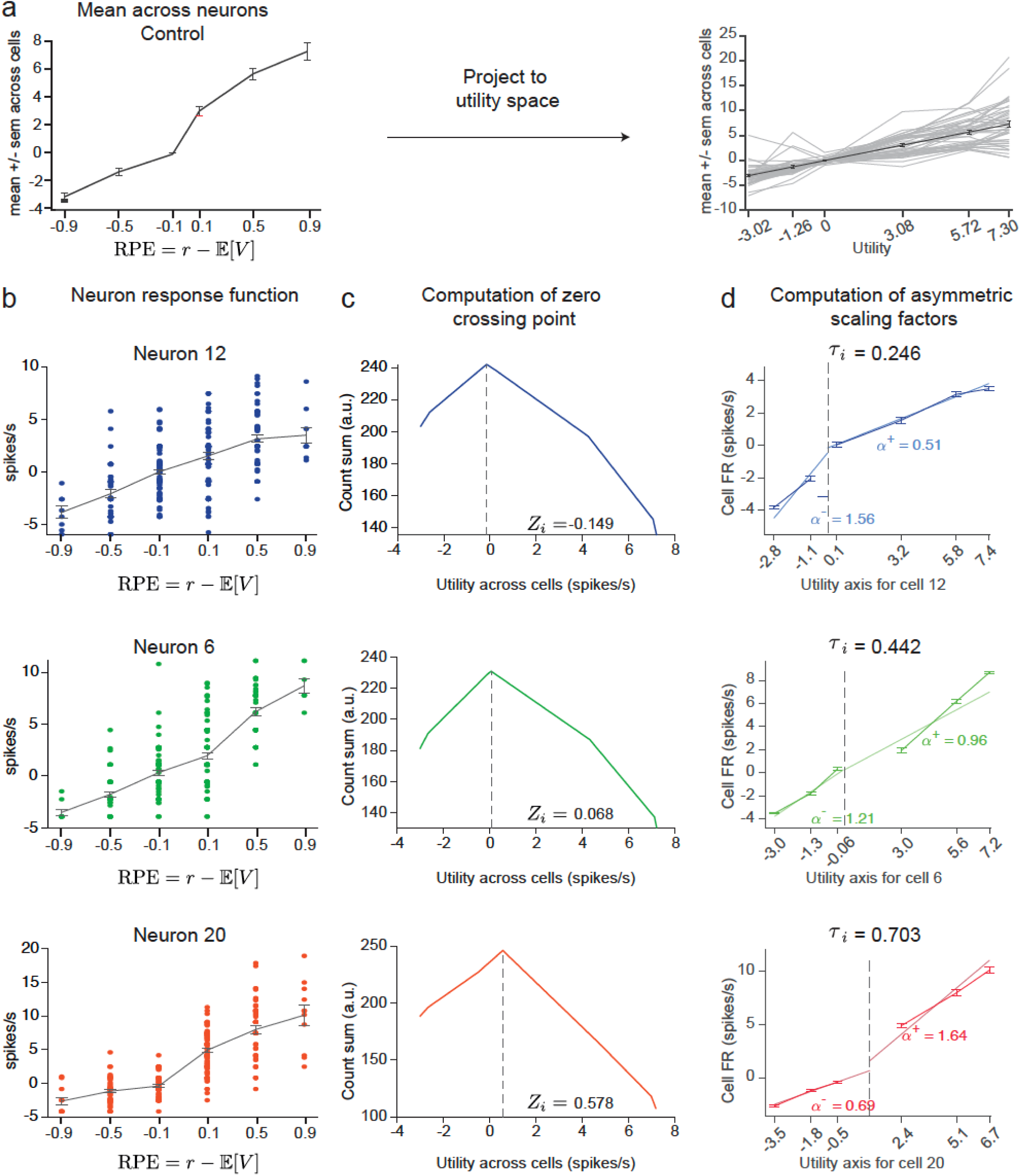
Signatures of distributional reinforcement learning model are preserved after habenula lesions. **a**. RPE -evoked responses at outcome as a function of the theoretical RPE for each trial type. The figure shows the average response function across neurons from the control group. The computation of zero-crossing points and asymmetric scaling factors is carried out in the ‘utility space’(the average response function) as in ^36^ to account for response nonlinearities. **b**. Example of the response function of 3 dopamine neurons from the control group ordered by their asymmetric scaling factors: pessimistic, neutral and optimistic, from top to bottom. **c**. Computation of zero-crossing points for the neurons in B. The reversal points for each cell (*Z*_*i*_) were defined as the point in utility space that maximized the number of positive responses to points greater than *Z*_*i*_ plus the number of negative responses to points less than *Z*_*i*_. The y-axis shows the sum of responses below and above each point in the utility space. The zero-crossing point is shown as the maxima in this function. **d**. Computation of asymmetric scaling factors for the neurons in c. Here, the responses functions in b have been projected to the utility space in A and realigned according to their zero-crossing points. The asymmetric learning rates (*α*^+^, *α*^−^) are taken to be the slopes of these response functions.

**Extended Data Fig. 4.**
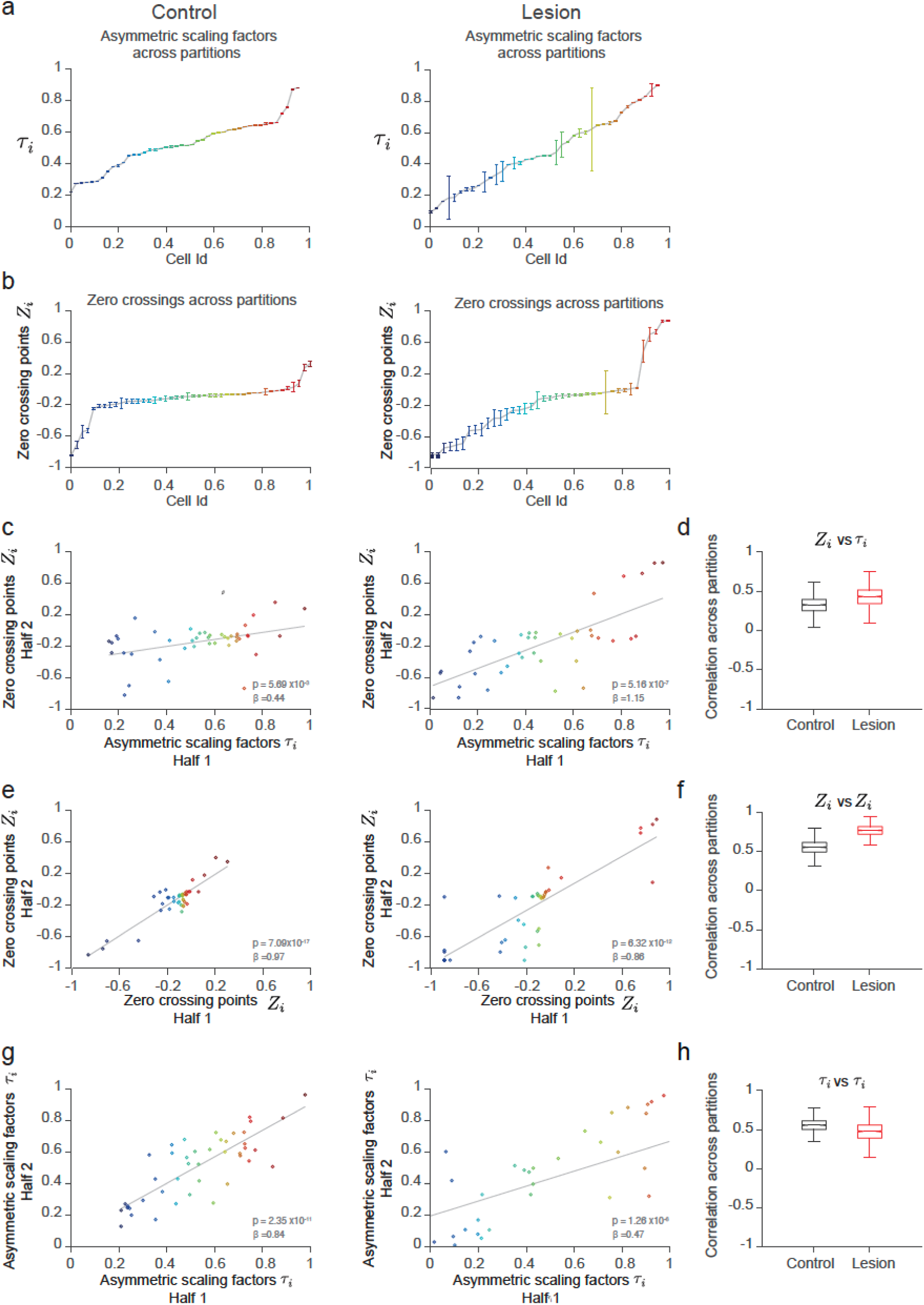
Distributional reinforcement learning variables from the Habenula lesion dataset. **a**.Distribution of asymmetric scaling factors for the dopamine neurons from the control (left) and lesion (right) groups. The error bars were derived by randomly sampling trials to compute the asymmetric scaling factors for 1000 iterations. **b**. Distribution of zero crossing points for the control (left) and lesion(right) groups. The error bars were derived as in a. **c**. Correlation of asymmetric scaling factors (x-axis) and zero-crossing points (y-axis) computed on disjoint halves of trials for an example partition. **d**. Distribution of correlation coefficients between asymmetric scaling factors (x-axis) and zero-crossing points (y-axis) across disjoint halves of trials for 1000 partitions for the control and lesion groups. **e**. Correlation between zero-crossing points computed on disjoint halves of trials for an example partition. **f**. Distribution of correlation coefficients between zero-crossing points computed on disjoint halves of trials for 1000 partitions for the control and lesion groups. **g**. Correlation between asymmetric scaling factors computed on disjoint halves of trials for an example partition. **h**. Distribution of correlation coefficients between asymmetric scaling factors computed on disjoint halves of trials for 1000 partitions for the control and lesion groups.

**Extended Data Fig. 5.**
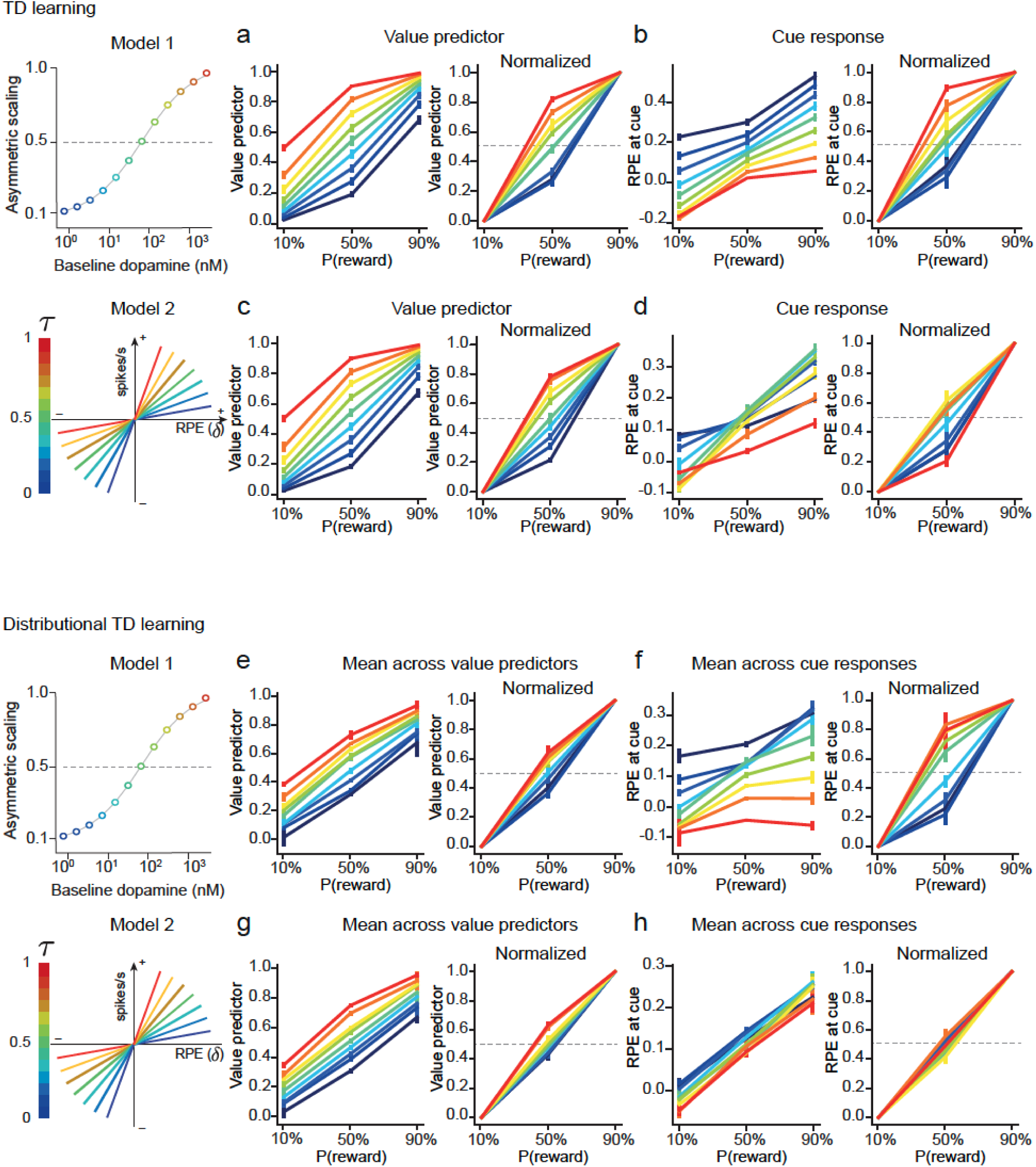
Biases in cue-evoked responses in the Habenula lesion data cannot be explained by asymmetric scaling of RPEs (Model 2). **a**. Value predictors derived from model 1 with TD learning for a set of baseline dopamine levels (colormap). The optimistic and pessimistic biases are present. **b**. Cue responses derived from model 1 with TD learning for a set of baseline dopamine levels (colormap). The optimistic and pessimistic biases are revealed when the responses are normalized. **c**. Value predictors derived from model 2 with TD learning for a set of baseline dopamine levels (colormap). The optimistic and pessimistic biases are present. **d**. Cue responses derived from model 2 with TD learning for a set of baseline dopamine levels (colormap). The optimistic and pessimistic biases are absent in both the normalized and the raw TD errors. **e**. Mean across the distribution of value predictors derived from model 1 with distributional TD learning for a set of baseline dopamine levels (colormap). The optimistic and pessimistic biases are present. **f**. Mean across the distribution of cue responses derived from model 1 with distributional TD learning for a set of baseline dopamine levels (colormap). The optimistic and pessimistic biases are revealed when the responses are normalized. **g**. Mean across the distribution of value predictors derived from model 2 with distributional TD learning for a set of baseline dopamine levels (colormap). The optimistic and pessimistic biases are present. **h**. Mean across the distribution of cue responses derived from model 2 with distributional TD learning for a set of baseline dopamine levels (colormap). The optimistic and pessimistic biases are absent in both the normalized and the raw TD errors.

**Extended Data Fig. 6.**
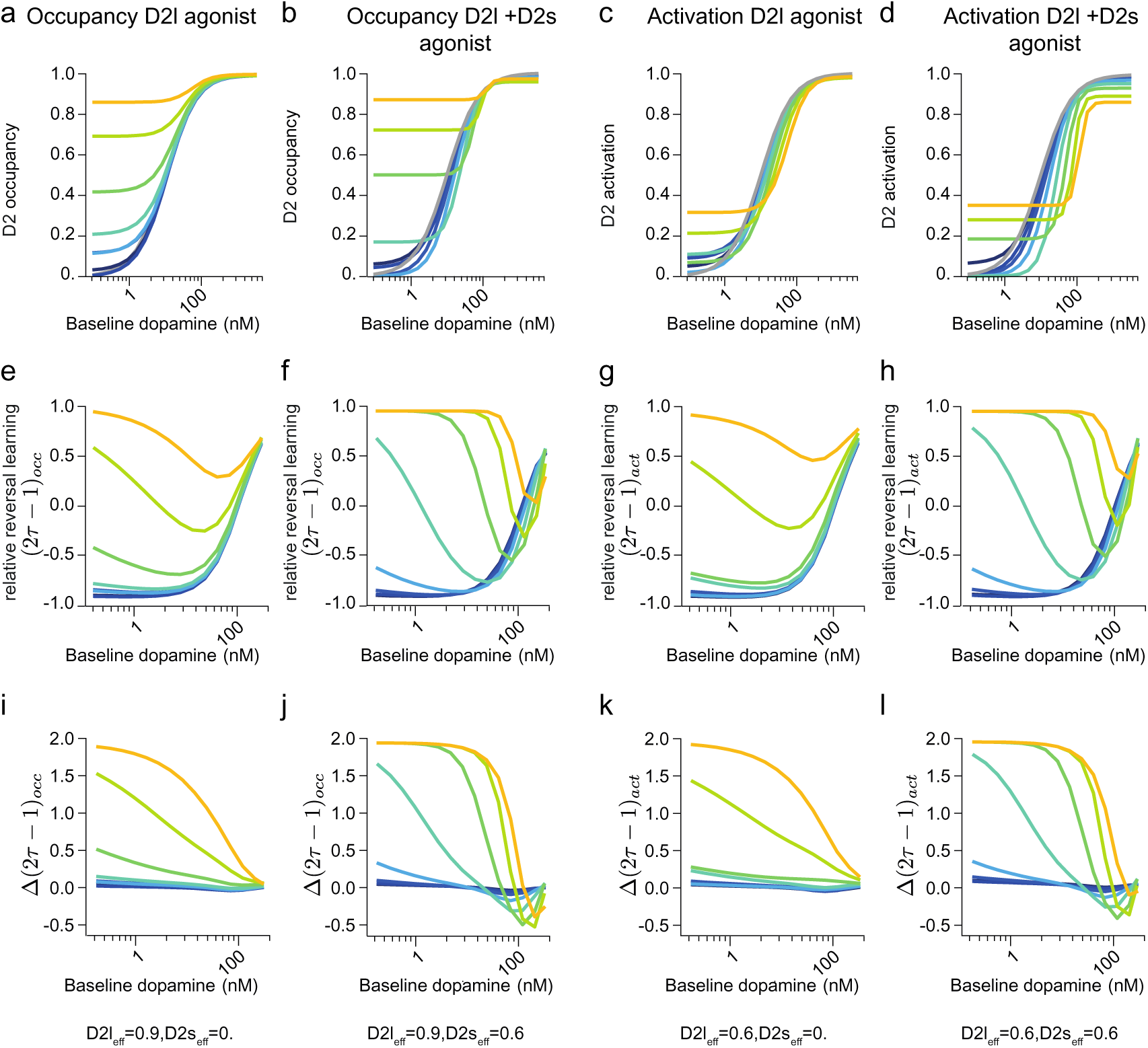
Model 1 predicts asymmetric learning rates and the effect of bromocriptine in healthy humans given inter-individual differences in baseline dopamine. **a**. Occupancy curves for the D2l receptors at baseline (grey line) and when considering bromocriptine’s effects in D2l receptors alone. The binding of the drug to the D2l receptors alone causes an increase in the occupancy. This happens to a larger extent when starting from a low dopamine level at baseline than in high dopamine levels. **b**. Occupancy curves for the D2l receptors at baseline (grey line) and when considering bromocriptine’s effects in both D2l and D2s receptors. The binding of the drug to D2s receptors causes a rightwards shifts in the curves. **c**. Activation curves for the D2l receptors at baseline (grey line) and when considering bromocriptine’s effects in D2l receptors alone, including the *partial* quality of the agonism of this drug on the receptor (i.e., efficiency <1). **d**. Activation curves for the D2l receptors at baseline (grey line) and when considering bromocriptine’s effects in both D2l and D2s receptors, including the *partial* quality of the agonism. **e**. Relative reversal learning (RRL) calculated as 2*τ* − 1 in model 1, as a function of baseline dopamine (x-axis) and drug concentration (color) using the receptors occupancy curve, considering only bromocriptine’s effect in D2l receptors. **f**. Relative reversal learning (RRL) calculated as 2*τ* − 1 in model 1, as a function of baseline dopamine (x-axis) and drug concentration (color) using the receptors occupancy curve, considering bromocriptine’s effect in both D2l and D2s receptors. **g**.2*τ* − 1 in model 1, as a function of baseline dopamine (x-axis) and drug concentration (color) using the receptors activation curve, considering only bromocriptine’s effect in D2l receptors. **h**.2*τ* − 1 in model 1, as a function of baseline dopamine (x-axis) and drug concentration (color) using the receptors activation curve, considering bromocriptine’s effect in both D2l and D2s receptors. **i**. The change in 2*τ* − 1 induced by the drug at different concentrations (color) with respect to the baseline condition, as a function of baseline dopamine (x-axis). The curves represent the change when calculating 2*τ* − 1 from the occupancy curves considering only D2l binding. **j**. The change in 2*τ* − 1 induced by the drug at different concentrations (color) with respect to the baseline condition, as a function of baseline dopamine (x-axis). The curves represent the change when calculating 2*τ* − 1 from the occupancy curves considering both D2l and D2s binding **k**. The change in 2*τ* − 1 induced by the drug at different concentrations (color) with respect to the baseline condition, as a function of baseline dopamine (x-axis). The curves represent the change when calculating 2*τ* − 1 from the activation curves considering only D2l activation. **l**. The change in 2*τ* − 1 induced by the drug at different concentrations (color) with respect to the baseline condition, as a function of baseline dopamine (x-axis). The curves represent the change when calculating 2*τ* − 1 from the activation curves considering both D2l and D2s activation. The parameters of efficiency of activation of D2 receptors by the drug (*D*2*l*_*eff*_, *D*2*s*_*eff*_) used in each column are reported at the bottom of the figure.

**Extended Data Fig. 7.**
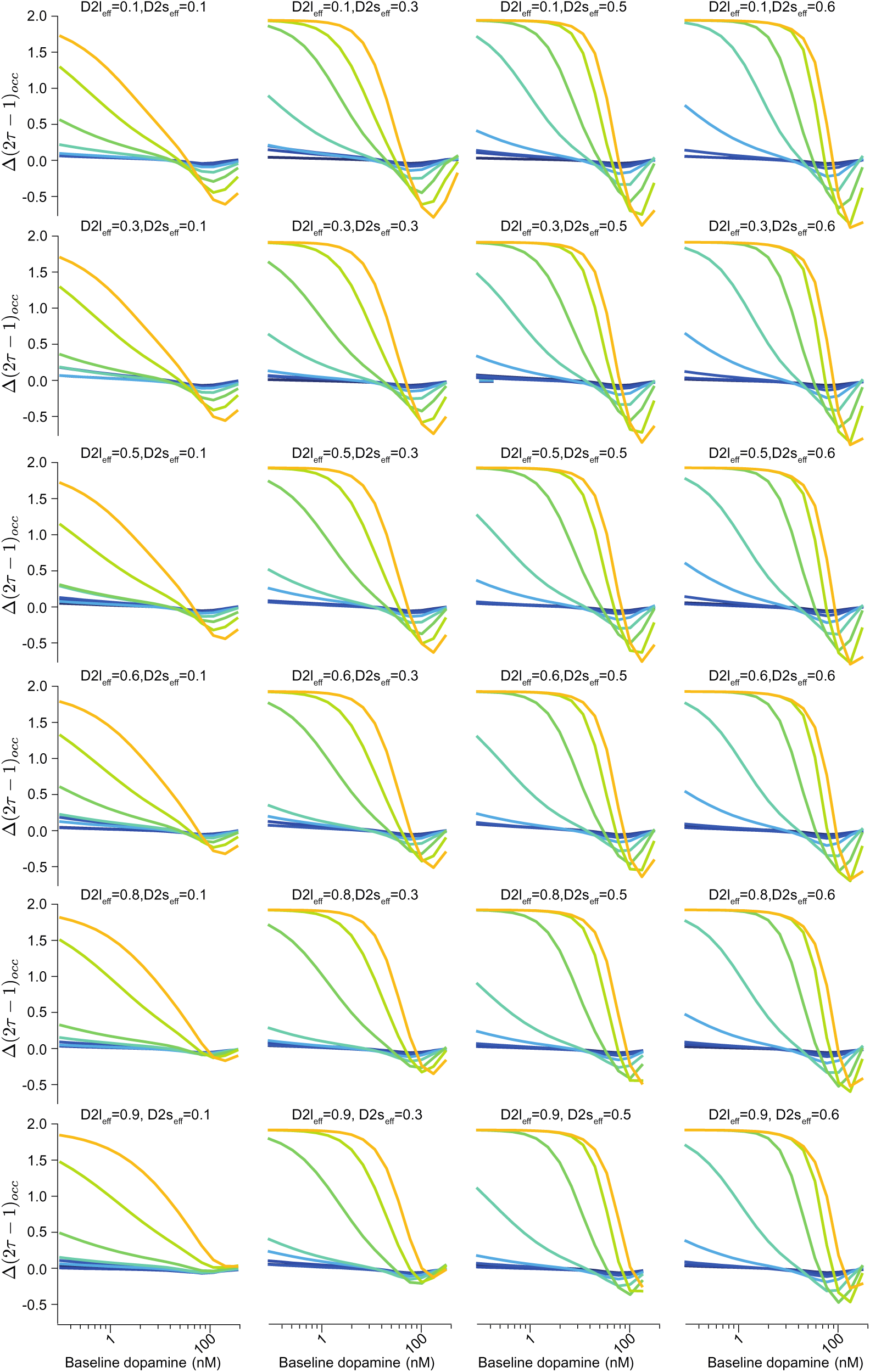
Robustness of the effect of bromocriptine in the relative reversal learning calculated from the occupancy curves to the choice of the drug efficiency parameter. The qualitative effects on bromocriptine in the change in relative reversal learning Δ(2*τ* − 1) calculated from the D2 *occupancy* curves. Results hold regardless of the choice of the efficiency of the drug on D2l (rows) or D2s (columns) efficiency.

**Extended Data Fig. 8.**
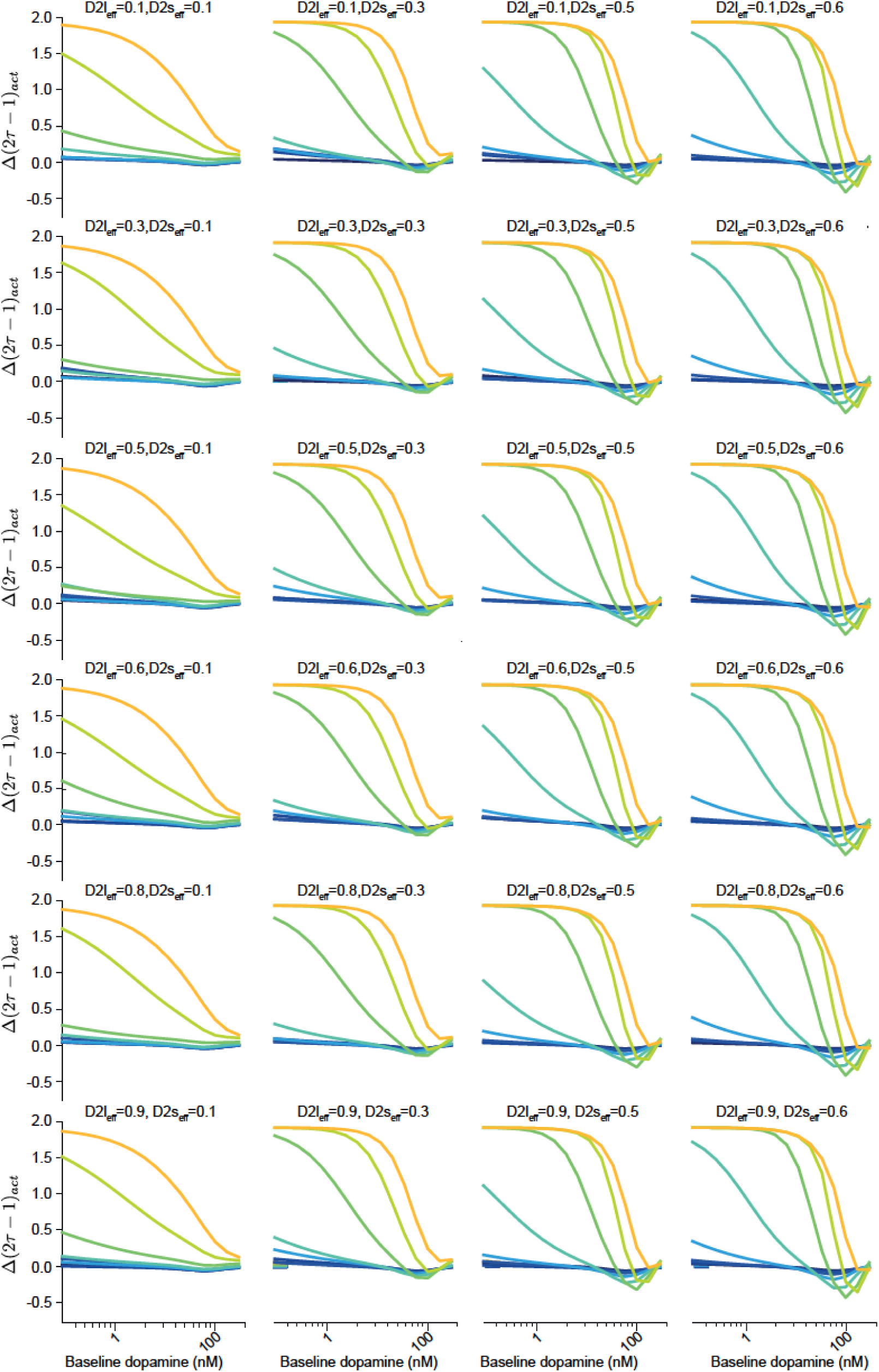
Robustness of the effect of bromocriptine in the relative reversal learning calculated from the activation curves to the choice of the drug efficiency parameter. The qualitative effects on bromocriptine in the change in relative reversal learning Δ(2*τ* − 1) calculated from the D2 *activation* curves. Results hold regardless of the choice of the efficiency of the drug on D2l (rows) or D2s (columns) efficiency.

**Extended Data Fig. 9.**
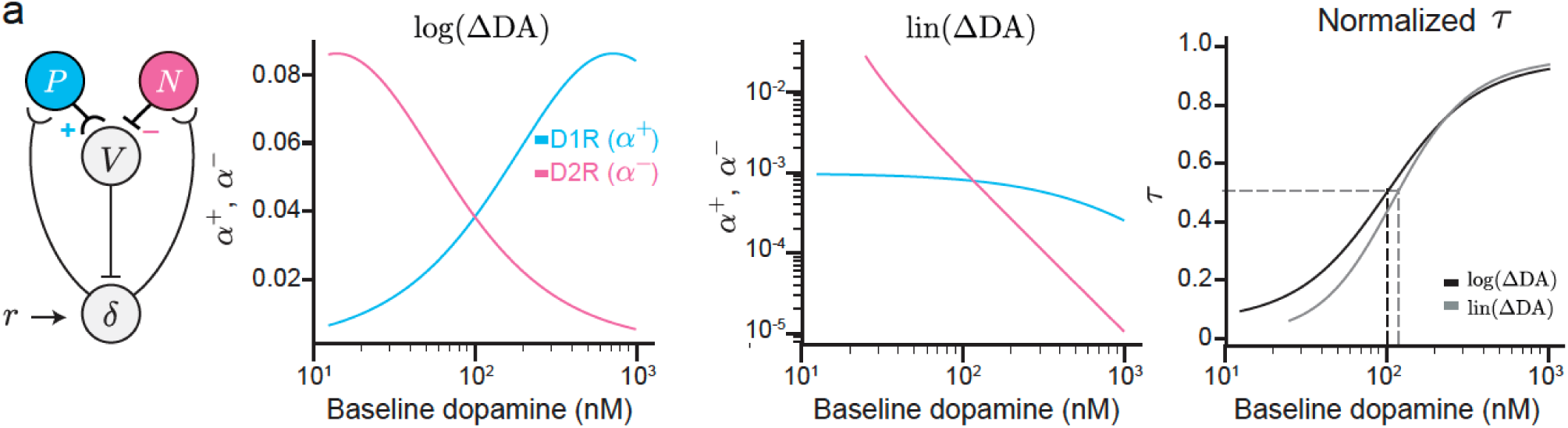
The qualitative aspects of Model 1 are preserved irrespective of the assumption made about the changes in baseline dopamine caused by dopamine transients. **a**. Computation of receptor sensitivities (i.e., slope of dose-occupancy curves, *α*^+^, *α*^−^) assuming logarithmic (left) or linear (middle) changes in baseline dopamine induced by dopamine transients (logΔDA, linΔDA, respectively). The absolute magnitude of the slopes differs depending on the assumption made about the changes in baseline dopamine (logarithmic vs linear) but the asymmetric scaling factor presents only a small shift in the curve as a function of baseline dopamine (right). The qualitative aspects of the model (i.e., non-monotonic relationship of the asymmetric scaling factor with baseline dopamine) is preserved regardless on this assumption.

**Extended Data Fig. 10.**
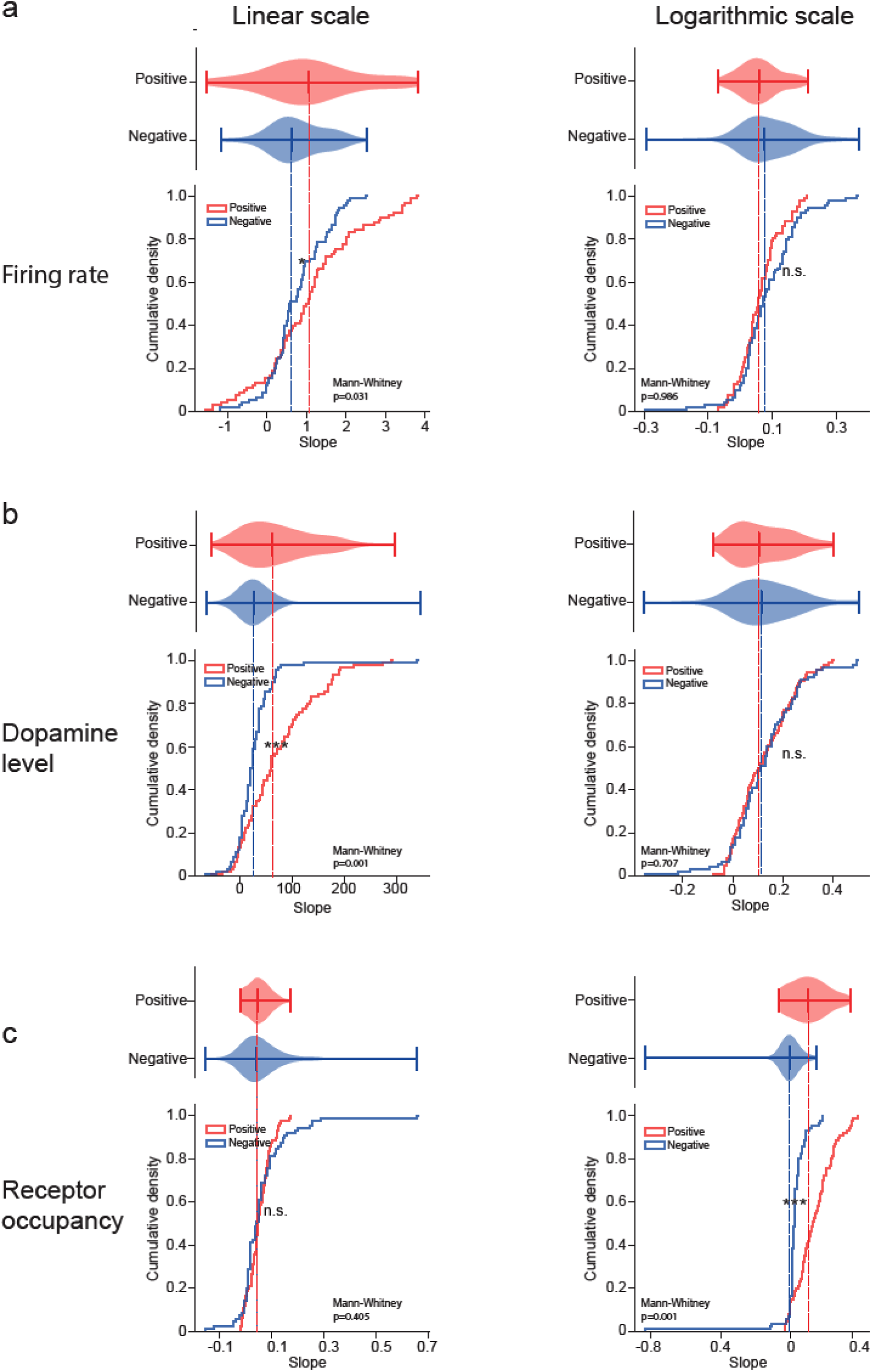
Change in dopamine firing rates, dopamine concentration and receptor occupancy as a function of RPEs in the linear scale or logarithmic scale. **a**.Distributions of the slopes of the change in firing rate derived as a function of RPEs in the positive and negative domains, computed in the linear (left) or logarithmic (right) scale, calculated at a single neuron level. The slopes are asymmetric if considered in the linear scale, with the negative transients presenting a shallower slope than the positive ones. The slopes are symmetric if considered in the logarithmic scale. **b**.Distributions of the slopes of the change in dopamine levels derived from the biophysical model as a function of RPEs in the positive and negative domains, computed in the linear (left) or logarithmic (right) scale, calculated at a single neuron level. The slopes are again asymmetric if considered in the linear scale but symmetric if considered in the logarithmic scale. **c**. Slope of the change in receptor occupancy derived from the biophysical model for a given RPE in the positive and negative domains, computed in the linear (left) or logarithmic (right) scale, calculated at a single neuron level. The slopes are symmetric if considered in the linear scale but asymmetric if considered in the logarithmic scale.

**Extended Data Fig. 11.**
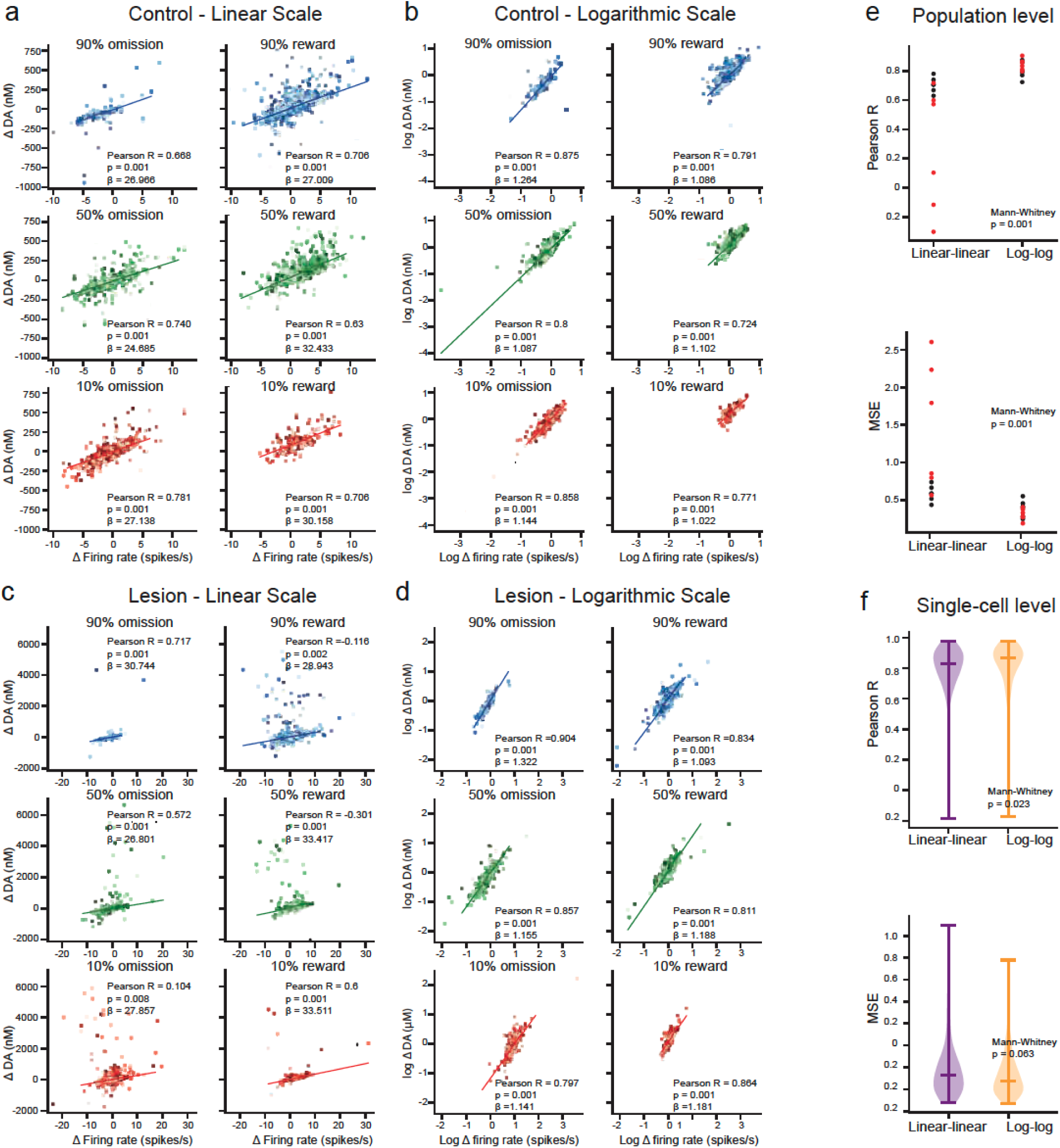
Relationship between changes in firing rates and changes in dopamine concentration derived from the biophysical model. **a**.Linear fits to the relationship between changes in firing rates and changes in dopamine concentration evoked by the TD error at outcome in the linear scale for the control group. The fits are done separately for each trial type. **b**. Same as a, but fits are done in the logarithmic scale. **c**. Same as a, but fits are done for the lesion group. **d**. Same as c, but fits are done in the logarithmic scale. **e**. Distribution of the Pearson correlation coefficients (top) and means squared error (MSE, bottom) between the predicted change in dopamine concentration by the linear regression and the ground truth derived from the biophysical model. The coefficients are derived from the fits in figures a-e, done by pooling all trials for each trial type (each point each trial type, with black for control and red for lesion group). There was a significant increase in the Pearson correlation coefficient and a near significant decrease in the MSE if the changes are considered to happen in the logarithmic scale. **f**. Same as E, but the linear regression fits are done for each neuron separately by pooling all trials. There was a significant increase in the single-cell distribution of Pearson correlation coefficients (top) and a significant decrease in the MSE distribution (bottom) if the changes are considered to happen in the logarithmic scale.

**Extended Data Fig. 12.**
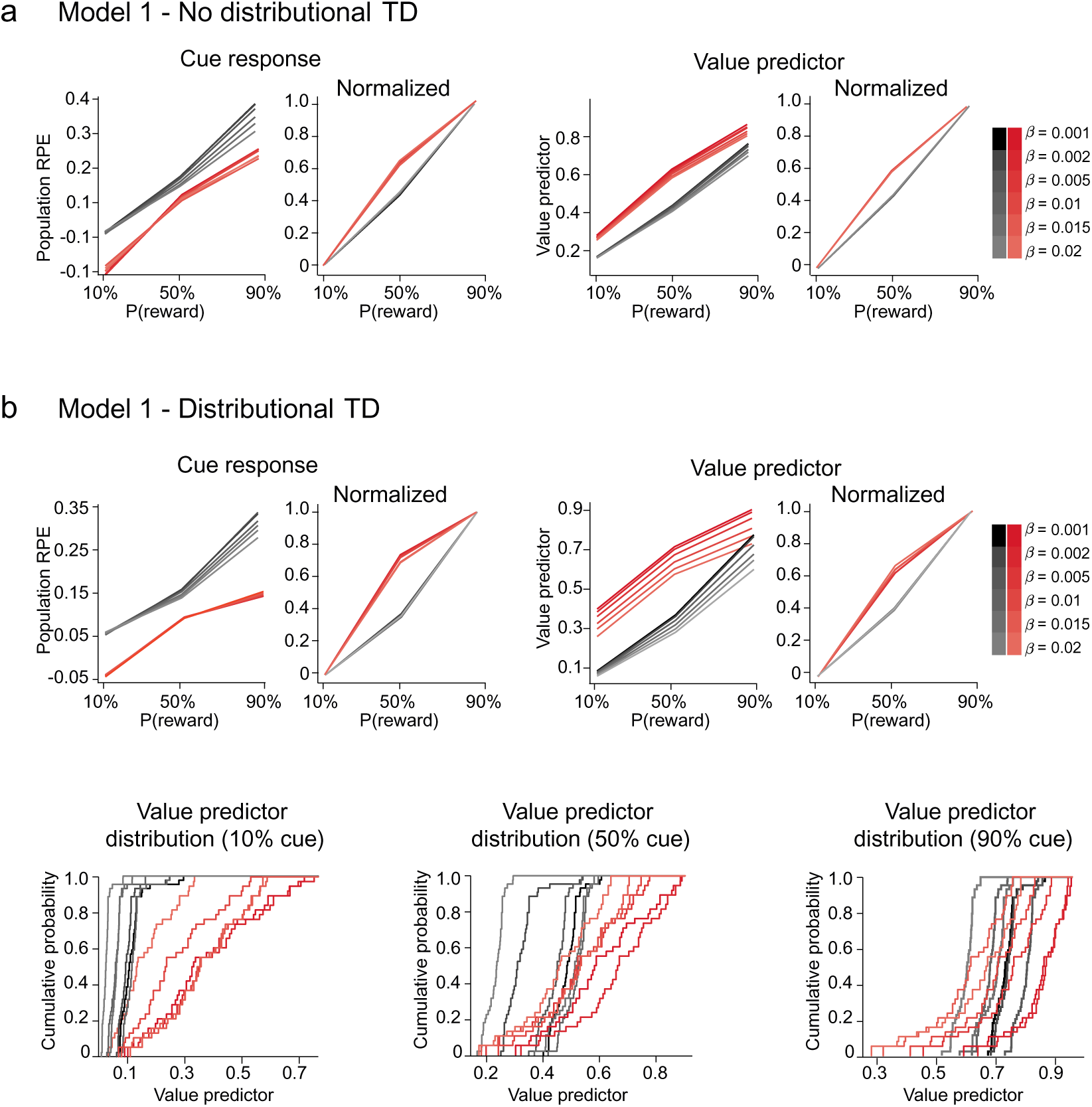
Model 1 captures signatures of the data irrespective of the choice of the decay factor and is compatible with distributional RL. **a**. Model 1 with standard TD learning. Simulations were run using the receptors sensitivities from the biophysical models and data-derived asymmetric scaling factors (see Methods 3.3). The model’s predictions capture the signatures in cue-evoked dopamine responses (left) and value predictions (right) irrespective of the choice of the decay factor (β). **b**. Model 1 within the distributional RL framework (see Methods 3.3). The model’s predictions also capture the signatures in cue-evoked dopamine responses (left) and value predictions (right) irrespective of the choice of the decay factor (β). Bottom row shows the distribution of value predictors for each reward-predictive cue.

**Extended Data Fig. 13.**
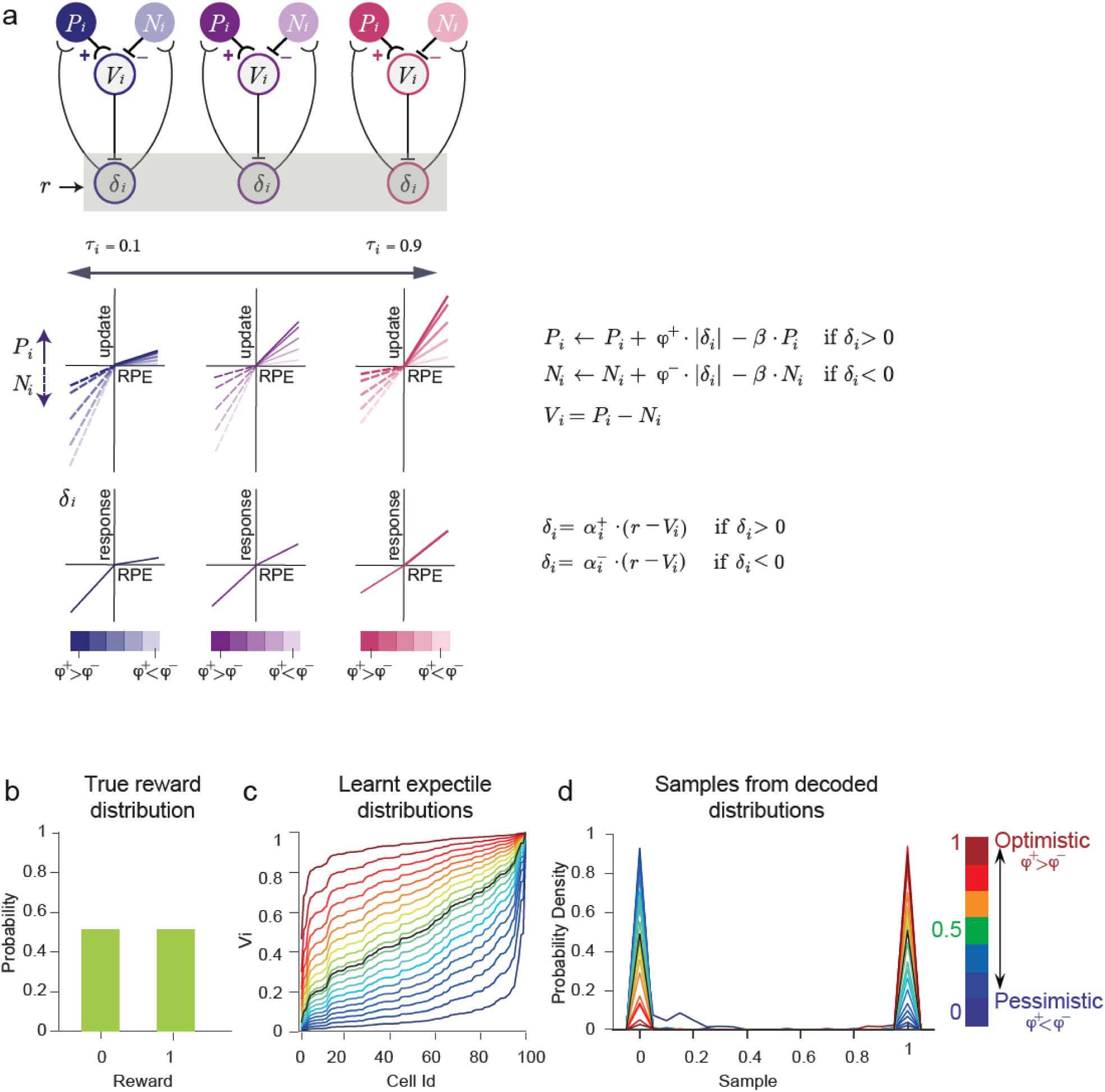
Distributional reinforcement learning with D1 and D2 populations (Model 1). Schematic of the distributional RL model with D1 and D2 populations. The schematic represents three different value predictors (pessimistic, neutral and optimistic from left to right) with their respective *P* and *N* neurons. The level of optimism of each individual value predictor is determined by the scaling factors of the individual dopamine RPE-evoked responses (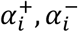 represented by the color in the colormap from purple to pink) and allows the model to encode information about the distribution of rewards (bottom). The global level of ‘optimism’ or ‘pessimism’ of the agent is given by the re-scaling of the RPEs by the P and N receptors sensitivities in the model (*ϕ*^+^, *ϕ* ^−^, represented with the color saturation). **b**. Example of a Bernoulli distribution, equivalent to the reward distribution predicted by the 50% cue. **c**. Distribution of expectiles learnt by the distributional RL model with D1 and D2 population for the reward distribution in b. The expectiles are sorted based on the asymmetric scaling factor of each individual dopamine neuron. Colormap represents the level of optimistic or pessimism of each agent. **d**. Samples from the decoded distributions for the set of expectiles in c. The probability density is bimodal, in accordance with the distribution in b. As the agents goes from pessimism to optimism, the probability density modes change in their relative magnitude.

## References

1. Brown, V. M., Zhu L., Solway A., Wang M., McCurry K., King-Casas B. & Chiu P. Reinforcement Learning Disruptions in Individuals With Depression and Sensitivity to Symptom Change Following Cognitive Behavioral Therapy. JAMA Psychiatry 78, 1113–1122 (2021).

2. Groman, S. M., Thompson, S. L., Lee, D. & Taylor, J. R. Reinforcement learning detuned in addiction: integrative and translational approaches. Trends Neurosci. 45, 96–105 (2022).

3. Ligneul, R., Sescousse, G., Barbalat, G., Domenech, P. & Dreher, J.-C. Shifted risk preferences in pathological gambling. Psychol. Med. 43, 1059–1068 (2013).

4. Mason, L., O’Sullivan, N., Bentall, R. P. & El-Deredy, W. Better than I thought: positive evaluation bias in hypomania. PLoS One 7, e47754 (2012).

5. Pizzagalli, D. A., Iosifescu, D., Hallett, L. A., Ratner, K. G. & Fava, M. Reduced hedonic capacity in major depressive disorder: evidence from a probabilistic reward task. J. Psychiatr. Res. 43, 76–87 (2008).

6. Verdejo-Garcia, A., Chong, T. T.-J., Stout, J. C., Yücel, M. & London, E. D. Stages of dysfunctional decision-making in addiction. Pharmacol. Biochem. Behav. 164, 99–105 (2018).

7. Lim, T. V., Cardinal, R. N., Bullmore, E. T., Robbins, T. W. & Ersche, K. D. Impaired Learning From Negative Feedback in Stimulant Use Disorder: Dopaminergic Modulation. Int. J. Neuropsychopharmacol. 24, 867–878 (2021).

8. Schönfelder, S., Langer, J., Schneider, E. E. & Wessa, M. Mania risk is characterized by an aberrant optimistic update bias for positive life events. J. Affect. Disord. 218, 313–321 (2017).

9. Dayan, P. & Daw, N. D. Decision theory, reinforcement learning, and the brain. Cogn. Affect. Behav. Neurosci. 8, 429–453 (2008).

10. Katahira, K. The relation between reinforcement learning parameters and the influence of reinforcement history on choice behavior. J. Math. Psychol. 66, 59–69 (2015).

11. Sutton, R. S. & Barto, A. G. Reinforcement Learning: An Introduction. (Bradford Books, 2018).

12. Maia, T. V. & Frank, M. J. From reinforcement learning models to psychiatric and neurological disorders. Nat. Neurosci. 14, 154–162 (2011).

13. Eldar, E., Rutledge, R. B., Dolan, R. J. & Niv, Y. Mood as Representation of Momentum. Trends Cogn. Sci. 20, 15–24 (2016).

14. Rutledge, R. B., Skandali, N., Dayan, P. & Dolan, R. J. A computational and neural model of momentary subjective well-being. Proc. Natl. Acad. Sci. U. S. A. 111, 12252–12257 (2014).

15. Floresco, S. B., West, A. R., Ash, B., Moore, H. & Grace, A. A. Afferent modulation of dopamine neuron firing differentially regulates tonic and phasic dopamine transmission. Nat. Neurosci. 6, 968–973 (2003).

16. Wang, Y., Toyoshima, O., Kunimatsu, J., Yamada, H. & Matsumoto, M. Tonic firing mode of midbrain dopamine neurons continuously tracks reward values changing moment-by-moment. Elife 10, (2021).

17. Schultz, W., Dayan, P. & Montague, P. R. A neural substrate of prediction and reward. Science 275, 1593–1599 (1997).

18. Eshel, N., Tian, J., Bukwich, M. & Uchida, N. Dopamine neurons share common response function for reward prediction error. Nat. Neurosci. 19, 479–486 (2016).

19. Steinberg, E. Keiflin, R., Boivin, J., Witten I., Deisseroth K. & Janak P. A causal link between prediction errors, dopamine neurons and learning. Nat. Neurosci. 16, 966–973 (2013).

20. Waelti, P., Dickinson, A. & Schultz, W. Dopamine responses comply with basic assumptions of formal learning theory. Nature 412, 43–48 (2001).

21. Korn, C. W., Sharot, T., Walter, H., Heekeren, H. R. & Dolan, R. J. Depression is related to an absence of optimistically biased belief updating about future life events. Psychol. Med. 44, 579–592 (2014).

22. Rutledge, R. B., Lazzaro, S. C., Lau, B., Myers, C. E., Gluck, M. A., & Glimcher, P. W. Dopaminergic drugs modulate learning rates and perseveration in Parkinson’s patients in a dynamic foraging task. J. Neurosci. 29, 15104–15114 (2009).

23. Frank, M. J., Seeberger, L. C. & O’Reilly, R. C. By carrot or by stick: Cognitive reinforcement learning in Parkinsonism. Science 306, 1940–1943 (2004).

24. Cools, R., Frank M.J., Gibbs S., Miyakawa A., Jagust W. & D’Esposito M. Striatal dopamine predicts outcome-specific reversal learning and its sensitivity to dopaminergic drug administration. J. Neurosci. 29, 1538–1543 (2009).

25. Timmer, M. H. M., Sescousse, G., van der Schaaf, M. E., Esselink, R. A. J. & Cools, R. Reward learning deficits in Parkinson’s disease depend on depression. Psychol. Med. 47, 2302–2311 (2017).

26. Gradin, V. B., Kumar, P., Waiter, G., Ahearn, T., Stickle, C., Milders, M., Reid, I., Hall, J., & Steele, J. D. Expected value and prediction error abnormalities in depression and schizophrenia. Brain 134, 1751–1764 (2011).

27. Kumar, P., Waiter, G., Ahearn, T., Milders, M., Reid, I., & Steele, J. D. Abnormal temporal difference reward-learning signals in major depression. Brain 131, 2084–2093 (2008).

28. Pizzagalli, D. A., Holmes, A. J., Dillon, D. G., Goetz, E. L., Birk, J. L., Bogdan, R., Dougherty, D. D., Iosifescu, D. V., Rauch, S. L., & Fava, M. Reduced caudate and nucleus accumbens response to rewards in unmedicated individuals with major depressive disorder. Am. J. Psychiatry 166, 702–710 (2009).

29. Robinson, O. J., Cools, R., Carlisi, C. O., Sahakian, B. J. & Drevets, W. C. Ventral striatum response during reward and punishment reversal learning in unmedicated major depressive disorder. Am. J. Psychiatry 169, 152–159 (2012).

30. Collins, A. G. E. & Frank, M. J. Opponent actor learning (OpAL): modeling interactive effects of striatal dopamine on reinforcement learning and choice incentive. Psychol. Rev. 121, 337–366 (2014).

31. Frank, M. J. Dynamic Dopamine Modulation in the Basal Ganglia: A Neurocomputational Account of Cognitive Decits in Medicated and Non-medicated Parkinsonism. J. Cogn. Neuroci. 17, 51–72 (2005).

32. Mikhael, J. G. & Bogacz, R. Learning Reward Uncertainty in the Basal Ganglia. PLoS Comput. Biol. 12, 1–28 (2016).

33. Yagishita, S., Hayashi-Takagi, A., Ellis-Davies, G.C.R., Urakubo, H., Ishii, S. & Kasai, H. A critical time window for dopamine actions on the structural plasticity of dendritic spines. Science 345, 1616–1620 (2014).

34. Iino, Y. Sawada, T., Yamaguchi, K. et al. Dopamine D2 receptors in discrimination learning and spine enlargement. Nature 579, 555–560 (2020).

35. Lee, S. J., Lodder, B., Chen, Y., Patriarchi, T., Tian, L & Sabatini, B. Cell-type-specific asynchronous modulation of PKA by dopamine in learning. Nature 590, 451–456 (2021).

36. Dabney, W., Kurth-Nelson, Z., Uchida, N., Starkweather, C., Hassabis, D., Munos, R. & Botvinick, M. A distributional code for value in dopamine-based reinforcement learning. Nature 1, (2019).

37. Bellemare, M. G., Dabney, W. & Munos, R. A distributional perspective on reinforcement learning. 34th International Conference on Machine Learning, ICML 2017 1, 693–711 (2017).

38. Lowet, A. S., Zheng, Q., Matias, S., Drugowitsch, J. & Uchida, N. Distributional Reinforcement Learning in the Brain. Trends Neurosci. 43, 980–997 (2020).

39. Rowland, M., Dadashi, R., Kumar, S., Munos, R., Bellemare, M. G. & Dabney, W. Statistics and samples in distributional reinforcement learning. 36th International Conference on Machine Learning, ICML 2019 6, 9727–9750 (2019).

40. Mihatsch, O. & Neuneier, R. Risk-sensitive reinforcement learning. Mach. Learn. 49, 267–290 (2002).

41. Bellemare, M. G. & Dabney, W. Distributional reinforcement learning. (MIT Press, 2023).

42. Jones, M. C. Expectiles and M-quantiles are quantiles. Stat. Probab. Lett. 20, 149–153 (1994).

43. Houk, J., Davis, J., & Beiser, D. Models of information processing in the basal ganglia. (Bradford Books, 2019).

44. Gerfen, C. The neostriatal mosaic: Multiple levels of compartmental organization in the basal ganglia. Annu. Rev. Neurosci. 15, 285–320 (1992).

45. Smith, Y., Bevan, M. D., Shink, E. & Bolam, J. P. Microcircuitry of the direct and indirect pathways of the basal ganglia. Neuroscience 86, 353–387 (1998).

46. Kravitz, A. V., Tye, L. D. & Kreitzer, A. C. Distinct roles for direct and indirect pathway striatal neurons in reinforcement. Nat. Neurosci. 15, 816–818 (2012).

47. Reynolds, J. N. J., Hyland, B. I. & Wickens, J. R. A cellular mechanism of reward-related learning. Nature 413, 67–70 (2001).

48. Gerfen, C. R. & Surmeier, D. J. Modulation of striatal projection systems by dopamine. Annu. Rev. Neurosci. 34, 441–466 (2011).

49. Richfield, E. K., Penney, J. B. & Young, A. B. Anatomical and affinity state comparisons between dopamine D1 and D2 receptors in the rat central nervous system. Neuroscience 30, 767–777 (1989).

50. Rice, M. E. & Cragg, S. J. Dopamine spillover after quantal release: Rethinking dopamine transmission in the nigrostriatal pathway. Brain Res. Rev. 58, 303–313 (2008).

51. Gonon, F. G. & Buda, M. J. Regulation of dopamine release by impulse flow and by autoreceptors as studied by in vivo voltammetry in the rat striatum. Neuroscience 14, 765–774 (1985).

52. Dodson, P. D., Dreyer, J. K., Jennings, K. A., Syed, E. C. J., Wade-Martins, R., Cragg, S. J., Bolam, J. P. & Magill, P. J. Representation of spontaneous movement by dopaminergic neurons is cell-type selective and disrupted in parkinsonism. Proc. Natl. Acad. Sci. U. S. A. 113, E2180–E2188 (2016).

53. Marcott, P. F., Mamaligas, A. A. & Ford, C. P. Phasic dopamine release drives rapid activation of striatal D2-receptors. Neuron 84, 164–176 (2014).

54. Jaskir, A. & Frank, M. J. On the normative advantages of dopamine and striatal opponency for learning and choice. Elife (2023).

55. Tian, J. & Uchida, N. Habenula Lesions Reveal that Multiple Mechanisms Underlie Dopamine Prediction Errors. Neuron 87, 1304–1316 (2015).

56. Cui, Y., Yang, Y., Ni, Z., Dong, Y., Cai, G., Foncelle, A., Ma, S., Sang, K., Tang, S., Li, Y., Shen, Y., Berry, H., Wu, S. & Hu, H. Astroglial Kir4.1 in the lateral habenula drives neuronal bursts in depression. Nature 554, 323–327 (2018).

57. Li, B., Piriz, J., Mirrione, M., Chung, C., D. Proulx, C., Schulz, D., Henn, F. & Malinow, R. Synaptic potentiation onto habenula neurons in the learned helplessness model of depression. Nature 470, 535–541 (2011).

58. Yang, Y., Cui, Y., Sang, K., Dong, Y., Ni, Z., Ma, S. & Hu, H. Ketamine blocks bursting in the lateral habenula to rapidly relieve depression. Nature 554, 317–322 (2018).

59. Dreyer, J. K., Herrik, K. F., Berg, R. W. & Hounsgaard, J. D. Influence of phasic and tonic dopamine release on receptor activation. J. of Neurosci. 30, 14273–14283 (2010).

60. Mikhael, J. G., Lai, L. & Gershman, S. J. Rational inattention and tonic dopamine. PLoS Comput. Biol. 17 e1008659 (2021).

61. Vingerhoets, F. J., Snow, B. J., Schulzer, M., Morrison, S., Ruth, T. J., Holden, J. E., Cooper, S. & Calne, D. B. Reproducibility of fluorine-18-6-fluorodopa positron emission tomography in normal human subjects. J. Nucl. Med. 35, 18–24 (1994).

62. Cools, R., Gibbs, S. E., Miyakawa, A., Jagust, W. & D’Esposito, M. Working memory capacity predicts dopamine synthesis capacity in the human striatum. J. Neurosci. 28, 1208–1212 (2008).

63. Hoffmann, I. S. & Cubeddu, L. X. Differential effects of bromocriptine on dopamine and acetylcholine release modulatory receptors. J. Neurochem. 42, 278–282 (1984).

64. Tissari, A. H., Rossetti, Z. L., Meloni, M., Frau, M. I. & Gessa, G. L. Autoreceptors mediate the inhibition of dopamine synthesis by bromocriptine and lisuride in rats. Eur. J. Pharmacol. 91, 463–468 (1983).

65. Lieberman, A. Depression in Parkinson’s disease -- a review. Acta Neurol. Scand. 113, 1–8 (2006).

66. Leentjens, A. F. G., Van den Akker, M., Metsemakers, J. F. M., Lousberg, R. & Verhey, F. R. J. Higher incidence of depression preceding the onset of Parkinson’s disease: a register study. Mov. Disord. 18, 414–418 (2003).

67. Nilsson, F. M., Kessing, L. V., Sørensen, T. M., Andersen, P. K. & Bolwig, T. G. Major depressive disorder in Parkinson’s disease: a register-based study. Acta Psychiatr. Scand. 106, 202–211 (2002).

68. Remy, P., Doder, M., Lees, A., Turjanski, N. & Brooks, D. Depression in Parkinson’s disease: loss of dopamine and noradrenaline innervation in the limbic system. Brain 128, 1314–1322 (2005).

69. Weintraub, D., Newberg, A. B., Cary, M. S., Siderowf, A. D., Moberg, P. J., Kleiner-Fisman, G., Duda, J. E., Stern, M. B., Mozley, D. & Katz, I. R. Striatal dopamine transporter imaging correlates with anxiety and depression symptoms in Parkinson’s disease. J. Nucl. Med. 46, 227–232 (2005).

70. Kish, S. J., Shannak, K. & Hornykiewicz, O. Uneven pattern of dopamine loss in the striatum of patients with idiopathic Parkinson’s disease. Pathophysiologic and clinical implications. N. Engl. J. Med. 318, 876–880 (1988).

71. Timmer, M. H. M., Sescousse, G., Esselink, R. A. J., Piray, P. & Cools, R. Mechanisms Underlying Dopamine-Induced Risky Choice in Parkinson’s Disease With and Without Depression (History). Comput Psychiatr 2, 11–27 (2018).

72. Hikida, T., Kimura, K., Wada, N., Funabiki, K. & Nakanishi Shigetada, S. Distinct Roles of Synaptic Transmission in Direct and Indirect Striatal Pathways to Reward and Aversive Behavior. Neuron 66, 896–907 (2010).

73. Hikida, T., Yawata, S., Yamaguchi, T., Danjo, T., Sasaoka, T., Wang, Y. & Nakanishi S. Pathway-specific modulation of nucleus accumbens in reward and aversive behavior via selective transmitter receptors. Proc. Natl. Acad. Sci. U. S. A. 110, 342–347 (2013).

74. Danjo, T., Yoshimi, K., Funabiki, K., Yawata, S. & Nakanishi, S. Aversive behavior induced by optogenetic inactivation of ventral tegmental area dopamine neurons is mediated by dopamine D2 receptors in the nucleus accumbens. Proc. Natl. Acad. Sci. U. S. A. 111, 6455–6460 (2014).

75. Yamaguchi, T., Goto, A., Nakahara, I., Yawata, S., Hikida, T., Matsuda, M., Funabiki, K & Nakanishi, S. Role of PKA signaling in D2 receptor-expressing neurons in the core of the nucleus accumbens in aversive learning. Proc. Natl. Acad. Sci. U. S. A. 112, 11383–11388 (2015).

76. Bayer, H. M. & Glimcher, P. W. Midbrain dopamine neurons encode a quantitative reward prediction error signal. Neuron 47, 129–141 (2005).

77. Hart, A. S., Rutledge, R. B., Glimcher, P. W. & Phillips, P. E. M. Phasic dopamine release in the rat nucleus accumbens symmetrically encodes a reward prediction error term. J. Neurosci. 34, 698–704 (2014).

78. Grace, A. A. Dysregulation of the dopamine system in the pathophysiology of schizophrenia and depression. Nat. Rev. Neurosci. 17, 524–532 (2016).

79. Anstrom, K. K., Miczek, K. A. & Budygin, E. A. Increased phasic dopamine signaling in the mesolimbic pathway during social defeat in rats. Neuroscience 161, 3–12 (2009).

80. Razzoli, M., Andreoli, M., Michielin, F., Quarta, D. & Sokal, D. M. Increased phasic activity of VTA dopamine neurons in mice 3 weeks after repeated social defeat. Behav. Brain Res. 218, 253–257 (2011).

81. Markovic, T., Pederson, C. E., Massaly, N., Vachez, Y., Ruyle, B., Murphy, C. A., Abiraman, K., Hoon Shin, J., Garcia, J. H., Jean Yoon, H., Alvarez, V. A., Bruchas, M. R., Creed, M. C. & Moron, J. A. Pain induces adaptations in ventral tegmental area dopamine neurons to drive anhedonia-like behavior. Nat. Neurosci. 24, 1601–1613 (2021).

82. Guo, Z., Li, S., Wu, J., Zhu, X. & Zhang, Y. Maternal deprivation increased vulnerability to depression in adult rats through DRD2 promoter methylation in the ventral tegmental area. Front. Psychiatry 13, 827667 (2022).

83. Peng, B., Hu, Q., Liu, J., Guo, S., Borgland, S. L.& Liu, S. Corticosterone attenuates reward-seeking behavior and increases anxiety via D2 receptor signaling in ventral tegmental area dopamine neurons. J. Neurosci. 41, 1566–1581 (2021).

84. Tye, K. M., Mirzabekov, J. J., Warden, M. R., Ferenczi, E. A., Tsai, H-C., Finkelstein, J., Kim, S-Y., Adhikari, A., Thompson, K. R., Andalman A. S., Gunaydin, L. A., Witten I. & Deisseroth K. Dopamine neurons modulate neural encoding and expression of depression-related behaviour. Nature 493, 537–541 (2013).

85. Baek, K., Kwon, J., Chae, J. H., Chung, Y. A., Kralik, J. D., Min, J. A., Huh, H., Choi, K. M., Jang, K. I., Lee, N. B., Kim, S., Peterson, B. S., & Jeong, J. Heightened aversion to risk and loss in depressed patients with a suicide attempt history. Sci. Rep. 7, 11228 (2017).

86. Smoski, M. J., Lynch, T. R., Rosenthal, M. Z., Cheavens, J. S., Chapman, A. L., & Krishnan R. R. Decision-making and risk aversion among depressive adults. J. Behav. Ther. Exp. Psychiatry 39, 567–576 (2008).

87. van Holst, R. J., Sescousse, G., Janssen, L. K., Janssen, M., Berry, A. S., Jagust, W. J., & Cools, R. Increased Striatal Dopamine Synthesis Capacity in Gambling Addiction. Biol. Psychiatry 83, 1036–1043 (2018).

88. Cools, R., Altamirano, L. & D’Esposito, M. Reversal learning in Parkinson’s disease depends on medication status and outcome valence. Neuropsychologia 44, 1663–1673 (2006).

89. Cools, R., Barker, R. A., Sahakian, B. J. & Robbins, T. W. Enhanced or impaired cognitive function in Parkinson’s disease as a function of dopaminergic medication and task demands. Cereb. Cortex 11, 1136–1143 (2001).

90. Cools, R., Barker, R. A., Sahakian, B. J. & Robbins, T. W. L-Dopa medication remediates cognitive inflexibility, but increases impulsivity in patients with Parkinson’s disease. Neuropsychologia 41, 1431–1441 (2003).

91. Swainson, R., Rogers, R. D., Sahakian, B. J., Summers, B. A., Polkey, C. E., & Robbins, T. W. Probabilistic learning and reversal deficits in patients with Parkinson’s disease or frontal or temporal lobe lesions: possible adverse effects of dopaminergic medication. Neuropsychologia 38, 596–612 (2000).

92. Cox, S. M., Frank, M. J., Larcher, K., Fellows, L. K., Clark, C. A., Leyton, M., & Dagher, A. Striatal D1 and D2 signaling differentially predict learning from positive and negative outcomes. Neuroimage 109, 95–101 (2015).

93. Savitz, J. B. & Drevets, W. C. Neuroreceptor imaging in depression. Neurobiol. Dis. 52, 49–65 (2013).

94. Rescorla, R. A. & Wagner, A. R. A theory of Pavlovian conditioning: Variations in the effectiveness of reinforcement and nonreinforcement, Classical Conditioning II. 64–99 (1972).

95. Kim, H. R., Malik, A. N., Mikhael, J. G., Bech, P., Tsutsui-Kimura, I., Sun, F., Zhang, Y., Li, Y., Watabe-Uchida, M., Gershman, S. J. & Uchida, N. A unified framework for dopamine signals across timescales. Cell 183, 1600–1616.e25 (2020).

96. Bertsekas, D. P. & Tsitsiklis, J. Neuro-Dynamic Programming. (Athena Scientific, 1996).

97. Mierau, J., Schneider, F. J., Ensinger, H. A., Chio, C. L., Lajiness, M. E., & Huff, R. M. Pramipexole binding and activation of cloned and expressed dopamine D2, D3 and D4 receptors. Eur. J of Pharmac. 290, 29–36 (1995).

98. Cohen, J. Y., Haesler, S., Vong, L., Lowell, B. B. & Uchida, N. Neuron-type-specific signals for reward and punishment in the ventral tegmental area. Nature 482, 85–88 (2012).

99. Kvitsiani, D., Ranade, S., Hangya, B., Taniguchi, H., Huang, J. Z. & Kepec, A. Distinct behavioural and network correlates of two interneuron types in prefrontal cortex. Nature 498, 363–366 (2013).

100. Stauffer, W. R., Lak, A. & Schultz, W. Dopamine reward prediction error responses reflect marginal utility. Curr. Biol. 24, 2491–2500 (2014).

